# Bezafibrate treatment rescues neurodevelopmental and neurodegenerative defects in 3D cortical organoid model of MAPT frontotemporal dementia

**DOI:** 10.1101/2024.08.02.606317

**Authors:** Federica Cordella, Lorenza Mautone, Lucrezia Tondo, Debora Salerno, Silvia Ghirga, Erika Parente, Maria Anele Romeo, Mara Cirone, Paola Bezzi, Silvia Di Angelantonio

## Abstract

**INTRODUCTION:** The intronic MAPT mutation IVS10+16 is linked to familiar frontotemporal dementia, causing hyperphosphorylation and accumulation of tau protein, resulting in synaptic and neuronal loss and neuroinflammation in patients. This mutation disrupts MAPT gene splicing, increasing exon 10 inclusion and leading to an imbalance of 3R and 4R Tau isoforms.

**METHODS:** We generated patterned cortical organoids from isogenic control and mutant human iPSC lines. Nanostring gene expression analysis immunofluorescence and calcium imaging recordings were used to study the impact of the MAPT IVS10+16 mutation on neuronal development and function.

**RESULTS:** Tau mutant cortical organoids showed altered mitochondrial function and gene expression related to neuronal development, with synaptic markers and neuronal activity reduction. Bezafibrate treatment, which restored mitochondrial content, rescued synaptic functionality and tau physiology.

**DISCUSSION:** These findings suggest that targeting mitochondrial function with bezafibrate could potentially reverse tau-induced neurodevelopmental deficits, highlighting its therapeutic potential for tauopathies like FTD.

**HIGLIGHTS:**

- The IVS 10+16 MAPT mutation significantly disrupts cortical differentiation and synaptic maturation, evidenced by downregulated genes associated with synapses and neuronal development.
- Tau-mutant cortical organoids exhibit mitochondrial dysfunction, with fewer and smaller mitochondria alongside with tau hyperphosphorylation and aggregation, which further contribute to neuronal damage and disease progression.
- Treatment with bezafibrate effectively normalizes mitochondrial parameters, enhances neuronal integrity and synaptic maturation, and restores network functionality, showcasing its promise as a therapeutic strategy for tauopathies.
- The 3D in vitro disease model used in this study proves valuable for studying tauopathies and testing new drugs, effectively mimicking key aspects of tau-related neurodegeneration.

**RESEARCH IN CONTEXT:**

1. **Systematic Review:** We searched PubMed, Google Scholar, and Web of Science for studies on the MAPT IVS10+16 mutation’s impact on tauopathies, focusing on neuronal development, synaptic function, and mitochondrial involvement. Key terms included “MAPT IVS10+16 mutation,” “tauopathy,” “neuronal development,” “synaptic function,” and “mitochondrial function.”
2. **Interpretation:** Our findings reveal that the MAPT IVS10+16 mutation disrupts mitochondrial function altering gene expression related to neuronal development, synaptic structures, impairing neuronal and glial maturation. Bezafibrate treatment restored mitochondrial content, synaptic functionality, and tau physiology in mutant-derived cortical organoids, suggesting it as a potential therapeutic strategy for tauopathies.
3. **Future Directions:** Future research should investigate the molecular mechanisms underlying the bezafibrate’s therapeutic effect and its long-term efficacy and safety in vivo, in humanized mouse models. Additionally, the possibility to combine bezafibrate with other therapeutic agents used to treat tauopathies will be worth to assess.

## 1 BACKGROUND

### The introduction cannot exceed 650 words (ora 658)

Frontotemporal dementia (FTD) is a debilitating neurodegenerative disorder characterized by the degeneration of neuronal cells within the frontal and temporal lobes of the brain, primarily affecting personality, behavior, language, and executive functioning (1) (Antonioni et al., 2023). Considering the diverse spectrum of pathological mechanisms contributing to FTD, abnormal changes in tau protein represent a critical avenue of investigation. Tau, encoded by the MAPT gene, is a multitasking protein, and among its functions interacts with tubulin, enhancing its polymerization into microtubules, and regulates their assembly, dynamics, and spatial organization (2) (3) (Chang et al., 2021; Barbier et al., 2019). The alternative splicing of tau pre-mRNA is developmentally regulated, resulting in the production of six isoforms that differ in containing three (3R) or four (4R) microtubule-binding regions in the carboxyl terminal and one (1N), two (2N), or zero (0N) amino terminal inserts (4) (Leveille et al., 2021).

Among tau mutations linked to FTD, the MAPT intronic IVS 10+16 mutation, located within the MAPT gene on chromosome 17, has garnered significant attention due to its association with familial a form of the disease (5) (Fujioka et al., 2011). This mutation enhances the probability of the inclusion of exon 10 within tau transcripts, resulting in the production of tau isoforms with altered microtubule-binding properties and a propensity for pathological aggregation (6) (Verheyen et al., 2019), with increase in 4R tau isoform splicing, leading to significant alterations in neuronal functionality (6) (7) (Verheyen et al., 2019; Kopach et al.,2021).

Among the role of tau in shaping neuronal homeostasis, its interaction with mitochondria has been highlighted in physiological and pathological conditions as FTD (2)(8)(9)(10)(11)(Chang et al., 2021; Declan et al., 2022; Tracy et al., 2022; Szabo et al., 2020, Quantanilla et al., 2018). Several studies suggest a potential feedback loop where tau dysregulation impairs mitochondrial homeostasis, and mitochondrial dysfunction exacerbates tau pathology by increasing oxidative stress and hindering the cellular clearance of abnormal tau aggregates. This mitochondria-tau crosstalk may play a crucial role in triggering and worsening diseases such as FTD (12) (Cheng et al., 2018).

Given the early onset and poor prognosis of FTD, there is an urgent need to develop innovative in vitro models that more accurately recapitulate the disease’s key features. These models are essential for dissecting molecular pathological mechanisms, identifying novel therapeutic targets, and testing novel and repurposed drug candidates. Over the years, transgenic animal models have been instrumental in elucidating FTD-associated genetic mutations and the resultant functional changes that characterize this pathology. Although mouse models can replicate key pathological hallmarks such as neurofibrillary tangle deposition, they do not fully reproduce the pathological phenotypes observed in humans, particularly in terms of neuronal loss and neurodegeneration (13)(14) (Cordella et al., 2022a; Brighi et al., 2020). Therefore, induced pluripotent stem cell (iPSC)-derived cortical organoids, characterized by 3D structure, represent a more suitable in vitro tool for studying pathological events that occur in vivo (15) (D’Antoni et al., 2023). Notably, the brain organoid model recapitulates the early stages of human cortical development, allowing the evaluation of phenotypic perturbations in neuro-glia development during critical developmental periods that are otherwise challenging to investigate in humans, thus giving the opportunity to underscore neurodevelopmental deficits associated to typical neurodegenerative disorders (16) (Wegiel et al., 2022).

In this study, we developed a human iPSC-derived cortical organoid model based on control and isogenic human iPSC harboring the intronic MAPT IVS 10+16 mutation. Our findings, by molecular barcoding approach important to identify molecular signatures in neurological diseases and degenerative disorders, indicate that the intronic MAPT IVS10+16 mutation leads to delayed and impaired cortical development due to reduced mitochondrial functions, affecting the maturation and function of both neuronal and glial cells. This mutation also results in the increased expression of 4R tau isoforms and in the accumulation of pathological tau forms. Additionally, we investigated the potential effects of Bezafibrate (BZ), agonist of the master regulator of mitochondrial biogenesis PGC1α in our in vitro disease model. Our results show that BZ treatment restores neuronal cytoskeletal morphology and mitochondrial function, promotes neuronal maturation, and rescues synaptic activity, reducing tau hyperphosphorylation, demonstrating a promising therapeutic potential for tauopathies.

## 2 METHODS

### 2.1 Maintenance of human iPSC

iPSC0028 (Ebisc – Sigma) and isogenic SIGI001-A-13 (IVS10+16) (Ebisc-Sigma) hiPSC lines were cultured in mTeSR plus medium (STEMCELL technologies) hESC-qualified Matrigel (CORNING) functionalized plates. The culture medium was refreshed every day and cells were passaged every 4–5 days using 1 mg/mL Dispase II (Thermo Fisher Scientific). Cells were routinely tested for mycoplasma contamination (17) (18)(Brighi et al., 2021; Cordella et al., 2022b).

### 2.2 Generation of cortical organoids

Cortical organoids had been generated through modification of an established protocol (19) (Sloan et al., 2018). Briefly, hiPSCs were treated with Accutase (Innovate Cell Technologies, AT-104) at 37°C for 4-7 minutes and dissociated into single cells. To obtain uniformly-sized spheroids, AggreWell800 (STEMCELL technologies) containing 1800 microwells were used (800um/each). Approximately 2 × 10^6^ single cells were added per AggreWell800 well in mTeSR PLUS medium (STEMCELL technologies) supplemented with 20µM ROCK inhibitor Y-27632 (STEMCELL technologies), centrifuged the plate at 100g for 3 minutes to capture the cells in the microwells and incubated at 37°C with 5% CO_2_. The seeding day is defined as day 0 (D0). After 24 hours (D1), half of the medium was replaced with neural induction medium (NIM) composed of DMEM/F-12 (1:1) (Sigma Aldrich) supplemented with 1X Sodium pyruvate (GIBCO), 1X B27 w/oA (GIBCO) 1X GlutaMAX (GIBCO), 1X N2 supplement (Thermo Fisher Scientific) and 1X MEM non-essential amino acids (NEAA) (GIBCO). NIM was always freshly supplemented with 1 µg/ml heparin (Sigma Aldrich), 1x SB 431542 (1000x; final 10µM) (Biogems), 1x DORSO (400x; final 2.5µM) (Sigma Aldrich), 1x XAV 939 (4000x; final 2.5µM). All the small molecules, including the heparin, were added from D1 to D8 and the medium was changed every day. At D8 organoids were harvested by firmly pipetting (with a cut the end of a P1000 tip) medium in the well up and down and transferred into 60mm ultra-low attachment plastic dishes (Corning, 3262). The spheroids were resuspended in 5ml of neural differentiation medium (NDM) containing Neurobasal basic (Thermo Fisher Scientific), 1x B-27 w/oA, 1x GlutaMax, and kept in floating condition by shaking the cultures on an orbital shaker (40-60 RMP). The NDM was supplemented with 20 ng/ml FGF2 (R&D Systems) and 1ug/ml heparin with medium change every other day. At around D30, to promote differentiation of the neural progenitors into neurons, FGF2 and heparin were replaced with 20 ng/ml BDNF (Peprotech) and 20 ng/ml GDNF (Peprotech), 20 ng/ml ascorbic acid. The medium was changed every three days. From D50 to D100 NDM was modified by replacing the Neurobasal basic with Neurobasal A. At this point, only 20 ng/ml BDNF (Peprotech) was freshly added and the medium was changed every four days until further characterization. BZ (B7273 - Merck) treatment was performed by adding 5 uM BZ to the culture medium from D70 to D100.

### 2.3 Dissociation of cortical neurons from cortical organoids

To obtain 2D cortical cultures we followed a published protocol (20) (Sakaguchi et al., 2019), up to 5 brain organoids at D70 were collected and treated with Accutase (Innovate Cell Technologies, AT-104) at 37°C for 4-7 minutes and gently dissociated into single cells. Cells were then centrifuged at 300g for 5 minutes and resuspended in 2 ml of NDM-A medium supplemented with 20 ng/ml BDNF (Peprotech) and plated onto poly-L-ornithine/laminin-coated dishes at a density of 80,000 cells per cm^2^. The medium was replaced every other day.

### 2.4 Immunostaining, image acquisition and analysis of neuronal and astrocytic markers

Cortical organoids were collected at D30 and 100, incubated in 4% PFA solution for 2 hours and then placed in PBS with 30% sucrose overnight at 4 °C. The following day, the sucrose solution was removed and organoids were placed on OCT (at least 3 organoids on each OCT drop) and cryosectioned using a standard cryostat (Leica CM1860, Leica Biosystems). 20 μm thick sections were collected on Ultra Plus slides (Thermo Fisher Scientific) and stored at +4 °C. For immunostaining, organoid sections were quickly washed for 10 min in PBS and then incubated for 40 min with a warm antigen retrieval solution (95°C) containing 1X Citrate buffer, at pH 6.0 (Sigma Aldrich). After a double washing of 5 minutes in PBS with Tween-20 (0.1%), sections were blocked in 0.3% Triton-X and 5% goat serum, 1% BSA and 200mM Glycine (Sigma Aldrich) in PBS for 1 h. Organoid slices were then incubated overnight with primary antibodies in 0.3% Triton-X and 5% goat serum in PBS at the following dilutions: mouse anti-PAX6 (MA1-109 Invitrogen, 1:100), chicken anti-GFAP (173006 Synaptic System, 1:300), chicken anti-MAP2 (ab5392 Abcam, 1:2000), rabbit anti-TBR1 (20932-1-AP Proteintech, 1:150), rabbit anti-β-TUBULIN III (TUJ1) (T2200 Sigma Aldrich, 1:1000), rat anti-CTIP2 (ab18465 Abcam, 1:200), rabbit anti-synapsin 1 (D12G5, 1:500, Cell signaling), rabbit anti-synaptogyrin (PA556226, 1:200, Invitrogen), mouse anti-4R Tau (MMS-5020, 1:200, Biolegend), mouse anti-AT8 (MN1020, 1:200, Invitrogen), anti phospho Tau181 (MM0194, 1:500, Medimabs) (Supplementary table 1). The day after, the primary antibody solution was washed out and slices were incubated with secondary antibodies for 2 h at room temperature in a dark room. AlexaFluor secondary antibodies (Thermo Fisher Scientific) were used at the concentration of 1:500 (Supplementary Table 2), and Hoechst (Sigma Aldrich, 1:500) was used for nuclei staining. Finally, after 3 washes with 0.1% Tween-20 in PBS, organoid sections were mounted with ProLong Diamond Antifade Mountant (Thermo Fisher Scientific) and sealed with nail polish. The slides were stored at +4 °C until acquisition. Images were acquired with an inverted microscope Olympus IX73 equipped with the X-Light V3 Spinning Disk Confocal module (Crest Optics), LDI laser source and a Prime BSI Scientific CMOS (sCMOS) 6.5um pixels (Photometric). All the images were acquired with a 60x/NA 1.35 and 100x/NA 1.45 oil objectives (Olympus) in stack with z-step of 0.2 μm exploiting the Metamorph software version 7.10.2 (Molecular devices, Wokingham, UK). Fluorescence images of PAX6, CTIP2, TBR1 and GFAP staining (images acquired with a 60x objective) were analyzed through a custom code developed in MATLAB environment. Mean Z-projections over 20 planes of the image stacks were subjected to a pre-processing step consisting of the following chain of operations: median filtering (sigma = 5 pixels), background removal, H-minima transformation and contrast-limited adaptive histogram equalization (CLAHE). More precisely, the median filter was used to smooth the images and reduce the noise, the background level was obtained with a histogram shape-based method beyond the peak of the intensity distribution, H-minima transform (H=0.05) was additionally applied for further denoise, and CLAHE method was used to enhances the contrast of the grayscale image. Subsequently, processed images were binarized using Otsu thresholding method, which automatically identified the optimal intensity threshold to divide the images into foreground and background regions. This procedure enabled us to finally evaluate the fraction of area covered by the signals and the corresponding integrated density.

### 2.4 Tridimensional analysis of neuronal cytoskeleton within cortical organoids

Fluorescence 3D image stacks of the cytoskeletal marker MAP2, acquired with a 100x objective, were segmented to perform morphological analysis of the cytoskeletal network using Huygens and MATLAB 2022b software. Before proceeding with binarization, we used Huygen Deconvolution Wizard to apply images deconvolution as a pre-processing step to improve the contrast and resolution of the digital images recorded. The software automatically generated a theoretical P.S.F. computed from the microscope parameters. The resulting deconvolved 3D images were further processed and segmented with an automatic code developed in MATLAB, described in the following. Gaussian three-dimensional filtering (sigma = 3 pixels), local background subtraction and Hessian-based Frangi vesselness filter (r=10 pixels) were applied to enhance tube-like structures in the volumetric images data and extract the binary masks from the enhanced images via Otsu thresholding method. Finally, the resulting binary images were skeletonized via morphological thinning algorithms. We quantified the total number of separated fragments over the volume scanned and the average length of the fragments.

### 2.5 Immunostaining and image acquisition and analysis of 2D cultures

Monolayer cortical cultures were fixed in 4% paraformaldehyde (PFA; Sigma Aldrich) solution for 15 min at room temperature and double washed in PBS (Thermo Fisher Scientific). Fixed cells were then permeabilized with PBS containing 0.2% Triton X-100 (Sigma Aldrich) for 15 min and blocked with a solution containing 0.2% Triton X-100 and 5% Goat Serum (Sigma Aldrich) for 20 min at room temperature. Cells were then incubated overnight at 4°C with primary antibodies at the following dilutions: chicken anti-MAP2 (ab5392 Abcam, 1:2000), rabbit anti-β-TUBULIN III (TUJ1) (T2200 Sigma Aldrich, 1:2000), mouse anti-VGluT1 (135303 Synaptic Systems, 1:250), rabbit anti-PSD95 (3450 Cell Signaling, 1:250), mouse anti-gephyrin (147021, Synaptic System, 1:250) and rabbit anti-VGAT (131003, 1:300, synaptic system). The day after, cells were washed twice and incubated with secondary antibodies for 1 h at room temperature in the dark. AlexaFluor secondary antibodies (Thermo Fisher Scientific) were used at the concentration of 1:750 and Hoechst (Sigma Aldrich) was used for nuclei staining. Some samples were incubated with an antibody dilution buffer without primary antibodies to discriminate that the synaptic signals were not confounded with non-specific binding of secondary antibodies in the sample. For synaptic markers quantification of 2D cortical networks, fluorescence images were acquired on an X-Light V3 Spinning Disk Confocal module (Crest Optics) with a 100x/NA 1.45 oil objectives (Olympus) in stack with z-step of 0.2 μm a Prime sCMOS camera (Photometrics), and a MetaMorph software (Molecular Devices). Quantification of synaptic markers has been carried out on Fiji software. Specifically each signal has been adjusted by removing the background noise and then a threshold has been performed to quantify both the area covered by each synaptic signal and the integrated density.

### 2.6 MitoTracker staining and confocal analysis

Mitochondria content and their morphology has been evaluated in live cells using MitoTracker Red FM (Invitrogen/Molecular Probes). 2D cultures have been obtained as previously reported. Briefly, up to 5 brain organoids have been dissociated at D70 and plated onto poly-L-ornithine/laminin-coated dishes at a density of 80,000 cells per cm2. Cultures have been maintained in an appropriate neuronal culture medium (NMD-A). On D100, cells were incubated with MitoTracker Red FM at a concentration of 300 nM for 30 minutes in a CO2 incubator at 37°C and double washed with NMD-A medium, each for 5 minutes at 37°C, followed by two washes with PBS. Monolayer cortical cultures were fixed in 4% paraformaldehyde (PFA; Sigma Aldrich) solution for 15 min at room temperature and double washed in PBS (Thermo Fisher Scientific). Fixed cells were then permeabilized with cold Methanol (Sigma Aldrich) for 7 minutes, then washed three times with PBS for 5 minutes each. Immunofluorescence protocol has been carried out as previously reported. MitoTracker signals were collected using an X-Light V3 Spinning Disk Confocal module (Crest Optics) with a 100x/NA 1.45 oil objectives (Olympus) in stack with z-step of 0.2 μm a Prime sCMOS camera (Photometrics), and a MetaMorph software (Molecular Devices). To analyze the images, the ImageJ software was utilized. Initially, Z-projections were performed by merging approximately 10 planes, which improved the accuracy of the measurements. Following this, background removal was conducted, and brightness and contrast were adjusted to enhance the quality of the mitochondrial signal. Both the amount and lengths of individual mitochondria within axons were then measured using the “freehand” tool in ImageJ, specifically focusing on the axonal region by excluding mitochondria present in the soma. The lengths of individual mitochondria were averaged for each axon to obtain a representative value for further analysis.

### 2.7 Protein extraction and western blot analysis

Cell lysates of D100 cortical organoids were homogenized with RIPA extraction buffer (50mM Tris-HCl pH 7.5, 150mM NaCl, 1% NP40, 0.5% SDS, 0.5% Sodium deoxycholate, 1mM EDTA and 50mM NaF) containing 1 mM PMSF, 1X protease inhibitor cocktail and 1mM DTT (all from Sigma-Aldrich). After incubation on ice for 15 min, samples were centrifuged at 13,000× *g* at 4°C for 30 min, and the supernatant containing total proteins were collected.

Protein concentrations were determined using the Bio-Rad Protein Assay (Bio-Rad laboratories GmbH) following the manufacturer’s instruction. Equal amounts of total protein (20 ug) were separated on 4–12% NuPage Bis-Tris gels (Life Technologies, Carlsbad, CA, USA), transferred to nitrocellulose membranes (Bio-Rad, Hercules, CA, USA) for 1 hour in Tris-Glycine buffer and blocked with in 1 X PBS-0.1% Tween20 solution containing 3% of BSA (Serva). Membranes were then immunoblotted with specific antibodies and developed using ECL Blotting Substrate (Advansta, San Jose, CA, USA). The quantification of protein bands was performed by densitometric analysis using the Image J software (1.47 version, NIH, Bethesda, MD, USA). Three different experiments (differentiation batches) were analyzed. Cortical organoids were pooled to provide population variability within each experiment.

#### Antibodies

To evaluate protein expression on Western blot membranes the following antibodies were used: mouse monoclonal anti-AT8 (1:250) (Invitrogen), mouse monoclonal anti-4R (1:500) (Biologend), rabbit monoclonal anti-Tau (D1M9X) (1:500) (Cell Signaling) and mouse monoclonal anti-β-actin (1:10000) (Sigma Aldrich) as a loading control. Goat anti-mouse IgG-HRP (1:20000) (Bethyl Laboratories) and goat anti-rabbit IgG-HRP (1:20000) (Bethyl Laboratories) were used as secondary antibodies. All primary and secondary antibodies were diluted in a PBS + 0.1% Tween 20 solution containing 3% BSA (SERVA, Reno, NV, USA).

### 2.8 PCR, RT-PCR, and RT-qPCR

Total RNA was extracted with the EZNA Total RNA Kit I (Omega Bio-Tek) and retro-transcribed using the iScript Reverse Transcription Supermix for RT-qPCR (Bio-Rad). Real-time RT-PCR was performed with iTaq Universal SYBR Green Supermix (Bio-Rad) on a ViiA 7 Real-Time PCR System (Applied Biosystems). A complete list of primers is provided in Supplementary material. (Supplementary table 3).

### 2.9 NanoString nCounter Gene Expression Assays and Analysis

nCounter gene expression assay (NanoString Technologies) was performed for the Human Neuropathology panel, including 760 genes covering pathways involved in neurodegeneration and other nervous system diseases, and 10 internal reference genes for data normalization. Briefly, panel codeset probes were hybridized with 100ng of total RNA for 20hr at 65°C according to the manufacturer’s instructions. Data were collected using the nCounter Digital Analyzer (NanoString). RNA counts were normalized using the geometric mean of the seven housekeeping genes (AARS, CCDC127, CNOT10, CSNK2A2, LARS, SUPT7L, TADA2B) that showed an average count >450, after validation against positive and negative controls, using the nSolver 4.0 software (NanoString). The significant Differentially Expressed (DE) genes were selected by multiple t-test performed with Graph Pad Prism 6, with *pvalue* ≤ 0.05 and a fold change cut-off values of ≥1.5 or ≤0.66 for up-modulated and downregulated genes, respectively. Gene ontology (GO) enrichment and pathway analysis were identified according to Database for Annotation, Visualization and Integrated Discovery (DAVID) functional annotation. The heat maps were performed for Hierarchical cluster using Average linkage and Pearson Distance Measurement Methods of Heatmapper tools (http://www.heatmapper.ca).

### 2.10 Calcium imaging recordings and data processing on whole cortical organoids

Calcium imaging acquisition on the whole cortical organoid was performed at room temperature using a custom fluorescence imaging microscope. The high-affinity calcium-sensitive dye Fluo4-AM (Invitrogen) was excited at a wavelength of 490 nm with the stable light source Lambda XL (Sutter Instrument) equipped with a Lambda 10-B optical filter changer (Sutter Instrument). The emitted light was collected through a 525/50 nm filter. The fluorescence imaging was performed using a Zeiss Axio observer A1 inverted microscope (Zeiss) with a Zeiss A-Plan 20x/NA 0.25 infinity corrected objective (Zeiss) and a CoolSNAPHQ2 camera (Photometrics). Images were acquired for 5 minutes at a sampling rate of 4 Hz and exposure time of 20 ms using Micromanager software. Neuronal cultures were incubated with the dye at a concentration of 5 μM for 30 minutes at 37°C in HEPES-buffered external solution (NES) containing 140 mM NaCl, 2.8 mM KCl, 2 mM CaCl_2_, 2 mM MgCl_2_, 10 mM HEPES, and 10 mM D-glucose (pH 7.3 with NaOH; 290 mOsm). Calcium imaging data were processed using custom MATLAB codes. The first step involved identifying the regions of interest corresponding to neurons. An automated algorithm was developed to analyze the cumulative difference of the signal between frames of the time series in the frequency domain using 2D Fourier transform. The resulting matrix was filtered to remove high-frequency noise, and the local maxima of the matrix were identified as neurons. Once the cell positions were determined, the fluorescence signals over time were collected. The calcium intensity traces were baseline corrected, normalized as ΔF/F_0_, and smoothed using a moving average filter. The processed traces were then analyzed to detect calcium transients and their characteristics. The criteria for automatic detection of a calcium transient were as follows: at the onset, both the fluorescence intensity and the slope of the trace showed an increase, and at the offset, the slope of the trace decreased with a maximum time interval occurring between the onset and offset. The default threshold for peak amplitude was set to 2%. The algorithm provided the option to manually adjust any false or missing detections in the identified neuron positions and calcium transients on the filtered traces. Once the peaks were detected, an exponential fitting procedure was used to obtain amplitude, rising time, and decay time. The rising time estimation was particularly important for distinguishing fast calcium transients associated with neuronal activity from slower calcium signals originating from internal stores or other cell types. A threshold of τ* = 1.5 s was established to recognize and discard non-neuronal signals. Amplitude, frequency, fraction of active neurons, and network synchrony (measured as the relative number of simultaneous events) were evaluated, exported to Microsoft Excel and compared for different cell lines.

### 2.11 Statistical analysis

Statistical analysis, graphs, and plots were generated using GraphPad Prism 9 (GraphPad Software) and MATLAB 2016b (MathWorks). To verify whether our data sets were reflecting normal distribution, the Shapiro–Wilk normality test was performed. Where the normality distribution was not fulfilled, statistical significance analysis was performed using the nonparametric two-sided Mann–Whitney test (MW test, *P* = 0.05). In all other cases, whether not stated otherwise, Student *t* test (*P* = 0.05) was performed. Data set are given as mean ± standard error of the mean (s.e.m.), the number of cells, replicates, fields of views, cultures, and organoid batches are reported for each experiment in the corresponding figure legend.

## 3 RESULTS

### 3.1 Tau Isoforms Dysregulation and Neurites Morphology Impairment in MAPT-Mutant 3D Cortical Organoids

We examined pathological tau species throughout the organoid’s growth phases to validate our 3D cortical organoid system as an effective and realistic in vitro microphysiological model for studying tauopathy. Our goal was to evaluate neuronal differentiation and maturation within the model and assess the influence of the familiar FTD-linked IVS 10+16 MAPT mutation. Building upon the established protocol (19) by Sloan et al. (2018), we promoted neuronal and glial differentiation and cortical organoid maturation by integrating heparin, ascorbic acid, BDNF, and GDNF to foster the proliferation and maturation of neuroepithelial-like stem cells through the stimulation of collagen production (21)(22) (Bai et al., 2021; Cortes et al., 2016). (Figure 1A, Supplementary Figure 1).

**Figure 1.**
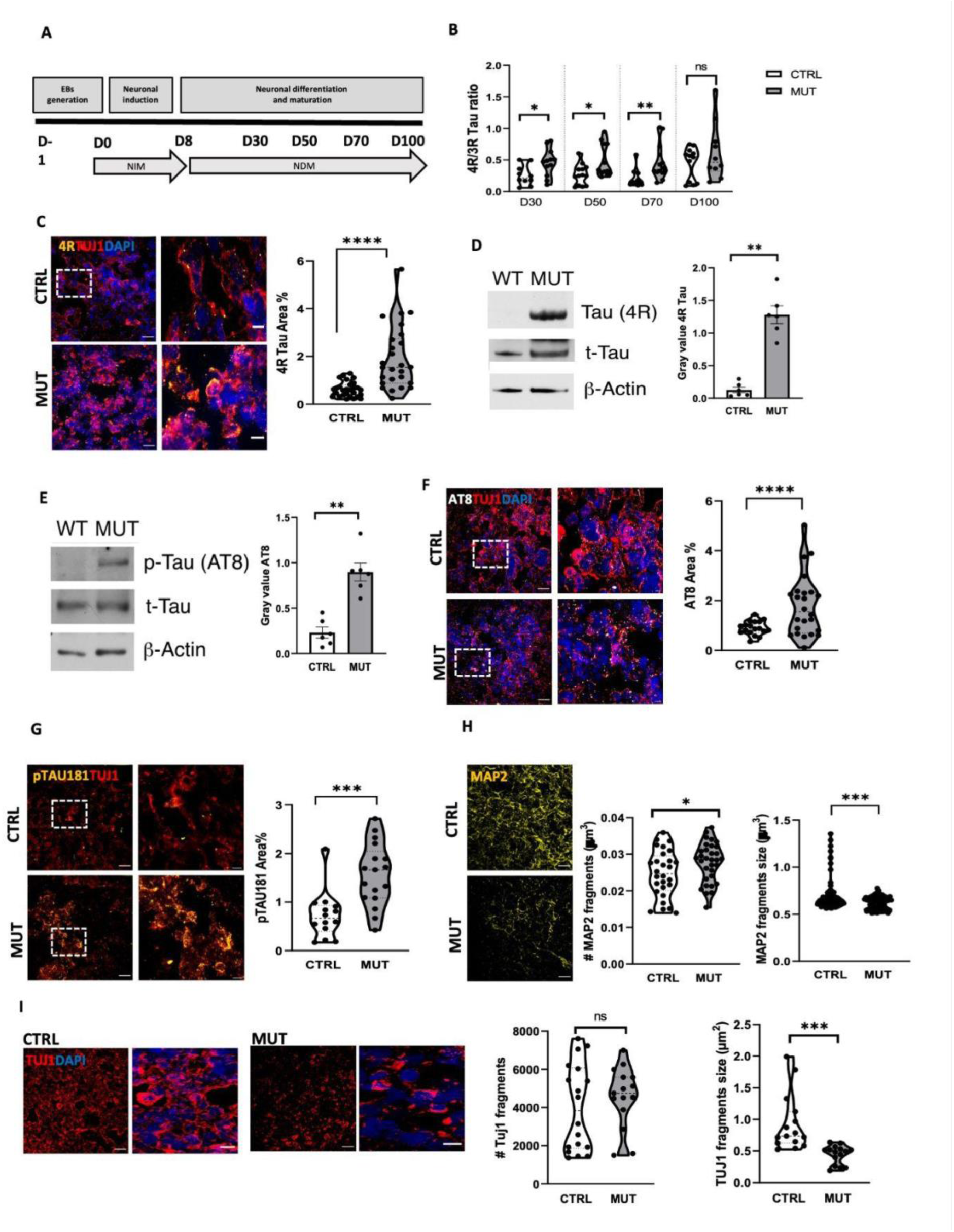
MAPT IVS10+16 Mutation Alters Tau Isoform Balance and Neurite Morphology in Cortical Organoids. **A)** Protocol timeline for the generation of cortical organoids, indicating the corresponding medium used at each developmental stage. **B)** Violin plot showing the quantitative Real-time PCR analysis of the 4R/3R tau ratio in control (CTRL) and tau-mutant (MUT) iPSC-derived cortical organoids at different time points (D30, 50, 70, and 100; CTRL n=18/8, MUT n=16/7; replicates (10 organoids each)/batches for each time point). Gene expression is normalized to the housekeeping gene ATP5O (\*\**p<0.001; *p<0.05, Mann-Whitney (MW) test*). **C)** Left: Representative fluorescence images of CTRL (top) and MUT (bottom) iPSC-derived cortical organoids at D100, immunostained for the 4R tau isoform (yellow), neuronal beta3 tubulin (TUJ1, red), and Hoechst for nuclei visualization (blue). Scale bar, 30 μm. Right: Violin plot representing the 4R tau signal as a percentage of the area covered within the field of view (FOV) in D100 CTRL and MUT cortical organoids (CTRL n = 27/3 FOVs/batches; MUT, n = 26/3 FOVs/batches; ***p<0.0001, MW test). **D)** Left, representative immunoblot of 4R tau isoform. Right, scatter dot plot showing the amount of 4R protein in D100 CTRL and MUT cortical organoids. Actin was used as a protein loading control, and total tau was used to normalize the signal. Values are expressed as median ± sem from 4 independent experiments (batches); *p <0.05, MW test. **E)** Left, representative immunoblot of phosphorylated tau at Ser202/Thr205 (AT8). Right, scatter dot plot reporting the amount of AT8 protein in D100 control CTRL and Tau-mutant (MUT) cortical organoids. Actin was used as a protein loading control, and total tau was used for normalization. Values are expressed as median ± sem from 4 independent experiments (batches); * p <0.05, MW test. **F)** Left, representative confocal images of control (CTRL, top) and Tau-mutant (MUT, bottom) cortical organoids at D100 immuno-stained for anti-AT8 (Grey), Tuj1 (red) and HOECHST (blue) for nuclei visualization. Right, violin plot representing the AT8 signal quantification as % of the area covered within the field of view (FOV) (CTRL n = 18/2 FOVs/batches; tau-MUT n = 24/3 FOVs/batches; *** p<0.0001 MW test;). **G)** Left, representative confocal images of control (CTRL, left) and Tau-mutant (MUT, right) cortical organoids at D 100, immunolabeled for pTau181 (yellow) and Tuj1 (red) (Scale bar 30um). Right, violin plot representing the amount of pTau181 signal as % of the area covered within the field of view (FOV). (CTRL n= 15FOVs/2 batches; MUT n= 18 FOVs/3 batches; ***p<0.01 MW test). **H)** Left, representative confocal images of control (CTRL, left) and Tau-mutant (MUT, right) cortical organoids at D 100, immunolabeled for MAP2 (yellow) (Scale bar 30um). Right, violin plots showing the quantification of the MAP2 signal as the number of fragments (CTRL n=71/5; tau-MUT n=75/5. ** p<0.001 MW test) and the fragment size/um^3^ (CTRL n=71/5; tau-MUT n=75/5. *** p<0.0001 MW test). **I)** Left, representative confocal images of control (CTRL, left) and Tau-mutant (MUT, right) cortical organoids at D 100, immunolabeled for TUJ1 (red) and Hoechst for nuclei visualization (Blue) (Scale bar 30um). Right, violin plots representing the quantification of the TUJ1 signal as the number of fragments and the size of the fragments (per um^2^) analyzed within the field of view (FOV). (CTRL n=15/2; tau-MUT n=15/3. NS p>0.05 MW; ***p<0.001 MW).

Initially, we investigated if the intronic IVS 10+16 MAPT mutation affected the balance between 3R and 4R tau isoforms during the maturation of cortical organoids. We performed Real-Time PCR analysis at different time points from D30 to D100 of organoid development (Figure 1A). The gene expression analysis indicated a skewed ratio favoring a higher expression of 4R over 3R tau isoforms in the tau-mutant organoids relative to their isogenic controls (Figure 1B). This imbalance was also observed at the protein level through immunofluorescence (Figure 1C) and western blot (Figure 1D) analyses, revealing an increased expression of the 4R isoform in D100 tau-mutant organoids. As it is known that the MAPT IVS 10+16 mutation, disrupting the equilibrium between 3R and 4R tau isoforms, leads to an increase in pathological tau variants (6)(7)(23)(Vereyen et al., 2018; Kopach et al., 2021, Young et al., 2021), we further evaluated the presence of hyperphosphorylated tau in our in vitro disease model. This was accomplished using antibodies targeting phosphorylated tau at various serine and threonine sites. Western blot analysis revealed a significant increase in phosphorylated tau at Ser202/Thr205 (AT8) in tau-mutant organoids (D100) compared to controls (Figure 1E). Immunofluorescence analysis confirmed the increase in AT8 staining (Figure 1F) and also revealed the increase of phosphorylated tau at Thr181 (pTau181)(Figure 1G).

Considering tau’s critical role in microtubule stabilization and how its hyperphosphorylation undermines this function (24)(25)(26)(27) (Wang and Mandelkow, 2016; Nataliya et al., 2019; Viet et al., 2023; Mueller et al., 2021), we also evaluated the neurites morphology by looking at neuronal cytoskeletal proteins. Immunostaining for MAP2 (Figure 1H) and β3-tubulin (TuJ1; Figure 1I) revealed that neurites in tau-mutant organoids displayed a highly fragmented morphology with smaller fragments than those in isogenic controls, indicating that the FTD-related tau mutation impairs neuronal integrity and/or neurite elongation at early developmental stages.

These findings indicate that the intronic MAPT IVS 10+16 mutation triggers imbalances in tau isoform levels and elevated tau phosphorylation from the early stages of development, potentially affecting the neuronal cytoskeleton in 3D cortical organoids.

### 3.2 Intronic MAPT IVS 10+16 mutation impairs Cortical Organoid Development and Maturation

The observation that this intronic tau mutation impacts neurites morphology at D100 of cortical organoid maturation, combined with tau’s established role in shaping different aspects of neuronal development, such as axonal guidance, synaptogenesis, and neuronal migration (24)(Wang and Mandelkow, 2016), led us to investigate the mutation’s effect on key genes at two critical organoids developmental points (D50 and D100). By applying nCounter Nanostring technology for total RNA examination in both the control and tau-mutants organoids using the Neuropathology Panel (which includes 760 genes), we detected significant gene expression alterations: 130 genes at D50 and 243 genes at D100 were differentially expressed due to the tau mutation (p<0.05) (See Supplementary Figure 2A, B, and Supplementary Table S4 and S5).

Initially, we analyzed gene expression modulation throughout the development of organoids obtained from both iPSC lines. In the control organoids, we noted a differential expression of 183 genes from D50 to D100 (Supplementary Figure 3A, Supplementary Table S6).

Gene ontology (GO) term enrichment analysis of the 164 upregulated genes at D100 with respect to D50 showed categories chiefly associated with synaptic transmission (biological processes, GO: BP), protein binding (molecular function, GO: MF), and neuronal shape development (cellular component, GO: CC)). Conversely, there was a reduction in 19 genes, predominantly related to the cell cycle (Supplementary Figure 3B), indicating cortical organoids’ maturation.

Within the tau-mutant organoids, we identified alterations in the expression levels of 180 genes at D100 compared to D50, with 114 genes upregulated and 66 genes downregulated (see Supplementary Figure 4A, Supplementary Table S7). These changes were only partially related to neuronal maturation and differentiation. Among the upregulated genes, categories associated with synaptic transmission were observed (Supplementary Figure 4B); however, these categories were also present among the downregulated genes (Supplementary Figure 4B). Furthermore, there was no downregulation of genes related to the cell cycle, indicating a persistent proliferation state in mutant organoids at D100.

We then analyzed the genes differentially expressed at D100 between control and tau-mutant organoids (tot 243 genes) (Supplementary Figure 2B and Supplementary Table S5). Gene ontology (GO) term enrichment analysis of the down-regulated subset at D100 in tau-mutant organoids with respect to their isogenic controls (163 genes) revealed categories mainly related to excitatory and inhibitory synapses, neuronal projection, dendrites, and axon (GO: CC); calcium homeostasis, glutamate and GABA homeostasis and signaling (GO: MF); nervous system development (GO: BP) (Figure 2A, left). When looking at the 80 upregulated genes in tau-mutant organoids, the GO term enrichment analysis showed categories mainly related to the centrosome, basolateral plasma membrane, and cytoplasm (GO: CC); protein, and nucleic acid binding (GO: MF); positive regulation of cell proliferation and intracellular cascades linked to cellular stress (GO: BP) (Figure 2A, right). These data suggest a delay in tau-mutant cortical organoids maturation and indicate that tau targets include genes that play crucial roles in neuronal and neuronal network development and maturation.

**Figure 2.**
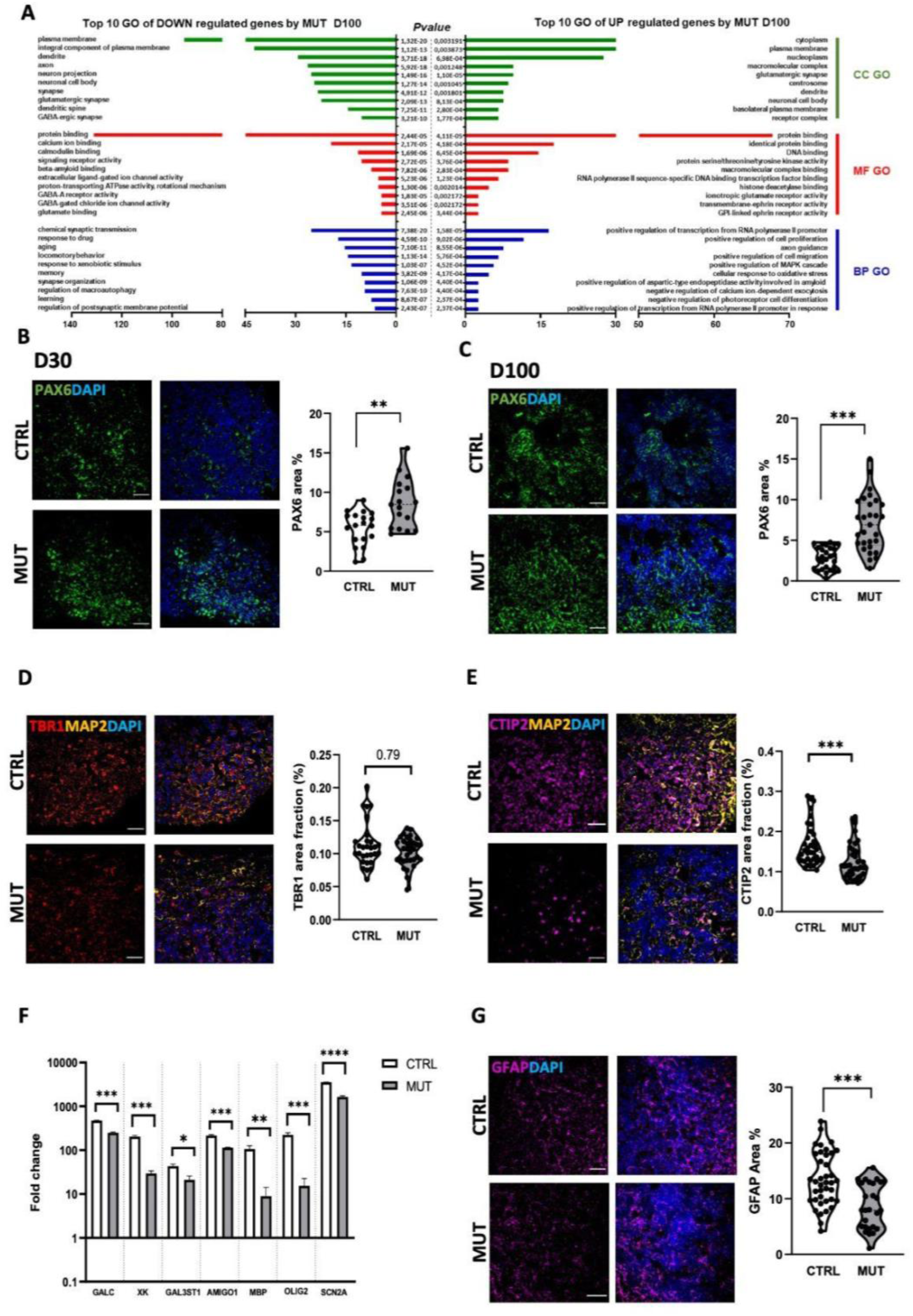
Intronic Tau mutation delays cortical maturation. **A)** GO enrichment analysis of down (left)- and up (right)-regulated genes by D100 MUT respect to CTRL organoids analyzed by DAVID database. An overview of top 10 significantly enriched terms in three categories: biological process (BP), cellular component (CC), and molecular function (MF). Number of genes involved in a GO process is shown in the x-axis. The cut-off of p-value was set to 0.05. **B)** Left, representative confocal images of CTRL (top) and MUT (bottom) cortical organoids at D30 immunolabeled for neuronal progenitor marker PAX6 (green) and Hoechst (blue) for nuclei visualization. Scale bar, 30 μm. Righ, violin plot representing the amount of PAX6 signal as % of the area covered within the field of view (FOV) (CTRL n =20/2 FOVs/batches; tau-MUT n =17/2 FOVs/batches; MW test **p<0.01). **C)** Left, representative confocal images of CTRL (top) and MUT (bottom) cortical organoids at D100 immunolabeled for neuronal progenitor marker PAX6 (green) and Hoechst (blue) for nuclei visualization. Scale bar, 30 μm. Righ, violin plot representing the amount of PAX6 signal as % of the area covered within the field of view (FOV) (CTRL n =30/3 FOVs/batches; MUTANT n =29/3 FOVs/batches; MW test p****<0.001). **D)** Left, representative confocal images of D100 CTRL (top) and MUT (bottom) cortical organoids immunolabeled for neuronal marker TBR1 (red), MAP2 (yellow), and Hoechst (blue) for nuclei visualization. Scale bar, 30 μm. Right, violin plot representing TBR1 signal as % of the area covered within the field of view (FOV) (CTRL n =29/2 FOVs/batches; tau-MUT n =33/3 FOVs/batches; MW test NS p=0.05). **E)** Left, representative confocal images of CTRL (top) and MUT (bottom) derived cortical organoids at D100 immunolabeled for neural marker CTIP2 (magenta), MAP2 (yellow) and Hoechst (blue) for nuclei visualization. Scale bar, 30 μm. Right, violin plot representing the quantification of CTIP2 signal as % of the area covered within the field of view (FOV) (CTRL n =37/2 FOVs/batches; tau-MUT n =38/3 FOVs/batches; MW test p***<0.001). **F)** Scatter dot plot showing the expression level of genes related to the myelination process in D100 CTRL and MUT organoids (CTRL n =3/1 batches; tau-MUT n =3/1 FOVs/batches; p***<0.001, **p<0.01, *p<0.05 MW test). **G)** Left, representative confocal images of CTRL (top) and MUT (bottom) derived cortical organoids at D 100 immunolabeled for glial marker GFAP (magenta), and Hoechst (blue) for nuclei visualization. Scale bar, 30 μm. Right, violin plot representing the quantification GFAP signal as % of the area covered within the field of view (FOV) (; CTRL n =93/3 FOVs/batches; tau-MUT n =62/3 FOVs/batches; MW test p***<0.001).

At the protein level, immunofluorescence analysis of the neural progenitor marker PAX6 showed higher levels of PAX6-positive cells in tau-mutant cortical organoids at both D30 (Figure 2B) and D100 (Figure 2C) compared to their isogenic control, confirming a delay in the maturation process and a delayed persistent presence of proliferative progenitors during organoids maturation, as also supported by the higher expression levels of cell cycle-related genes PCNA and CCNA2 (Supplementary Figure 5). Looking at the neuronal fate commitment during organoid maturation, we analyzed the labeling for neuronal cortical markers TBR1 and CTIP2. We observed that most cells were positive for the postmitotic neuronal marker TBR1 in both control and tau-mutant organoids (Figure 2D), suggesting that the initial fate commitment was similar in the two lines. Conversely, D100 tau-mutant organoids were characterized by fewer layer 6 cortical neurons (CTIP2-positive cells) than controls (Figure 2E), suggesting that the intronic MAPT IVS 10+16 mutation delays neuronal maturation and subtype specification, further impacting cortical development. Furthermore, we also observed that the glial maturation was delayed in tau-mutant organoids, with downregulation of genes related to the myelination process (GALC, XK, GAL3ST1, AMIGO1, MBP, OLIG2, SCN2A, Figure 2F), and reduced GFAP protein expression at D100 (Figure 2G).

These data indicate that tau IVS 10+16 mutation strongly delays cortical development affecting both the neuronal and the glial maturation.

### 3.3 Synaptic and Network Functionality Deficits in Tau-Mutant Cortical Organoids

Among the genes differentially expressed in tau-mutant organoids at D100 we observed a significant downregulation of transcripts associated with both glutamatergic and GABAergic synapses, as well as those linked to synaptic structure (Figure 3A). Immunofluorescence analysis of the pan synaptic markers synapsin1 and syngirin1 revealed reduced expression levels of both proteins in tau-mutant cortical organoids (Figure 3B, 3C), confirming the nanostring data indicating impaired synapse formation.

**Figure 3.**
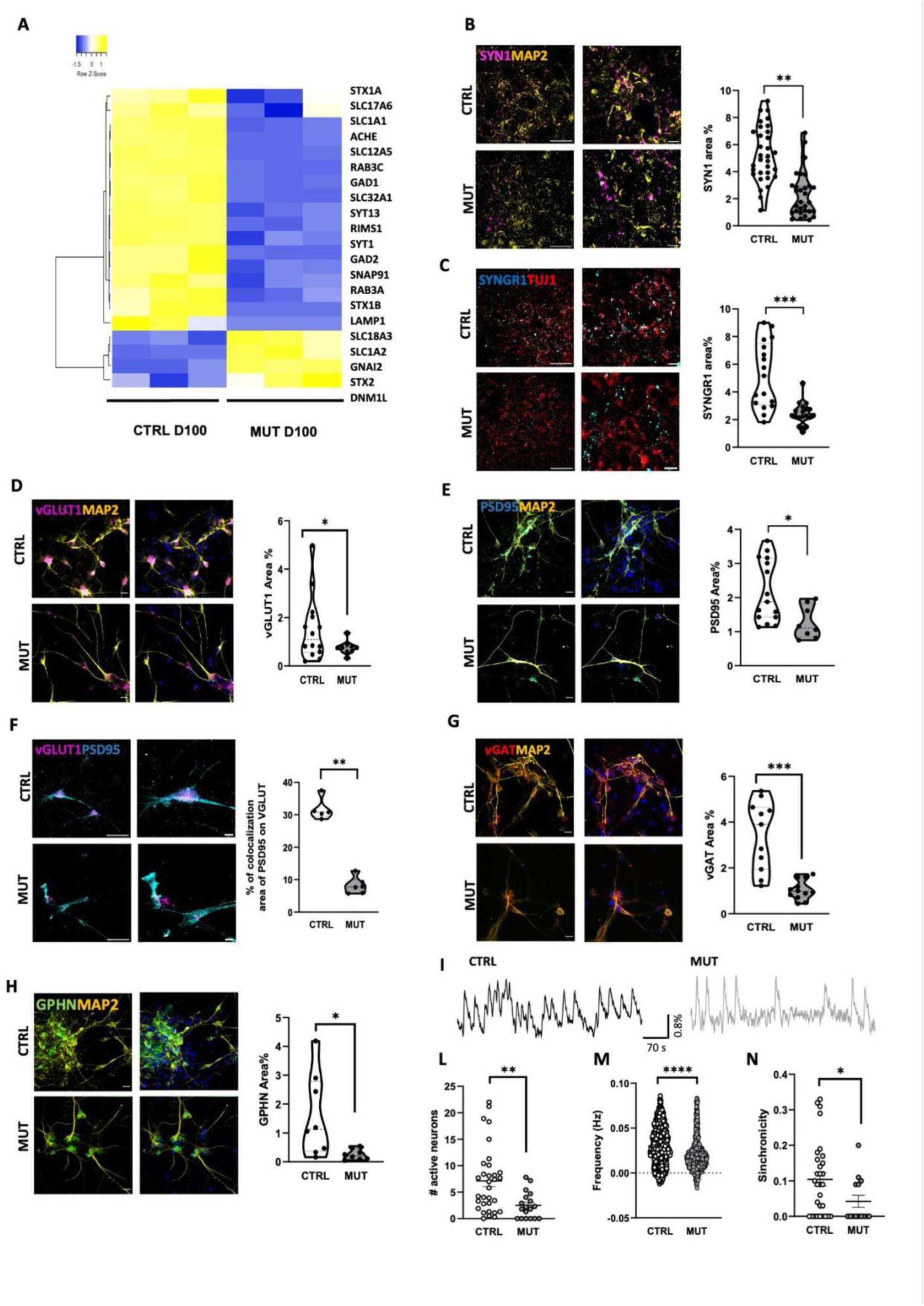
IVS10+16 tau mutation impairs synaptic maturation. **A)** Heat map of unsupervised hierarchical clustering of 21 «Synaptic Pathways» genes between D100 MUT and CTRL organoids. Average linkage and Pearson Distance Measurement Methods was performed for Hierarchical cluster using Heatmapper tools (http://www.heatmapper.ca). The significant genes are selected by multiple t-test performed with Graph Pad Prism 6, with p value ≤ 0.05. **B)** Left, representative confocal images of D100 CTRL (top) and MUT (bottom) cortical organoids immunolabeled for synapsin1 (SYN1) (magenta) and MAP2 (yellow). Scale bar 30µm and 5µm. Right, violin plot representing SYN1 signal as % of the area covered within the field of view (FOV) (CTRL n=32/3 FOVs/batches; tau-MUT n=30/4 FOVs/batches; MW test *p<0.05). **C)** Left, representative confocal images of CTRL (top) and MUT (bottom) derived cortical organoids immunolabeled for Synaptogyrin-1 (SYNGR1) (cyan) and MAP2 (yellow) at D100. Scale bar 30µm and 5µm. Right, violin plot representing the amount of SYNGR1 signal as % of the area covered within the field of view (FOV) (CTRL n = 17/3 FOVs/batches; MUT n = 20/4 FOVs/batches; ***p < 0.001; MW test). **D)** Left, representative images of control CTRL (top) and MUT (bottom) 2D cortical neurons immunolabeled for anti-VGluT1 (magenta), MAP2 (yellow) and Hoechst for nuclei visualization (blue) at D100. Scale bar 30µm. Right, Bar chart (violin plot) representing VGluT1 signal as % of the area covered within the field of view (FOV) (*p < 0.05 MW test; CTRL n = 14/3 FOVs/batches; MUT n = 9/1 FOVs/batches). **E)** Left, representative images of CTRL (top) and MUT (bottom) derived 2D cortical neurons immunolabeled for anti-PSD95 (cyan), MAP2 (yellow) and Hoechst for nuclei visualization (blue) at D100. Scale bar 30µm. Right: Bar chart (violin plot) representing PSD95 signal as % of the area covered within the field of view (FOV) (*p < 0.05 MW test; CTRL n = 13/3 FOVs/batches; MUT n = 10/1 FOVs/batches). **F)** Left, representative images of PSD95 and VGluT1 colocalization in CTRL (top) and MUT (bottom) derived 2D cortical neurons. Right: Bar chart (violin plot) representing VGluT1 and PSD95 colocalization signal as % of the area covered within the field of view (FOV) (**p < 0.01 MW test; CTRL n = 5/2 FOVs/batches; MUT n = 6/2 FOVs/batches). **G)** Left, representative images of CTRL (top) and MUT (bottom) derived 2D cortical neurons immunolabeled for anti-VGAT (red), anti-MAP2 for neuronal cytoskeleton (yellow) and Hoechst for nuclei visualization (blue) at D100. Scale bar 30µm. Right: Bar chart (violin plot) representing VGAT signal as % of the area covered within the field of view (FOV) (***p < 0.0001 MW test; CTRL n = 11/2 FOVs/batches; MUT n = 12/2 FOVs/batches). **H)** Left, representative images of CTRL (top) and MUT (bottom) derived 2D cortical neurons immunolabeled for anti-Gepherin (green), anti-MAP2 for neuronal cytoskeleton (yellow) and Hoechst for nuclei visualization (blue) at D100. Scale bar 30µm. Right: Bar chart (violin plot) representing Gepherin signal as % of the area covered within the field of view (FOV) (*p < 0.05 MW test; CTRL n = 8/2 FOVs/batches; MUT n = 8/1 FOVs/batches). **I)** Representative traces of spontaneous calcium oscillations recorded CTRL (left) and MUT (right) cortical organoids at D 100 of the differentiation protocol. Dot plots representing **L)** the number of active neurons, **M)** the frequency and **N)** the synchronicity of spontaneous calcium oscillation in CTRL (empty circles) and MUT (filled circles) cortical organoids (*p<0.05, **p<0.01,****p<0.001, MW test; CTRL 47/organoids/batches; MUT 29/organoids/batches).

To deeply investigate the excitatory and inhibitory synapses, immunostaining analysis was performed in cortical neurons obtained after dissociating cortical organoids at D70 and cultured for an additional 30 days (Supplementary Figure 1). Labeling for glutamatergic and GABAergic synaptic markers revealed that tau mutation reduced the formation of both synapses. Specifically, for glutamatergic synapses, we observed lower levels of pre- (VGluT1, Figure 3D) and post-synaptic (PSD95, Figure 3E) proteins, with reduced colocalization (Figure 3F). When we looked at GABAergic synapses, we observed lower levels of pre- (VGAT, Figure 3G) and post-synaptic (Gephyrin, Figure 3H) proteins, suggesting an impaired neuronal network development in tau-mutant organoids.

To test the network functionality, we recorded spontaneous calcium transients in whole control and tau-mutant cortical organoids at D100, loading organoids with Fluo-4M, and acquiring large fields of view (260 × 200 microns) at a high sample rate (4 Hz). Data from 5-minute time-lapse recordings (Supplementary Movie1, Movie2) were collected and analyzed using a custom-made algorithm to identify cells, select active zones, and extract functional properties such as amplitude, frequency, kinetic parameters, and network synchronicity (17)(Brighi et al., 2021). For each field of view, we divided active cells into two populations using the 1.5-second value as the rising time threshold of spontaneous calcium transients, and we analyzed the properties of the spontaneous activity of fast cells (rise time <1.5 s), most probably neurons (Figure 3I). As represented in the dot plots, in tau-mutant organoids, we found a smaller number of active neurons (Figure 3L), characterized by weaker spontaneous activity. Indeed, we found that both the frequency (Figure 3M) and the synchronicity (Figure 3N) of calcium events were significantly lower in tau-mutant organoids with respect to isogenic control. The analysis of the peaks showed that peak amplitude and kinetic parameters were similar in tau-mutant and control organoids (Supplementary Figure 6). Altogether, these data suggest that intronic tau mutation leads to significant impairments in synaptic development and network functionality, highlighting the critical role of tau in maintaining synaptic integrity and neuronal connectivity in cortical organoids.

### 3.4 Molecular Pathways and Mitochondrial Dysfunction in Tau-Mutant Cortical Organoids

To dissect the possible molecular pathways leading to the observed impairment in cortical development, we matched the differentially regulated genes in tau-mutant organoids (at D 100) with the targets of tau described by interactome analysis (9)(Tracy et al., 2022).

We identified 17 “tau signature” genes differentially regulated in tau-mutant organoids at both developmental time points D50 and D100 (Figure 4A). These genes are involved in synaptic vesicle-associated functions, cytoskeletal structure, and mitochondrial and lysosome functions, thus confirming the impairment of neuronal network development and suggesting a possible impairment of mitochondrial functions in tau-mutant cortical organoids.

**Figure 4.**
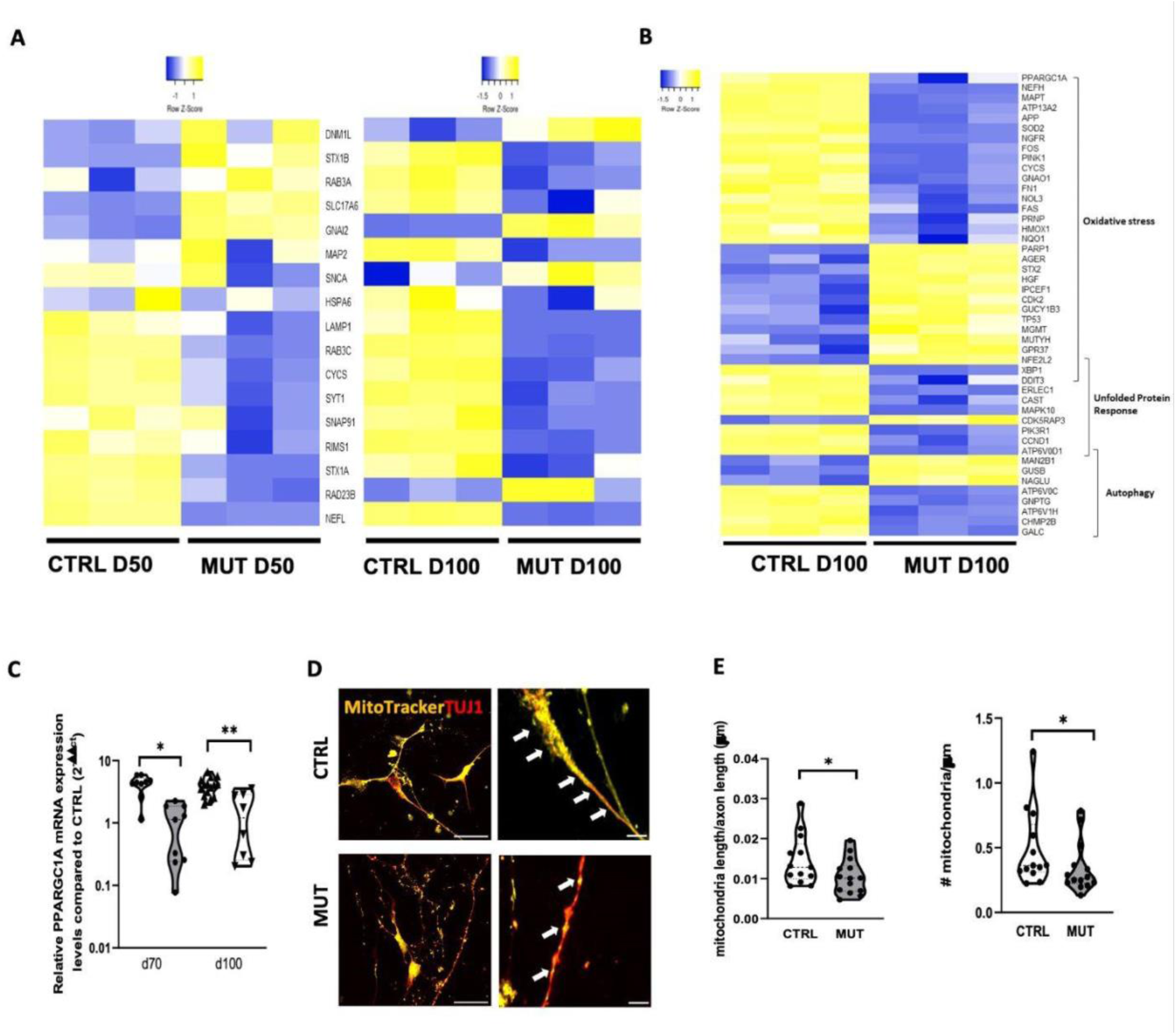
MAPT IVS 10+16 mutation induces mitochondrial alterations. **A)** Heat map of supervised hierarchical clustering of 17 «TAU signature» genes between D50 CTRL vs D50 MUT and D100 CTRL vs D 100 MUT. Average linkage and Pearson Distance Measurement Methods was performed for Hierarchical cluster using Heatmapper tools (http://www.heatmapper.ca). The significant genes are selected by multiple t-test performed with Graph Pad Prism 6, with *p value* ≤ 0.05. **B)** Heat map of supervised hierarchical clustering of 46 «Oxidative Stress», «Unfolded Protein Response» and «Autophagy» genes between D100 CTRL vs D100 MUT. Average linkage and Pearson Distance Measurement Methods was performed for Hierarchical cluster using Heatmapper tools (http://www.heatmapper.ca). The significant genes are selected by multiple t-test performed with Graph Pad Prism 6, with *p value* ≤ 0.05 and a fold change cut-off values of ≥1.5 or ≤0.66 for upmodulated and downregulated genes respectively. **C)** Violin plot showing the quantitative Real-time PCR analysis of PPARGC1A in CTRL and MUT iPSC-derived cortical organoids at different time points D70 (CTRL n = 10/6, MUT n= 9/4; replicates (10 organoids each)/batches for each time point) and D100 (CTRL n = 15/6, MUT n= 8/3; replicates (10 organoids each)/batches for each time point). Gene expression is normalized to the housekeeping gene ATP5O (*p<0.05 MW test; ** p<0.001 MW test;). **D)** Representative images of control CTRL (top) and MUT (bottom) derived 2D cortical neurons immunolabeled with MitoTracker Red FM (yellow) and TUJ1 for neuronal cytoskeleton (red) at D100 (scale bar 30µm; 5 µm). **E)** Left: Violin plot representing mitochondria length in same-length segments of TUJ1-positive branches (*p < 0.05 MW test; CTRL n = 12/2 neurites/batches; MUT n = 13/2 neurites/batches). Right, Violin plot representing the number of mitochondria in same-length segments of TUJ1-positive branches (*p < 0.05 MW test; CTRL n = 12/2 neurites/batches; MUT n = 13/2 neurites/batches).

Indeed, out of the 243 genes differentially expressed at D100 between control and mutant organoids, 46 were associated with autophagy, unfolded protein response, and oxidative stress, all linked to mitochondrial function (Figure 4B).

Our attention was particularly drawn to the significant downregulation of the mitochondrial biogenesis master gene, PPARGC1A, observed D100 in mutant organoids (*p-value* = 0.021) (Figure 4B and Suppplementary Table S5. Although no significant difference in PPARGC1A expression was noted at D50 (Supplementary Table S4), closer examination using Real-Time PCR at the intermediate time point of D70 and final time point of D100 revealed a marked down-regulation of the PPARGC1A transcript in tau-mutant organoids (Figure 4C). This finding supports the nanostring data at D100 and underscores the period between D50 and D70 as potentially critical for cortical development.

We thus employed MitoTracker labeling to evaluate mitochondrial quantity and structure within 2D cortical cultures at D100, following the dissociation of organoids at D70. After mitochondrial labeling, the cells underwent fixation and were stained with the neuronal TUJ1 antibody (Figure 4D). We confirmed that neurons with the tau mutation exhibited heterogeneous TUJ1 staining as in the whole organoids (see Figure 1). Moreover, the analysis of mitochondria in same-length segments of TUJ1-positive branches showed that in tau-mutant neurons, neurites contained fewer and smaller mitochondria (Figure 4E) compared to those in isogenic control neurons, indicating that the intronic MAPT IVS 10+16 mutation impairs mitochondrial biogenesis and maturation in cortical neurons.

### 3.5 Bezafibrate Treatment Mitigates Mitochondrial Dysfunction and Enhances Neuronal Integrity in Tau-Mutant Cortical Organoids

We thus explored the efficacy of pharmacological activation of mitochondrial biogenesis in tau-mutant cortical organoids, characterized by low expression of PPARGC1A, the gene encoding the master regulator protein PGC1A. For this purpose, we employed bezafibrate (BZ), a peroxisome proliferator-activated receptor (PPAR) agonist.

BZ is proposed as a therapeutic option for neurological and mitochondrial disorders and has shown promise in mitigating tau pathology (28)(29) (Dumont et al., 2012; Lin et al., 2022).

Since we observed a timely decrease in PPARGC1A expression during the critical maturation phase (D50 - D70) in tau-mutant organoids, we administered 5 µM BZ within this specific timeframe (Supplementary Figure 1). The impact of BZ treatment was then assessed at D100, with comparative analyses conducted on both treated and untreated tau-mutant cortical organoids, alongside their isogenic controls, derived from parallel experimental batches.

First, we found that BZ treatment effectively normalized mitochondrial parameters in tau-mutant cortical organoids.

Specifically, Mitotracker labeling followed by TUJ1 immunostaining of 2D cortical cultures (Figure 5A) demonstrated that both the quantity and the dimension (Figure 5B) of neuronal mitochondria after BZ application approached those observed in isogenic control samples.

**Figure 5.**
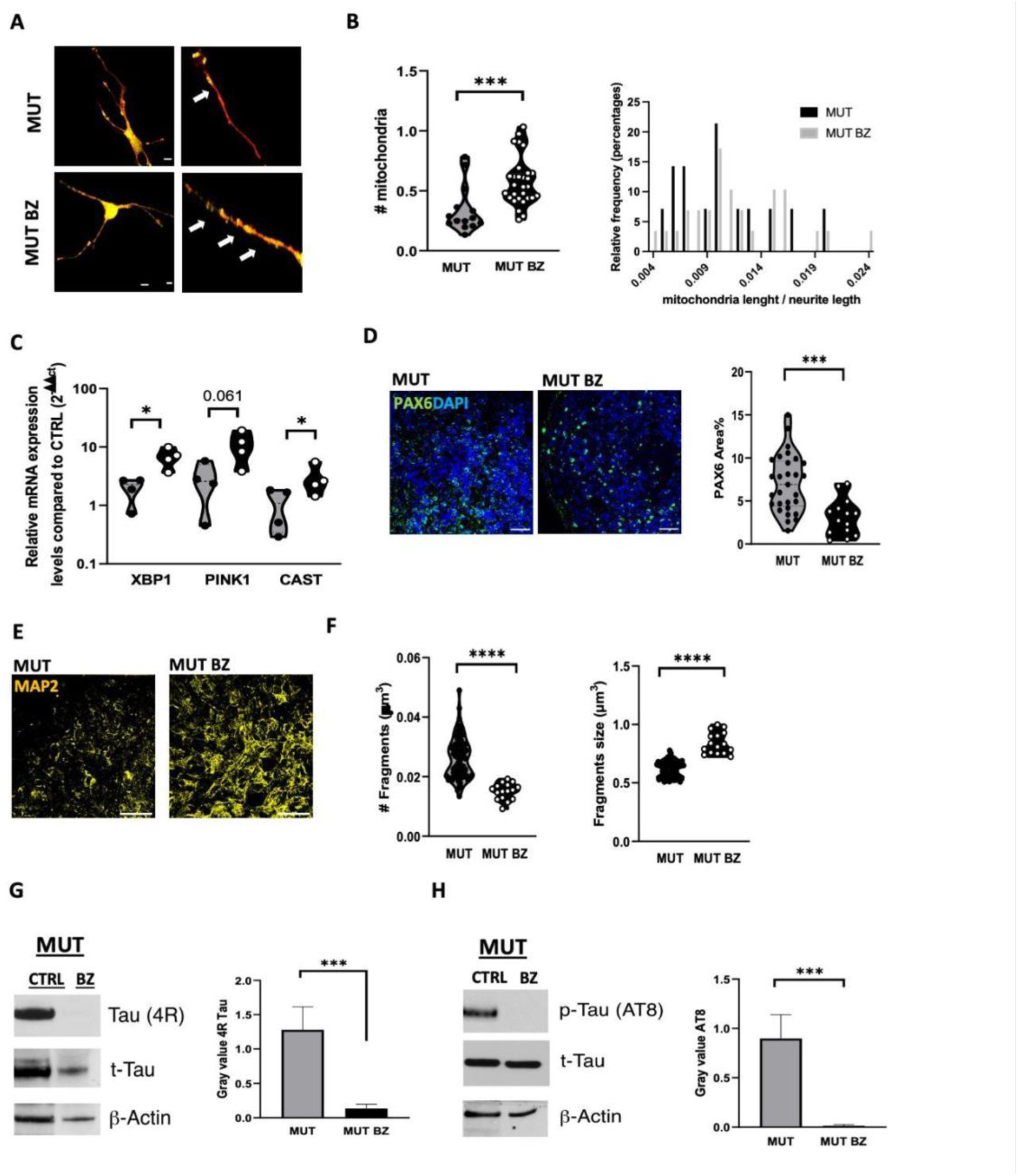
Bezafibrate treatment partially reverses the effects of MAPT IVS 10+16 on mitochondria and neuronal cytoskeleton. **A)** Representative images of untreated Tau mutant (MUT, top) and treated Tau-mutant (MUT BZ, bottom) derived 2D cortical neurons immunolabeled with MitoTracker Red FM (yellow) and TUJ1 for neuronal cytoskeleton (red) at D100 (scale bar 30µm; 5 µm). **B)** Left, Violin plot representing the number of mitochondria in same-length segments of TUJ1-positive branches; Right, bar chart representing the distribution of mitochondria length in same-length segments of TUJ1-positive branches (***p < 0.001 MW test; tau-MUT n = 13/2 neurites/batches tau-MUT BZ n = 29/3 neurites/ batches). (NS p> 0.05) **C)** Violin plot showing the quantitative Real-time PCR analysis of XBP1, PINK1 and CAST gene in MUT and (MUT BZ iPSC-derived cortical organoids at different time points D 100 (MUT n = 4/2, MUT BZ n= 4/2; replicates (10 organoids each)/batches for each time point). Gene expression is normalized to the housekeeping gene ATP5O (*p<0.05 MW test; NS p>0.05MW test). **D)** Right, representative images of MUT (left) and MUT BZ (right) derived iPSC-derived cortical organoids immunolabeled with PAX6 (green) and hoechst (blue) for nuclei visualization at D100 (scale bar 30µm). Left, violin plot representing the amount of PAX6 signal as % of the area covered within the field of view (FOV) (MUT n =29/3 FOVs/ batches; MUT BZ n=16/3; MW test p***<0.01). **E)** Representative images of MUT (left) and MUT BZ (right) derived iPSC-derived cortical organoids immunolabeled with MAP2 (yellow) at D100 (scale bar 30µm). **F)** Left, violin plot showing the quantification of the MAP2 signal as the number of fragments (MUT n=75/5; MUT BZ n=16/2 ** p<0.001 MW test). Right, violin plot showing the quantification of the MAP2 signal as fragment size/um^3^ (MUT n=75/5; MUT BZ n= 16/2 *** p<0.0001 MW test). **G)** Left, representative immunoblot of 4R tau isoform. Right, bar graph showing the amount of 4R protein in D100 untreated Tau-mutant (named as CTRL) and treated Tau-mutant (named as BZ) cortical organoids. Actin was used as a protein loading control, and total tau was used to normalize the signal. Values are expressed as median ± sem from 3 independent experiments (batches); *** p <0.01, MW test. **H)** Left, representative immunoblot of phosphorylated tau at Ser202/Thr205 (AT8). Right, bar graph reporting the amount of AT8 protein in D 100 untreated Tau-mutant (named as CTRL) and treated Tau-mutant (named as BZ) cortical organoids. Actin was used as a protein loading control, and total tau was used for normalization. Values are expressed as median ± sem from 3 independent experiments (batches); *** p <0.01, MW test.

Furthermore, RT-PCR analysis indicated an upregulation of genes associated with oxidative stress, unfolded protein response, and autophagy (XBP1, PINK1, and CAST, Figure 5C) in tau-mutant organoids treated with BZ. These genes previously showed reduced expression (see Figure 4A), whereas PGC1alpha levels remained unchanged (n=8 replicates/4 batches, p=0.94).

We also found that BZ treatment of tau-mutant organoids reduced PAX6 expression to the level of control organoids (Figure 5D), supporting the idea that the rescue of mitochondrial biogenesis favors neuronal differentiation and maturation.

In line with these results, immunostaining for the neurite marker MAP2 demonstrated that BZ treatment rescue the neurite phenotype, decreasing the number of MAP2 fragments (Figure 5 E,F) and augmenting their length (Figure 5F). Moreover, the expression level of the 4R tau isoform (Figure 5G) and the amount of phosphorylated tau at Ser202/Thr205 (AT8) (Figure5H) were restored to control levels, hinting at a beneficial effect on neuronal maturation and branching.

These results suggest that the imparment of mitochondrial biogenesis contributes to reduce neuronal differentiation and maturation and that the rescue of mitochondrial biogenesis restores neuronal development correlated to a physiological tau pattern.

### 3.6 Bezafibrate treatment rescues synaptic maturation and neuronal functions

Based on the findings of BZ efficacy in normalizing mitochondrial function and promoting neuronal development in tau-mutant cortical organoids, we extended our investigation to assess its potential in restoring synaptic connectivity, a crucial aspect of cortical development.

Immunofluorescence analysis revealed that BZ treatment effectively rescued the synapsin1 expression in tau-mutant organoids, achieving levels comparable to those in control organoids by D100 (Figure 6A). Examining glutamatergic and GABAergic synaptic markers in neurons derived from organoid dissociation further supported BZ’s therapeutic potential. Specifically, for glutamatergic synapses, BZ treatment enhanced the expression of both pre-synaptic (VGluT1, Figure 6B) and post-synaptic (PSD95, Figure 6C) proteins, as well as their colocalization (Figure 6D), bringing tau-mutant neuron levels on par with control conditions. In the case of GABAergic synapses, BZ treatment led to an increased expression of the presynaptic marker (VGAT, Figure 6E) in tau-mutant neurons without altering post-synaptic Gephyrin levels (Figure 6F).

**Figure 6.**
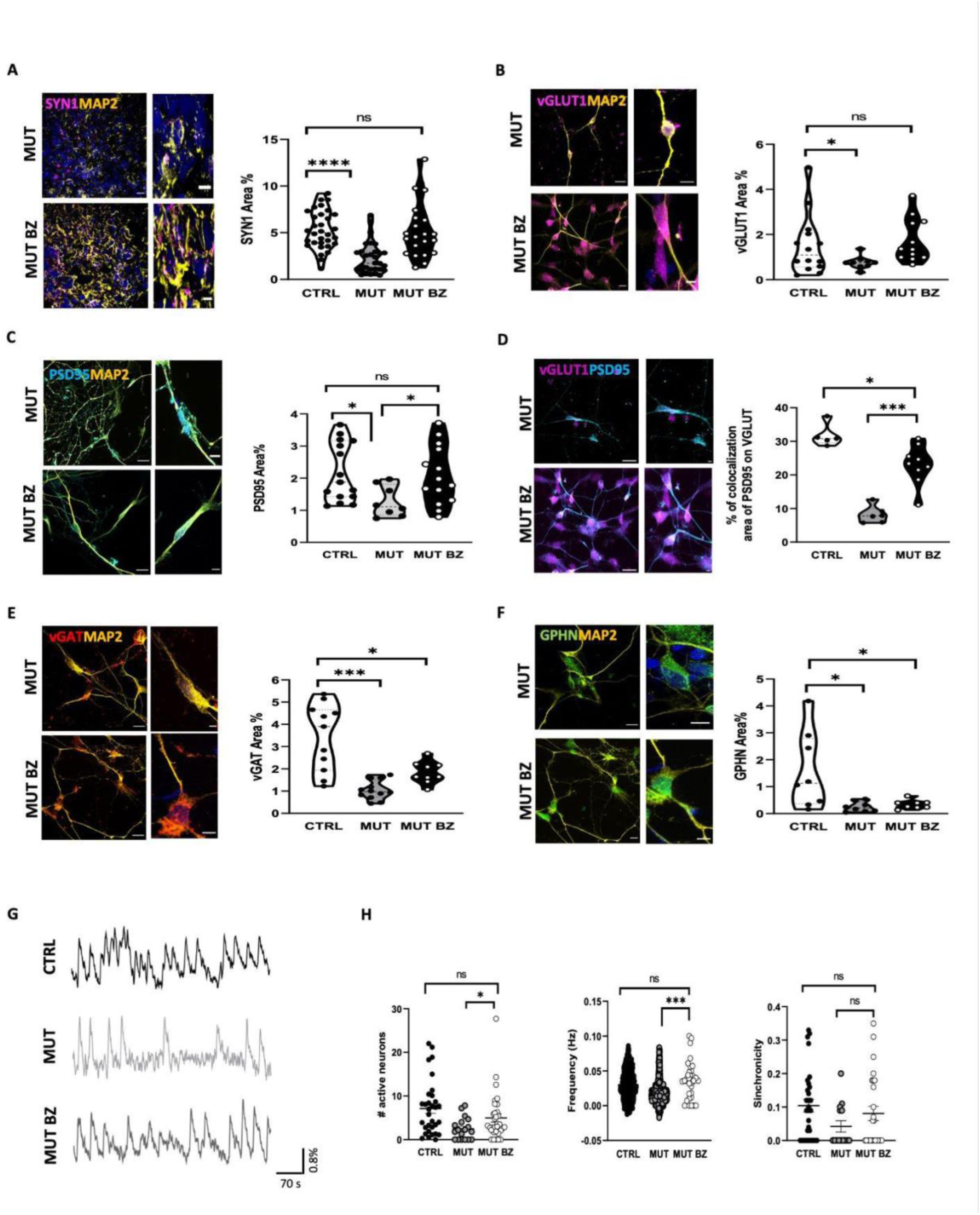
Bezafibrate treatment rescues synaptic functionality. **A)** Left, representative, MUT (top) and MUT BZ (bottom) derived cortical organoids immunolabeled for anti-Synapsin-1 (SYN1) (magenta), anti-MAP2 for neuronal cytoskeleton (yellow) and Hoechst (blue) for nuclei visualization at D100. Scale bar 30µm and 5µm. Right, violin plot representing SYN1 signal as % of the area covered within the field of view (FOV) (***p < 0.001 MW test, NS p>0.05; CTRL n = 32/3 FOVs/ batches; MUT n=30/4 FOVs/batches; MUT BZ n = 21/2 FOVs/ batches). **B)** Left: Representative images of MUT (top) and MUT BZ (bottom) cortical neurons immunolabeled for anti-VGluT1 (VGluT1) (magenta) and anti-MAP2 for neuronal cytoskeleton (yellow) at D100. Scale bar 30µm and 5µm. Right: violin plot representing VGluT1 signal as % of the area covered within the field of view (FOV) (MW test *p<0.05, **p<0.01, NS= 0.27; MUT n=7/2; MUT BZ n=12/2 FOVs/batches). **C)** Left, representative images of MUT (top) and MUT BZ (bottom) cortical neurons immunolabeled for anti-PSD95 (PSD95) (cyan) and anti-MAP2 for neuronal cytoskeleton (yellow) at D100. Scale bar 30µm and 5µm. Right: violin plot representing PSD95 signal as % of the area covered within the field of view (FOV) (*p < 0.05; NS= 0.88 MW test; MUT n = 10/2; MUT BZ n=14/2 FOVs/batches). **D)** Left, representative images of MUT (top) and MUT BZ (bottom) cortical neurons immunolabeled for PSD95 and VGluT1 at D100. Scale bar 30µm and 5µm. Right: violin plot representing VGluT1 and PSD95 colocalization signal as % of the area covered within the field of view (FOV) (*p<0.05; **p < 0.01 MW test; CTRL n = 5/2 FOVs/ batches; MUT n = 6/2 FOVs/ batches; MUT BZ n 9/3 FOVs/batches) **E)** Left, representative images of MUT (top) and MUT BZ (bottom) 2D cortical neurons immunolabeled for VGAT (red), anti-MAP2 for neuronal cytoskeleton (yellow) at D100. Scale bar 30µm. Right: violin plot representing VGAT signal as % of the area covered within the field of view (FOV) (***p < 0.0001; *p<0.05 MW test; CTRL n = 11/2 FOVs/ batches; MUT n = 12/2 FOVs/ batches; MUT BZ 9/2 FOVs/ batches). **F)** Left, representative images of MUT (top) and MUT BZ (bottom) 2D cortical neurons immunolabeled for Gepherin (green), anti-MAP2 for neuronal cytoskeleton (yellow) at D100. Scale bar 30µm. Right: violin plot representing Gepherin signal as % of the area covered within the field of view (FOV) (*p < 0.05 MW test; CTRL n = 8/2 FOVs/ batches; MUT n = 8/1 FOVs/ batches; MUT BZ n=8/2 batches). **G)** Representative traces of spontaneous calcium oscillations recorded CTRL (up), untreated tau-MUT (MUT, middle) and treated tau-MUT (MUT BZ, bottom) derived cortical organoids at D 100 of the differentiation protocol. **H)** Scatter dot plots representing the number of active neurons (left), the frequency (middle), and the synchronicity (right) of spontaneous calcium oscillations (*p<0.01, ***p<0.01, MW test; CTRL n= 47/ FOVs/ batches; MUT n= 29/ FOVs/ batches; MUT BZ n= 29/4 FOVs/batches).

Furthermore, BZ treatment enhanced network functionality within tau-mutant organoids. Calcium imaging recordings (Supplementary Movie3, Movie4) of spontaneous calcium transients in fast cells, most likely neurons (Figure 6G), showed that BZ treatment normalized the number of active neurons in tau-mutant organoids to levels observed in controls (Figure 6H). This normalization was not only quantitative but also qualitative, as the spontaneous activity patterns of active cells in BZ-treated tau-mutant organoids mirrored those in control organoids. Specifically, BZ treatment elevated the frequency (Figure 6H) and synchronicity (Figure 6H) of calcium events to match those seen in controls, while maintaining peak amplitude characteristics (Supplementary Figure 6).

These data indicate that BZ besides supporting cellular and mitochondrial health, restores neuronal network morphology and dynamics, suggesting a comprehensive therapeutic effect on tauopathy-impacted neural systems.

## 4 Discussion

Tauopathies, characterized by abnormal tau protein aggregation, represent a significant unmet clinical need due to their association with various neurodegenerative diseases, including AD and FTD. The pathogenic mechanisms underlying tauopathies, particularly the role of tau isoform dysregulation and tau hyperphosphorylation, remain incompletely understood. Developing effective treatments for tauopathies is challenging, suggesting that advanced in vitro disease models are needed to elucidate disease mechanisms and identify new therapeutic targets.

In this study, we found that the FTD-linked IVS 10+16 MAPT mutation on tau isoforms impaired the neuronal development and network functionality in 3D cortical organoids, and explored the therapeutic potential of bezafibrate to promote the mitochondrial biogenesis and counteract the effects of tau mutation using an in vitro 3D disease model.

Specifically, tau-mutant cortical organoids were characterized by: i) tau isoform imbalance and increased levels of hyperphosphorylated tau, with impaired neuronal integrity; ii) defective neuronal and glial maturation; iii) synaptic and network dysfunction; and iv) mitochondrial dysfunction. Treatment with bezafibrate was mainly effective in normalizing: i) mitochondrial parameters; ii) neuronal integrity and synaptic maturation, and iii) network functionality, highlighting its promise as a therapeutic strategy for tauopathies.

For the 3D cortical organoids used in this study to characterize the effects of the FTD-linked IVS 10+16 MAPT mutation on cortical homeostasis isogenic hiPSC lines were used. We modified an established protocol to obtain cortical organoids from human iPSCs (19)(Sloan et al., 2018), fostering organoid maturation by the supplementation of heparin, ascorbic acid, BDNF, and GDNF (21)(22) (Bai et al., 2021; Cortes et al., 2016). Maturation was assessed in the control line, by the upregulation of several genes (from D50 to D100) chiefly associated with synaptic transmission, protein binding, and acquisition of neuronal morphology, paralleled by the reduction of genes related to the cell cycle.

This disruption of neuronal differentiation and synaptic maturation, correlated with the intronic IVS 10+16 MAPT mutation, was evidenced by the downregulation of genes primarily associated with excitatory and inhibitory synapses, neuronal morphology, and nervous system development, alongside the upregulation of genes linked to cell proliferation and cellular stress. At the protein level, tau-mutant organoids showed increased levels of the neural progenitor marker PAX6 and delays in both neuronal and glial maturation. These organoids also exhibited fragmented neurite morphology, delayed specification of neuronal subtypes and GFAP expression and impaired synaptic formation and network functionality in whole cortical organoids.

Differential gene expression analysis at D 100 highlighted a significant downregulation of transcripts associated with both glutamatergic and GABAergic synapses and synaptic structures in tau-mutant organoids. Immunofluorescence analysis confirmed the reduced expression of pan synaptic markers synapsin1 and syngirin1, aligning with previous reports of downregulated synaptic maturation in other in vitro models of tauopathies (9)(Tracy et al., 2022). Furthermore, immunostaining of cortical neurons from dissociated organoids revealed reduced formation of both glutamatergic and GABAergic synapses. Specifically, lower levels of pre-synaptic (VGluT1) and post-synaptic (PSD95) proteins were observed for glutamatergic synapses, and reduced levels of pre-synaptic (VGAT) and post-synaptic (Gephyrin) proteins were noted for GABAergic synapses.

Consistently, in our model, calcium imaging of spontaneous transients in whole organoids revealed a reduced number of active neurons in tau-mutant organoids, characterized by weaker spontaneous activity. Both the frequency and synchronicity of calcium events were significantly lower in tau-mutant organoids compared to isogenic controls, indicating impaired network functionality.

These findings align with previous reports in 2D cortical neurons derived from the same iPSC lines, which exhibited impaired neuronal specification (6) (Verheyen et al., 2018), altered neuronal excitability, reduced firing, and decreased calcium transients (7)(30)(Kopach et al., 2021; Minaya et al., 2023). This underscores a causal link between the IVS 10+16 MAPT mutation and neuronal dysfunction.

Strikingly, our observations are also consistent with the recent examination of post-mortem brain tissue from people who died with FTD with tau pathology caused by the MAPT intronic exon 10+16 mutation. Indeed, bulk tissue RNA sequencing revealed substantial downregulation of gene expression associated with synaptic function, further confirmed by histopathological analysis of the patient’s cortex (31)(Dando et al., 2024).

In this study, we also found that the IVS 10+16 MAPT mutation caused a significant imbalance in tau isoforms, favoring 4R tau over 3R tau. This was evident from Real-Time PCR, immunofluorescence, and western blot analyses, indicating higher 4R tau expression in mutant organoids compared to controls, in line with data in 2D cortical cultures (6)(Vereyen et al., 2018). This was paralleled, in the organoid system, by elevated levels of hyperphosphorylated tau at Ser202/Thr205 (AT8) and Thr181, revealed by western blot and immunofluorescence analyses.

In line with this result, and with the impact of hyperphosphorylation in disrupting tau’s role in microtubule stabilization (32)(Parato et al., 2023), we reported a fragmented neurite morphology in tau-mutant organoids, underscoring the mutation’s impact on neuronal integrity and suggesting early-stage disruptions in neuronal development, likely contributing to the observed deficits in synaptic maturation and neuronal differentiation. Moreover, pathological tau, mislocalized into pre- and postsynaptic neuronal compartments, has been suggested to induce synaptic dysfunction (33)(Wu et al., 2021).

The observed impairment of cortical organoid maturation and function can be ascribed to the impact of tau protein abnormalities, including hyperphosphorylation and aggregation, on defective mitochondrial function, which is known to contribute to neuronal damage and disease progression in AD and FTD (34)(35)(Korn et al., 2022; Hope et al., 2023).

Indeed, on D100, we found 17 “tau signature” genes (9)(Tracy et al., 2022) differentially regulated in tau-mutant organoids, encompassing synaptic vesicles, cytoskeletal structure, and mitochondrial and lysosome functions. This points to a broad impairment of neuronal network development and mitochondrial functions.

Specifically, when looking at all mitochondrial function-related genes analyzed in the nanostring panel, we found 46 deregulated genes associated with autophagy, unfolded protein response, and oxidative stress (36)(37)(38)(39)(40)(41)(42)(Han et al., 2023; Patergnani et al., 2021; Cobley et al., 2018; Munoz-Carvajal et al., 2020; Duran-Aniotz et al., 2023; Tunold et al., 2023; Uddin et al., 2020).

We found impaired mitochondrial biogenesis and function in tau-mutant organoids, evidenced by fewer and smaller mitochondria. Moreover, the mitochondrial biogenesis master gene, PPARGC1A (43)(44)(Zhender et al., 2021; Kuczynska et al., 2021), was significantly downregulated in tau-mutant organoids at D100. This downregulation was observed at a critical period for cortical organoid development (D50-70), indicating the crucial role of PPARGC1A reduction in contributing to the observed deficits in mitochondrial quantity and size, which are crucial for sustaining neuronal energy demands and overall cellular health.

In turn, mitochondrial dysfunction has been shown to induce tau abnormality favoring tau phosphorylation and aggregation, further contributing to disease progression (12)(Cheng et al., 2018)

The relationship between impaired mitochondrial biogenesis and tau mutation suggests that targeting mitochondrial pathways could be a viable therapeutic strategy for treating tauopathies. Enhancing mitochondrial biogenesis and function could potentially alleviate some of the neurodegenerative processes associated with tau mutations, that contribute to synaptic impairment and cognitive decline in AD and FTD (11)(12)(Quantanilla et al., 2020; Cheng et al., 2018).

Our study also demonstrates the potential of bezafibrate, a peroxisome proliferator-activated receptor agonist, in mitigating mitochondrial dysfunction and enhancing neuronal integrity and function in tau-mutant cortical organoids, as demonstrated by the fact that BZ treatment effectively normalized mitochondrial parameters in tau-mutant cortical organoids, based on mitoTracker labeling and TUJ1 immunostaining. RT-PCR analysis showed that BZ treatment also restored to control level, key genes associated with oxidative stress, unfolded protein response, and autophagy in tau-mutant organoids, confirming the role of BZ in enhancing mitochondrial function, previously reported in human iPSC-derived neural progenitor cells (45)(Augustyniak et al., 2019).

The enhanced neuronal differentiation and maturation in tau-mutant cortical organoids induced by BZ treatment was evidenced by the reduction of PAX6 expression to control levels, the decrease in the number of MAP2 fragments, and the increase in their length, indicating enhanced neuronal differentiation, maturation, and neurite integrity. Consistently, BZ treatment also restored synaptic connectivity and network functionality. The enhanced expression of synaptic markers and improved calcium signaling patterns in BZ-treated tau-mutant organoids underscore its ability to restore synaptic integrity and neuronal connectivity.

The rescue of neuronal homeostasis and morphogenesis aligns with previous findings in an in vitro disease model of Leigh Syndrome, a rare and severe neurometabolic disorder (46)(Inak et et.., 2021), and in a mouse model of Huntington’s Disease (47)(Johri et al., 2012). Importantly, BZ treatment not only improved neuronal maturation but also restored the expression levels of the 4R tau isoform and reduced phosphorylated tau to control levels.

A relationship between tau phosphorylation and cortical neuron development has been reported in rats, showing that fibrillary tau and fetal tau share several phosphorylated sites (48)(Watanabe et al., 1993) and that during neurite stabilization and synaptogenesis tau phosphorylation decreases (49)(Brion et al., 1994).

Therefore, the observed effects of BZ on neuronal maturation and synaptic activity can be interpreted as a consequence of the reduction of tau phosphorylation, which is likely mediated by the improved mitochondrial biogenesis and function (12)(Chen et al., 2018), highlighting the interconnection between these cellular processes.

Our findings may explain the mechanisms underlying previous results observed in rodent models of AD and FTD. Specifically, in a rat model of sporadic AD induced by streptozotocin, BZ treatment significantly improved cognitive function, attenuated tau pathology, reduced neuronal loss, and decreased neuroinflammation (29)(Lin et al., 2022), while in P301S transgenic mice, a model of familial FTD with tau pathology, BZ administration decreased tau hyperphosphorylation, reduced microglial activation, and improved behavioral deficits (28) (Dumont et al., 2012).

The findings from our study, along with others, suggest that BZ adopts a multi-targeted approach in treating AD and tauopathies, making it a promising candidate for future therapeutic development. BZ improves neuronal differentiation and maturation, restores synaptic connectivity, and reduces pathological tau phosphorylation, potentially through enhanced mitochondrial function.

Collectively, our findings suggest that bezafibrate could be a promising therapeutic strategy for tauopathies by targeting mitochondrial dysfunction and synaptic deficits. By enhancing mitochondrial health and promoting neuronal maturation, BZ may mitigate some of the key pathological features of tau-related neurodegeneration. These results underscore the potential of bezafibrate as a comprehensive therapeutic strategy for addressing the multifaceted pathology of tauopathies.

Our 3D in vitro disease model has proven to be a valuable tool for studying tauopathies and testing new drugs, as well as for drug repurposing. The model’s ability to mimic key aspects of tau-related neurodegeneration makes it an excellent platform for evaluating the therapeutic potential of compounds like bezafibrate. However, several broader questions persist. All in vitro iPSC-derived models are confined to an early developmental stage, and the applicability of these findings to other MAPT mutations and sporadic tauopathies remains uncertain. Moreover, including microglia in the model would be beneficial, given its role in tau pathology, propagation, and synaptic degeneration (50)(15)(Vogels et al., 2019; D’Antoni et al., 2023).

## AUTHOR CONTRIBUTION

FC performed organoids generation experiments, molecular, cellular and functional characterization; LM performed molecular and immunofluorescence analysis on organoids; SG wrote the MATLAB code and performed quantification of confocal and calcium imaging acquisitions; DS performed and anayzed nanostring experiments, MAR and MC designed, performed and analyzed western blot experiments; PB and SDA acquired funds and supervised the project; SDA conceived the study, designed the experiments, interpreted the results, and wrote the manuscript with the help of FC. All authors have read and agreed to the published version of the manuscript.

## ACKNOWLEDGMENTS

The authors wish to thank the Imaging Facility of the Center for Life Nano- and Neuro-Science Imaging, Istituto Italiano di Tecnologia. This work was supported by MUR PRIN 2022 (CUP: 2022CFP7RF, to SDA). This research was also funded by the D-Tails-IIT Joint Lab (to SDA), the Regione Lazio FSE 2014–2020 (19036AP000000019 and A0112E0073) grants (to SDA), Sapienza University grants (RM118163E0297F84, PH12017270934C3C, and MA32117A7B698029 to SDA), and Fondazione Istituto Italiano di Tecnologia (to LM). LM was also supported by the PhD program in Life Science at Sapienza University in Rome. SDA was also supported by Progetto ECS 0000024 Rome Technopole, - CUP B83C22002820006, PNRR Missione 4 Componente 2 Investimento 1.5, finanziato dall’Unione europea – NextGenerationEU. SDA was also supported by Alternative Methods to Animal Testing Grant 2023 (NEURO-3R).

## CONFLICT OF INTEREST STATEMENT

The funders had no role in the design of the study; in the collection, analysis, or interpretation of data; in the writing of the manuscript; or in the decision to publish the results.

SDA is a scientific advisor of D-Tails s.r.l. The remaining authors declare that the research was conducted in the absence of any commercial or financial relationships that could be construed as a potential conflict of interest.

## CONSENT STATEMENT

Consent from human subjects was not necessary.

## REFERENCES

1. Antonioni A, Raho EM, Lopriore P, Pace AP, Latino RR, Assogna M, Mancuso M, Gragnaniello D, Granieri E, Pugliatti M, Di Lorenzo F, Koch G. Frontotemporal Dementia, Where Do We Stand? A Narrative Review. Int J Mol Sci. 2023 Jul 21;24(14):11732. doi: 10.3390/ijms241411732. PMID: 37511491; PMCID: PMC10380352.

2. Chang CW, Shao E, Mucke L. Tau: Enabler of diverse brain disorders and target of rapidly evolving therapeutic strategies. Science. 2021 Feb 26;371(6532):eabb8255. doi: 10.1126/science.abb8255. PMID: 33632820; PMCID: PMC8118650.

3. Barbier P, Zejneli O, Martinho M, Lasorsa A, Belle V, Smet-Nocca C, Tsvetkov PO, Devred F, Landrieu I. Role of Tau as a Microtubule-Associated Protein: Structural and Functional Aspects. Front Aging Neurosci. 2019 Aug 7;11:204. doi: 10.3389/fnagi.2019.00204. PMID: 31447664; PMCID: PMC6692637.

4. Leveille E, Ross OA, Gan-Or Z. Tau and MAPT genetics in tauopathies and synucleinopathies. Parkinsonism Relat Disord. 2021 Sep;90:142–154. doi: 10.1016/j.parkreldis.2021.09.008. Epub 2021 Sep 14. PMID: 34593302; PMCID: PMC9310195.

5. Fujioka S, Wszolek ZK. Clinical aspects of familial forms of frontotemporal dementia associated with parkinsonism. J Mol Neurosci. 2011 Nov;45(3):359–65. doi: 10.1007/s12031011-9568-5. Epub 2011 Jun 8. PMID: 21656039; PMCID: PMC3909923.

6. Verheyen A, Diels A, Reumers J, Van Hoorde K, Van den Wyngaert I, van Outryve d’Ydewalle C, De Bondt A, Kuijlaars J, De Muynck L, De Hoogt R, Bretteville A, Jaensch S, Buist A, Cabrera-Socorro A, Wray S, Ebneth A, Roevens P, Royaux I, Peeters PJ. Genetically Engineered iPSC-Derived FTDP-17 MAPT Neurons Display Mutation-Specific Neurodegenerative and Neurodevelopmental Phenotypes. Stem Cell Reports. 2018 Aug 14;11(2):363–379. doi: 10.1016/j.stemcr.2018.06.022. Epub 2018 Jul 26. Erratum in: Stem Cell Reports. 2019 Aug 13;13(2):434-435. doi: 10.1016/j.stemcr.2019.07.007. PMID: 30057263; PMCID: PMC6093179.

7. Kopach, O., Esteras, N., Wray, S. et al. Genetically engineered MAPT 10+16 mutation causes pathophysiological excitability of human iPSC-derived neurons related to 4R tau-induced dementia. Cell Death Dis 12, 716 (2021). 10.1038/s41419-021-04007-w

8. Declan, James, Peter, King., Tara, L., Spires-Jones. (2022). Tau talk – synaptic and mitochondrial proteins interact with Tau in human neurons. Trends in Neurosciences, doi: 10.1016/j.tins.2022.02.004

9. Tracy TE, Madero-Pérez J, Swaney DL, Chang TS, Moritz M, Konrad C, Ward ME, Stevenson E, Hüttenhain R, Kauwe G, Mercedes M, Sweetland-Martin L, Chen X, Mok SA, Wong MY, Telpoukhovskaia M, Min SW, Wang C, Sohn PD, Martin J, Zhou Y, Luo W, Trojanowski JQ, Lee VMY, Gong S, Manfredi G, Coppola G, Krogan NJ, Geschwind DH, Gan L. Tau interactome maps synaptic and mitochondrial processes associated with neurodegeneration. Cell. 2022 Feb 17;185(4):712–728.e14. doi: 10.1016/j.cell.2021.12.041. Epub 2022 Jan 20. PMID: 35063084; PMCID: PMC8857049.

10. Szabo L, Eckert A, Grimm A. Insights into Disease-Associated Tau Impact on Mitochondria. Int J Mol Sci. 2020 Sep 1;21(17):6344. doi: 10.3390/ijms21176344. PMID: 32882957; PMCID: PMC7503371.

11. Quntanilla RA, Tapia-Monsalves C. The Role of Mitochondrial Impairment in Alzheimeŕs Disease Neurodegeneration: The Tau Connection. Curr Neuropharmacol. 2020;18(11):1076–1091. doi: 10.2174/1570159X18666200525020259. PMID: 32448104; PMCID: PMC7709157.

12. Cheng Y, Bai F. The Association of Tau With Mitochondrial Dysfunction in Alzheimer’s Disease. Front Neurosci. 2018 Mar 22;12:163. doi: 10.3389/fnins.2018.00163. PMID: 29623026; PMCID: PMC5874499.

13. Cordella F, Brighi C, Soloperto A, Di Angelantonio S. Stem cell-based 3D brain organoids for mimicking, investigating, and challenging Alzheimer’s diseases. Neural Regen Res. 2022 Feb;17(2):330–332. doi: 10.4103/1673-5374.317976. PMID: 34269204; PMCID: PMC8463991.

14. Brighi C, Cordella F, Chiriatti L, Soloperto A, Di Angelantonio S. Retinal and Brain Organoids: Bridging the Gap Between in vivo Physiology and in vitro Micro-Physiology for the Study of Alzheimer’s Diseases. Front Neurosci. 2020 Jun 17;14:655. doi: 10.3389/fnins.2020.00655. PMID: 32625060; PMCID: PMC7311765.

15. D’Antoni C, Mautone L, Sanchini C, Tondo L, Grassmann G, Cidonio G, Bezzi P, Cordella F, Di Angelantonio S. Unlocking Neural Function with 3D In Vitro Models: A Technical Review of Self-Assembled, Guided, and Bioprinted Brain Organoids and Their Applications in the Study of Neurodevelopmental and Neurodegenerative Disorders. Int J Mol Sci. 2023 Jun 28;24(13):10762. doi: 10.3390/ijms241310762. PMID: 37445940; PMCID: PMC10341866.

16. Wegiel J, Flory M, Kuchna I, Nowicki K, Wegiel J, Ma SY, Zhong N, Bobrowicz TW, de Leon M, Lai F, Silverman WP, Wisniewski T. Developmental deficits and staging of dynamics of age associated Alzheimer’s disease neurodegeneration and neuronal loss in subjects with Down syndrome. Acta Neuropathol Commun. 2022 Jan 4;10(1):2. doi: 10.1186/s40478-021-01300-9. PMID: 34983655; PMCID: PMC8728914.

17. Brighi C, Salaris F, Soloperto A, Cordella F, Ghirga S, de Turris V, Rosito M, Porceddu PF, D’Antoni C, Reggiani A, Rosa A, Di Angelantonio S. Novel fragile X syndrome 2D and 3D brain models based on human isogenic FMRP-KO iPSCs. Cell Death Dis. 2021 May 15;12(5):498. doi: 10.1038/s41419-021-03776-8. PMID: 33993189; PMCID: PMC8124071.

18. Cordella F, Ferrucci L, D’Antoni C, Ghirga S, Brighi C, Soloperto A, Gigante Y, Ragozzino D, Bezzi P, Di Angelantonio S. Human iPSC-Derived Cortical Neurons Display Homeostatic Plasticity. Life (Basel). 2022 Nov 14;12(11):1884. doi: 10.3390/life12111884. PMID: 36431019; PMCID: PMC9696876.

19. Sloan SA, Andersen J, Pașca AM, Birey F, Pașca SP. Generation and assembly of human brain region-specific three-dimensional cultures. Nat Protoc. 2018 Sep;13(9):2062–2085. doi: 10.1038/s41596-018-0032-7. PMID: 30202107; PMCID: PMC6597009.

20. Sakaguchi H, Ozaki Y, Ashida T, Matsubara T, Oishi N, Kihara S, Takahashi J. Self-Organized Synchronous Calcium Transients in a Cultured Human Neural Network Derived from Cerebral Organoids. Stem Cell Reports. 2019 Sep 10;13(3):458–473. doi: 10.1016/j.stemcr.2019.05.029. Epub 2019 Jun 27. PMID: 31257131; PMCID: PMC6739638.

21. Bai, R., Chang, Y., Saleem, A. et al. Ascorbic acid can promote the generation and expansion of neuroepithelial-like stem cells derived from hiPS/ES cells under chemically defined conditions by promoting collagen synthesis. Stem Cell Res Ther 12, 48 (2021). 10.1186/s13287-020-02115-6

22. Cortés D, Robledo-Arratia Y, Hernández-Martínez R, Escobedo-Ávila I, Bargas J, Velasco I. Transgenic GDNF Positively Influences Proliferation, Differentiation, Maturation and Survival of Motor Neurons Produced from Mouse Embryonic Stem Cells. Front Cell Neurosci. 2016 Sep 12;10:217. doi: 10.3389/fncel.2016.00217. PMID: 27672361; PMCID: PMC5018488.

23. Young AL, Bocchetta M, Russell LL, Convery RS, Peakman G, Todd E, Cash DM, Greaves CV, van Swieten J, Jiskoot L, Seelaar H, Moreno F, Sanchez-Valle R, Borroni B, Laforce R Jr, Masellis M, Tartaglia MC, Graff C, Galimberti D, Rowe JB, Finger E, Synofzik M, Vandenberghe R, de Mendonça A, Tagliavini F, Santana I, Ducharme S, Butler C, Gerhard A, Levin J, Danek A, Otto M, Sorbi S, Williams SCR, Alexander DC, Rohrer JD; Genetic FTD Initiative (GENFI). Characterizing the Clinical Features and Atrophy Patterns of MAPT-Related Frontotemporal Dementia With Disease Progression Modeling. Neurology. 2021 Aug 31;97(9):e941–e952. doi: 10.1212/WNL.0000000000012410. Epub 2021 Jun 22. PMID: 34158384; PMCID: PMC8408507.

24. Wang Y, Mandelkow E. Tau in physiology and pathology. Nat Rev Neurosci. 2016 Jan;17(1):5–21. doi: 10.1038/nrn.2015.1. Epub 2015 Dec 3. PMID: 26631930.

25. Nataliya, I., Trushina., Lidia, Bakota., Armen, Y., Mulkidjanian., Roland, Brandt. (2019). The Evolution of Tau Phosphorylation and Interactions.. Frontiers in Aging Neuroscience, doi: 10.3389/FNAGI.2019.00256

26. Man VH, He X, Gao J, Wang J. Phosphorylation of Tau R2 Repeat Destabilizes Its Binding to Microtubules: A Molecular Dynamics Simulation Study. ACS Chem Neurosci. 2023 Feb 1;14(3):458–467. doi: 10.1021/acschemneuro.2c00611. Epub 2023 Jan 20. PMID: 36669127; PMCID: PMC10032563.

27. Mueller RL, Combs B, Alhadidy MM, Brady ST, Morfini GA, Kanaan NM. Tau: A Signaling Hub Protein. Front Mol Neurosci. 2021 Mar 19;14:647054. doi: 10.3389/fnmol.2021.647054. PMID: 33815057; PMCID: PMC8017207.

28. Dumont M, Stack C, Elipenahli C, Jainuddin S, Gerges M, Starkova N, Calingasan NY, Yang L, Tampellini D, Starkov AA, Chan RB, Di Paolo G, Pujol A, Beal MF. Bezafibrate administration improves behavioral deficits and tau pathology in P301S mice. Hum Mol Genet. 2012 Dec 1;21(23):5091–105. doi: 10.1093/hmg/dds355. Epub 2012 Aug 24. PMID: 22922230; PMCID: PMC3490516.

29. Lin LF, Jhao YT, Chiu CH, Sun LH, Chou TK, Shiue CY, Cheng CY, Ma KH. Bezafibrate Exerts Neuroprotective Effects in a Rat Model of Sporadic Alzheimer’s Disease. Pharmaceuticals (Basel). 2022 Jan 18;15(2):109. doi: 10.3390/ph15020109. PMID: 35215222; PMCID: PMC8877080.

30. Minaya MA, Mahali S, Iyer AK, Eteleeb AM, Martinez R, Huang G, Budde J, Temple S, Nana AL, Seeley WW, Spina S, Grinberg LT, Harari O, Karch CM. Conserved gene signatures shared among MAPT mutations reveal defects in calcium signaling. Front Mol Biosci. 2023 Feb 9;10:1051494. doi: 10.3389/fmolb.2023.1051494. PMID: 36845551; PMCID: PMC9948093.

31. Dando O, McGeachan R, McQueen J, Baxter P, Rockley N, McAlister H, Prasad A, He X, King D, Rose J, Jones PB, Tulloch J, Chandran S, Smith C, Hardingham G, Spires-Jones TL. Synaptic gene expression changes in frontotemporal dementia due to the MAPT 10+16 mutation. medRxiv [Preprint]. 2024 Apr 12:2024.04.09.24305501. doi: 10.1101/2024.04.09.24305501. PMID: 38645146; PMCID: PMC11030522.

32. ParatoJ, Qu X, Richters E, Sproul A, Bartolini F. Impaired Microtubule Dynamics Are a Driver of Tau Phosphorylation. 2023 doi: 10.1002/alz.080227

33. Wu M, Zhang M, Yin X, Chen K, Hu Z, Zhou Q, Cao X, Chen Z, Liu D. The role of pathological tau in synaptic dysfunction in Alzheimer’s diseases. Transl Neurodegener. 2021 Nov 10;10(1):45. doi: 10.1186/s40035-021-00270-1. PMID: 34753506; PMCID: PMC8579533.

34. Korn L, Speicher AM, Schroeter CB, Gola L, Kaehne T, Engler A, Disse P, Fernández-Orth J, Csatári J, Naumann M, Seebohm G, Meuth SG, Schöler HR, Wiendl H, Kovac S, Pawlowski M. MAPT genotype-dependent mitochondrial aberration and ROS production trigger dysfunction and death in cortical neurons of patients with hereditary FTLD. Redox Biol. 2023 Feb;59:102597. doi: 10.1016/j.redox.2022.102597. Epub 2022 Dec 30. PMID: 36599286; PMCID: PMC9817175.

35. Hope, I., Needs., Kevin, A., Wilkinson., Jeremy, M., Henley., Ian, Collinson. (2023). Aggregation-prone Tau impairs mitochondrial import, which affects organelle morphology and neuronal complexity. Journal of Cell Science, doi: 10.1242/jcs.260993

36. Han R, Liu Y, Li S, Li XJ, Yang W. PINK1-PRKN mediated mitophagy: differences between in vitro and in vivo models. Autophagy. 2023 May;19(5):1396–1405. doi: 10.1080/15548627.2022.2139080. Epub 2022 Nov 3. PMID: 36282767; PMCID: PMC10240983.

37. 37. 37 Patergnani et al., 2021; Patergnani S, Bouhamida E, Leo S, Pinton P, Rimessi A. Mitochondrial Oxidative Stress and “Mito-Inflammation”: Actors in the Diseases. Biomedicines. 2021 Feb 20;9(2):216. doi: 10.3390/biomedicines9020216. PMID: 33672477; PMCID: PMC7923430.

38. Cobley JN, Fiorello ML, Bailey DM. 13 reasons why the brain is susceptible to oxidative stress. Redox Biol. 2018 May;15:490–503. doi: 10.1016/j.redox.2018.01.008. Epub 2018 Feb 3. PMID: 29413961; PMCID: PMC5881419.

39. Muñoz-Carvajal F, Sanhueza M. The Mitochondrial Unfolded Protein Response: A Hinge Between Healthy and Pathological Aging. Front Aging Neurosci. 2020 Sep 11;12:581849. doi: 10.3389/fnagi.2020.581849. PMID: 33061907; PMCID: PMC7518384

40. Duran-Aniotz C, Poblete N, Rivera-Krstulovic C, Ardiles ÁO, Díaz-Hung ML, Tamburini G, Sabusap CMP, Gerakis Y, Cabral-Miranda F, Diaz J, Fuentealba M, Arriagada D, Muñoz E, Espinoza S, Martinez G, Quiroz G, Sardi P, Medinas DB, Contreras D, Piña R, Lourenco MV, Ribeiro FC, Ferreira ST, Rozas C, Morales B, Plate L, Gonzalez-Billault C, Palacios AG, Hetz C. The unfolded protein response transcription factor XBP1s ameliorates Alzheimer’s disease by improving synaptic function and proteostasis. Mol Ther. 2023 Jul 5;31(7):2240–2256. doi: 10.1016/j.ymthe.2023.03.028. Epub 2023 Apr 4. PMID: 37016577; PMCID: PMC10362463..

41. Tunold JA, Tan MMX, Koga S, Geut H, Rozemuller AJM, Valentino R, Sekiya H, Martin NB, Heckman MG, Bras J, Guerreiro R, Dickson DW, Toft M, van de Berg WDJ, Ross OA, Pihlstrøm L. Lysosomal polygenic risk is associated with the severity of neuropathology in Lewy body disease. Brain. 2023 Oct 3;146(10):4077–4087. doi: 10.1093/brain/awad183. PMID: 37247383; PMCID: PMC10545498.

42. Uddin MS, Stachowiak A, Mamun AA, Tzvetkov NT, Takeda S, Atanasov AG, Bergantin LB, Abdel-Daim MM, Stankiewicz AM. Autophagy and Alzheimer’s Disease: From Molecular Mechanisms to Therapeutic Implications. Front Aging Neurosci. 2018 Jan 30;10:04. doi: 10.3389/fnagi.2018.00004. PMID: 29441009; PMCID: PMC5797541.

43. Zehnder T, Petrelli F, Romanos J, De Oliveira Figueiredo EC, Lewis TL Jr, Déglon N, Polleux F, Santello M, Bezzi P. Mitochondrial biogenesis in developing astrocytes regulates astrocyte maturation and synapse formation. Cell Rep. 2021 Apr 13;35(2):108952. doi: 10.1016/j.celrep.2021.108952. PMID: 33852851.

44. Kuczynska Z, Metin E, Liput M, Buzanska L. Covering the Role of PGC-1α in the Nervous System. Cells. 2021 Dec 30;11(1):111. doi: 10.3390/cells11010111. PMID: 35011673; PMCID: PMC8750669.

45. Augustyniak J, Lenart J, Gaj P, Kolanowska M, Jazdzewski K, Stepien PP, Buzanska L. Bezafibrate Upregulates Mitochondrial Biogenesis and Influence Neural Differentiation of Human-Induced Pluripotent Stem Cells. Mol Neurobiol. 2019 Jun;56(6):4346–4363. doi: 10.1007/s12035-0181368-2. Epub 2018 Oct 13. PMID: 30315479; PMCID: PMC6505510.

46. Inak G, Rybak-Wolf A, Lisowski P, Pentimalli TM, Jüttner R, Glažar P, Uppal K, Bottani E, Brunetti D, Secker C, Zink A, Meierhofer D, Henke MT, Dey M, Ciptasari U, Mlody B, Hahn T, Berruezo-Llacuna M, Karaiskos N, Di Virgilio M, Mayr JA, Wortmann SB, Priller J, Gotthardt M, Jones DP, Mayatepek E, Stenzel W, Diecke S, Kühn R, Wanker EE, Rajewsky N, Schuelke M, Prigione A. Defective metabolic programming impairs early neuronal morphogenesis in neural cultures and an organoid model of Leigh syndrome. Nat Commun. 2021 Mar 26;12(1):1929. doi: 10.1038/s41467-021-22117-z. PMID: 33771987; PMCID: PMC7997884.

47. Johri A, Calingasan NY, Hennessey TM, Sharma A, Yang L, Wille E, Chandra A, Beal MF. Pharmacologic activation of mitochondrial biogenesis exerts widespread beneficial effects in a transgenic mouse model of Huntington’s disease. Hum Mol Genet. 2012 Mar 1;21(5):1124–37. doi: 10.1093/hmg/ddr541. Epub 2011 Nov 17. PMID: 22095692; PMCID: PMC3277311.

48. Watanabe A, Hasegawa M, Suzuki M, Takio K, Morishima-Kawashima M, Titani K, Arai T, Kosik KS, Ihara Y. In vivo phosphorylation sites in fetal and adult rat tau. J Biol Chem. 1993 Dec 5;268(34):25712–7. PMID: 8245007.

49. Brion JP, Octave JN, Couck AM. Distribution of the phosphorylated microtubule-associated protein tau in developing cortical neurons. Neuroscience. 1994 Dec;63(3):895–909. doi: 10.1016/0306-4522(94)90533-9. PMID: 7898684.

50. Vogels T, Murgoci AN, and Hromádka T (2019). Intersection of pathological tau and microglia at the synapse. Acta Neuropathol. Commun 7, 109.

