## Supplementary Figures and Tables for "Bezafibrate treatment rescues neurodevelopmental and neurodegenerative defects in 3D cortical organoid model of MAPT frontotemporal dementia"

**Cordella et al.**

### SUPPORTING INFORMATION

**Supplementary Table 1: primary antibodies.**

| | | |
|---|---|---|
| *MAP2* | ABCAM – ab5392 | Chicken |
| *B-Tubulin III* | Sigma Aldrich – T2200 | Rabbit |
| *PAX6* | Invitrogen – sc81649 | Mouse |
| *GFAP* | Synaptic System - 173006 | Chicken |
| *TBR1* | ProteinTech – 20932-1-AP | Rabbit |
| *CTIP2* | ABCAM – ab18465 | Rat |
| *Synapsin1* | Cell Signaling – D12G5 | Rabbit |
| *Synaptogyrin 1* | Invitrogen – PA5- 56226 | Rabbit |
| *vGLUT1* | Sigma Aldrich – SAB5200258 | Mouse |
| *PSD95* | Cell Signaling - 3450 | Rabbit |
| *vGAT* | Synaptic System - 131003 | Rabbit |
| *Gephyrin* | Synaptic System - 147021 | Mouse |
| *AT8* | Invitrogen – MN1020 | Mouse |
| *4R Tau* | Biolegend – MMS-5020 | Mouse |
| *pTau 181* | MediMABS – MM0194 | Mouse |

**Supplementary Table 2: secondary antibodies**.

| | | | |
|---|---|---|---|
| Alexa fluor 488 | Goat | Anti-mouse | A11001 |
| Alexa fluor 488 | Goat | Anti-chicken | A11039 |
| Alexa fluor 594 | Goat | Anti-mouse | A11032 |
| Alexafluor 594 | Goat | Anti-rat | A11007 |
| Alexa fluor 647 | Goat | Anti-rabbit | A32733 |
| Hoechst | | 94403 | Sigma Aldrich |

### Supplementary Table 3: Primers for RT-PCR

| *Primers* | *Forward* | *Reverse* |
|---|---|---|
| NANOG | AGA ACA TGT GTA AGC TGC GG | GTT GCC TCT CAC TCG GTT C |
| PAX6 | GCC CTC ACA AAC ACC TAC AG | TCA TAA CTC CGC CCA TTC AC |
| TBR1 | GGA GCT TCA AAT AAC AAT GGG C | GAG TCT CAG GGA AAG TGA ACG |
| CTIP2 | GTT GTG CAA ATG TAG CTG GAA | GAA GAT GAC CAC CTG CTC TC |
| GFAP | GAT CAA CTC ACC GCC AAC AG | ATA GGC AGC CAG GTT GTT CT |
| PPARGC1A | CCA GAG TCA CCA AAT GAC CC | CCA CAG TCT TGC AAG AGG ACT |
| GS | TGG GAG CAG ACA GAG CCT AT | TCC CAG GAA TGG GCT TAG GA |
| SYN1 | CCC CAA TCA CAA AGA AAT GCT C | ATG TCC TGG AAG TCA TGC TG |
| GLUR1 | TGA TGG AAA ATA CGG AGC CC | CTT CCC GGA CCA AAG TGA TAG |
| NR1 | GAG AAG GAG AAC ATC ACC GAC | GTC CCC ATC CTC ATT GAA CTC |
| GABRA2 | TTA GCC AGC ACC AAC CTG | TCG TCA AGA TCA GGG CAA AAG |
| XBP1 | CCC TCC AGA ACA TCT CCC CAT | ACAT GAC TGG GTC CAA GTT GT |
| PINK1 | CCC AAG CAA CTA GCC CCT C | GGC AGC ACA TCA GGG TAG TC |
| CAST | CAA GCC GGG TGA CAA GAA AAA | CCC GAT GGT TTA TCC GGT TTA G |
| PCNA | CCT GCT GGG ATA TTA GCT CCA | CAG CGG TAG GTG TCG AAG C |
| CCNA2 | CGC TGG CGG TAC TGA AGT C | GAG GAA CGG TGA CAT GCT CAT |
| MAPT 4R | GCT CCA CTG AGA ACC TGA AG | TTG AGC CAC ACT TGG ACT G |
| MAPT 3R | GCT CCA CTG AGA ACC TGA AG | CCT AAT GAG CCA CAC TTG GA |

### Supplementary Table 4

Supplementary Figure 1

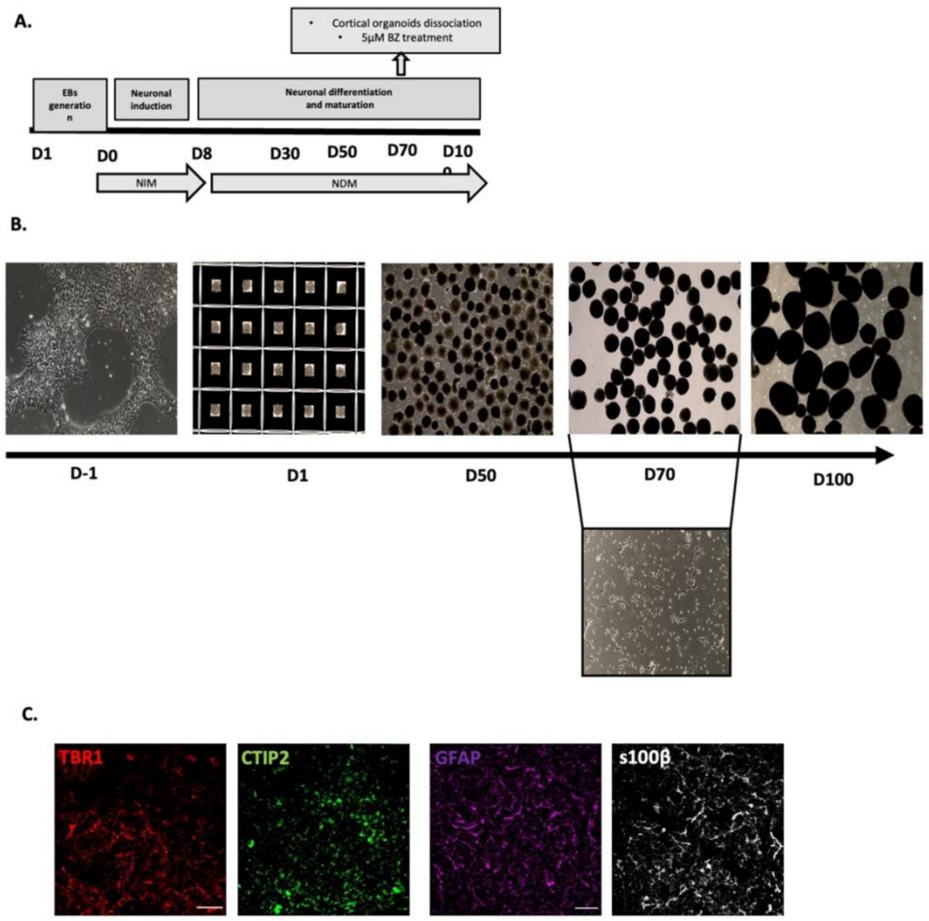


**Figure S1**.

A. Representative scheme of differentiation protocol for hIPSC-derived cortical organoids and organoid dissociation at D70.

B. Representative images of CTRL-derived cortical organoids development from D-1 to D100 of the differentiation protocol and 2D cortical colture derived from organoid dissociation at D70

C. Representative images of CTRL-derived cortical organoids at D100 immunolabeled for neuronal and astrocytic markers. TBR1 (red), CTIP2 (green) (scale bar 30um); GFAP (magenta), S100b (grey) (Scale bar 30um).

Supplementary Figure 2

**A**

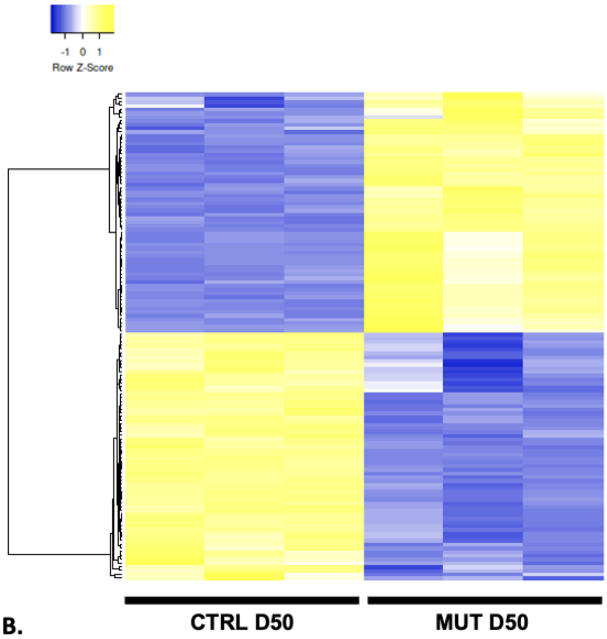


**B.**

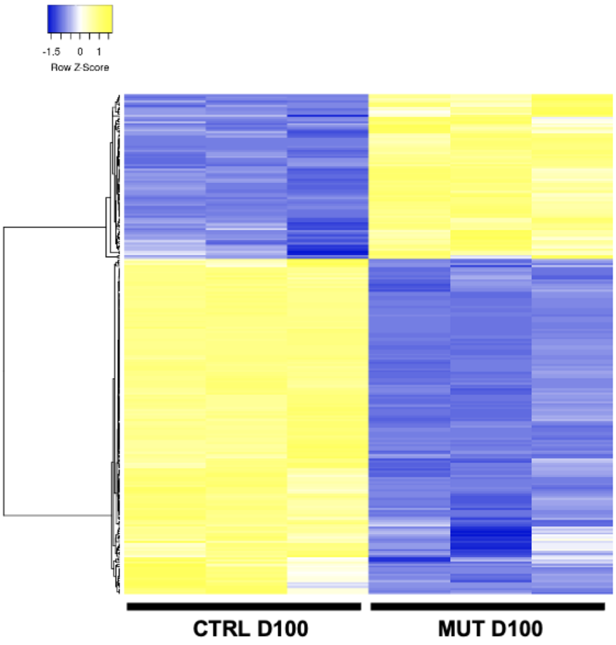


**Figure S2.** Heat map of supervised hierarchical clustering of **(A)** 130 differentially expressed (DE) genes between D50 CTRL and D50 MUT (64 genes up- and 66 genes down-regulated in D50 MUT) and **(B)** 243 DE genes between D100 CTRL and D100 MUT (80 genes up- and 163 genes down-regulated in D50 MUT). Average linkege and Pearson Distance Measurement Methods was performed for Hierarchical cluster using Heatmapper tools (http://www.heatmapper.ca) . The significant DE genes are selected by multiple t-test performed with Graph Pad Prism 6, with *pvalue* $\leq$ 0.05 and a fold change cut-off values of $\geq$1.5 or $\leq$0.66 for upmodulated and downregulated genes respectively.

Supplementary Table 4

| NAME | IPS d50 _1 | IPS d50 _2 | IPS d50 _3 | IVS d50 _1 | IVS d50 _2 | IVS d50 _3 |
|---|---|---|---|---|---|---|
| ABAT | 3857.39 | 3844.04 | 3785.12 | 2652.00 | 2240.60 | 2262.43 |
| ACHE | 1275.57 | 1264.85 | 1250.12 | 650.52 | 684.25 | 638.12 |
| ADCYAP1 | 137.12 | 94.14 | 129.80 | 25.61 | 19.82 | 22.64 |
| AKT3 | 2878.61 | 2825.40 | 2769.11 | 6357.90 | 6491.78 | 6394.66 |
| APOE | 39.69 | 38.62 | 37.09 | 334.22 | 128.42 | 199.50 |
| ARHGEF10 | 147.94 | 167.76 | 152.98 | 290.68 | 306.67 | 285.10 |
| ATP6V0D1 | 3028.36 | 2748.16 | 2728.16 | 2028.38 | 1435.59 | 1606.62 |
| ATP8A2 | 1990.04 | 2102.46 | 2121.64 | 1349.69 | 946.84 | 1166.59 |
| B4GALT6 | 609.82 | 597.43 | 586.43 | 1061.57 | 906.59 | 959.31 |
| C6 | 17.14 | 14.69 | 19.65 | 43.54 | 42.17 | 53.06 |
| C9orf72 | 1032.91 | 1123.64 | 1135.77 | 1703.12 | 1730.76 | 1837.25 |
| CALB1 | 1403.67 | 1218.99 | 1162.81 | 808.02 | 170.58 | 409.61 |
| CAMK2D | 1843.90 | 2033.66 | 1989.52 | 3576.56 | 2704.43 | 3292.48 |
| CAMK2G | 421.28 | 456.22 | 485.98 | 723.51 | 663.17 | 723.72 |
| CASP3 | 1312.56 | 1250.37 | 1296.47 | 1910.57 | 2089.18 | 1949.74 |
| CAST | 209.29 | 220.87 | 244.92 | 449.47 | 329.67 | 381.32 |
| CCNH | 782.12 | 872.60 | 851.44 | 1376.58 | 1381.92 | 1453.81 |
| CD9 | 16.37 | 14.69 | 19.65 | 78.11 | 49.83 | 62.26 |
| CDK5 | 697.33 | 628.81 | 633.56 | 432.82 | 417.84 | 393.34 |
| CDKN1A | 1649.95 | 1599.17 | 1668.88 | 911.75 | 841.42 | 857.43 |
| CHRNB2 | 783.93 | 819.50 | 769.54 | 487.89 | 379.50 | 461.26 |
| CNTNAP1 | 616.14 | 726.57 | 665.24 | 452.03 | 381.42 | 440.04 |
| CNTNAP2 | 973.37 | 1071.75 | 1033.01 | 2066.80 | 1548.67 | 1973.08 |
| CRH | 94.72 | 71.21 | 70.31 | 558.32 | 530.92 | 492.39 |
| DDC | 123.59 | 103.80 | 90.40 | 39.70 | 55.58 | 34.67 |
| DGKB | 726.19 | 833.98 | 836.76 | 340.62 | 195.50 | 242.66 |
| DLX1 | 3834.84 | 3630.42 | 3785.89 | 997.54 | 1368.51 | 1146.78 |
| DLX2 | 2358.10 | 2358.32 | 2229.81 | 513.50 | 898.92 | 695.43 |
| DRD2 | 170.50 | 166.56 | 168.43 | 47.38 | 46.00 | 39.62 |
| DRD4 | 22.55 | 21.72 | 34.00 | 90.92 | 74.75 | 75.70 |
| EFNA1 | 16.37 | 14.69 | 19.65 | 55.06 | 53.67 | 50.94 |
| EGF | 16.37 | 18.10 | 19.65 | 60.19 | 70.92 | 62.26 |
| EGFL7 | 30.67 | 14.69 | 19.65 | 39.70 | 46.00 | 36.79 |
| EGFR | 115.47 | 126.73 | 125.17 | 197.20 | 201.25 | 175.45 |
| EGR1 | 94.72 | 91.73 | 102.76 | 40.98 | 49.83 | 48.11 |
| EMP2 | 85.70 | 49.48 | 64.13 | 142.14 | 157.17 | 132.29 |
| EPHA3 | 548.48 | 685.53 | 666.78 | 3187.27 | 2125.59 | 3003.14 |
| FAS | 209.29 | 245.01 | 224.84 | 176.71 | 92.00 | 149.27 |
| FGF12 | 589.07 | 523.80 | 601.11 | 396.97 | 300.92 | 349.48 |
| FGF14 | 3610.21 | 3549.56 | 3724.08 | 2538.04 | 1539.09 | 2142.17 |
| FN1 | 478.11 | 526.22 | 555.52 | 284.28 | 230.00 | 261.05 |
| FRMPD4 | 200.27 | 203.97 | 213.25 | 47.38 | 67.08 | 53.06 |
| GABRA1 | 259.81 | 296.90 | 294.37 | 140.86 | 143.75 | 152.10 |
| GABRA4 | 165.99 | 176.21 | 203.20 | 683.81 | 483.00 | 633.88 |
| GABRG2 | 392.41 | 377.77 | 390.18 | 608.26 | 576.92 | 594.26 |
| GAD1 | 3974.66 | 4476.47 | 4614.92 | 1243.41 | 1259.26 | 1267.05 |
| GAD2 | 739.72 | 903.98 | 851.44 | 413.62 | 552.00 | 489.56 |
| GAL3ST1 | 26.16 | 27.76 | 35.54 | 48.66 | 115.00 | 96.21 |
| GDNF | 27.06 | 35.00 | 25.50 | 19.01 | 19.82 | 15.32 |

| | | | | | | |
|---|---|---|---|---|---|---|
| **GNAI1** | 2721.64 | 2463.33 | 2572.86 | 1272.86 | 1082.92 | 1197.01 |
| **GPR37** | 99.23 | 85.69 | 108.94 | 299.65 | 228.08 | 256.81 |
| **GRIA2** | 2956.19 | 3463.86 | 3281.36 | 9068.81 | 8628.88 | 10127.18 |
| **GRIA4** | 2193.01 | 2352.29 | 2337.21 | 778.57 | 588.42 | 761.22 |
| **GRIN2A** | 127.20 | 118.28 | 100.44 | 60.19 | 67.08 | 59.43 |
| **GRM1** | 72.17 | 78.45 | 82.67 | 366.24 | 233.83 | 365.75 |
| **GUCY1B3** | 308.52 | 271.56 | 291.28 | 797.78 | 868.25 | 839.04 |
| **HAP1** | 317.54 | 343.97 | 347.68 | 103.72 | 84.33 | 120.97 |
| **HCN1** | 73.07 | 74.83 | 83.44 | 40.98 | 42.17 | 38.20 |
| **HNRNPM** | 7893.40 | 7399.63 | 7636.68 | 4780.27 | 3718.35 | 4101.81 |
| **IDH1** | 2553.85 | 2524.88 | 2450.79 | 1385.55 | 1217.09 | 1279.07 |
| **INHBB** | 25.26 | 24.14 | 29.36 | 55.06 | 63.25 | 53.77 |
| **IPCEF1** | 158.77 | 164.14 | 165.34 | 357.27 | 245.33 | 311.28 |
| **ITGA5** | 27.06 | 25.35 | 27.81 | 97.32 | 51.75 | 55.18 |
| **KCNA1** | 23.45 | 18.10 | 19.65 | 49.94 | 86.25 | 66.50 |
| **KIAA1161** | 324.76 | 301.73 | 284.33 | 505.81 | 521.34 | 505.83 |
| **LAMA2** | 18.94 | 24.14 | 19.65 | 60.19 | 44.08 | 48.11 |
| **LOX** | 51.42 | 25.35 | 50.99 | 72.99 | 103.50 | 72.87 |
| **MAPKAPK2** | 451.05 | 432.08 | 441.94 | 781.13 | 596.09 | 622.56 |
| **MGMT** | 16.37 | 14.69 | 21.63 | 87.08 | 70.92 | 67.92 |
| **MMP16** | 2183.99 | 2569.54 | 2323.30 | 1379.14 | 975.59 | 1165.17 |
| **MYD88** | 33.38 | 42.24 | 30.13 | 19.01 | 19.82 | 25.47 |
| **NAGLU** | 58.64 | 55.52 | 47.90 | 107.57 | 105.42 | 89.85 |
| **NEFH** | 78.48 | 77.24 | 84.99 | 19.01 | 19.82 | 28.30 |
| **NEFL** | 12890.14 | 11903.86 | 11811.21 | 4921.13 | 5293.86 | 5064.65 |
| **NEGR1** | 3594.88 | 3494.04 | 3574.96 | 1863.19 | 1512.26 | 1762.97 |
| **NFE2L2** | 480.82 | 477.94 | 465.90 | 962.97 | 701.50 | 858.14 |
| **NGFR** | 298.60 | 292.07 | 276.60 | 117.81 | 105.42 | 99.75 |
| **NKX6-2** | 79.39 | 56.73 | 51.77 | 19.01 | 19.82 | 15.32 |
| **NLGN4X** | 1475.84 | 1535.20 | 1479.59 | 2412.54 | 2238.68 | 2384.82 |
| **NOTCH1** | 849.78 | 957.09 | 895.48 | 1253.65 | 1657.93 | 1452.40 |
| **NOTCH3** | 566.52 | 491.22 | 465.90 | 996.26 | 1042.67 | 951.52 |
| **NOVA1** | 6173.09 | 6560.82 | 6729.62 | 4613.80 | 3413.60 | 4294.24 |
| **NR4A2** | 200.27 | 176.21 | 195.48 | 143.42 | 63.25 | 75.70 |
| **NRG1** | 1143.87 | 1188.82 | 1236.98 | 489.17 | 318.17 | 418.10 |
| **NSF** | 1699.56 | 1589.52 | 1582.35 | 1143.52 | 931.50 | 1008.12 |
| **NTF3** | 16.37 | 14.69 | 19.65 | 76.83 | 44.08 | 53.06 |
| **NTNG1** | 123.59 | 196.73 | 181.57 | 1083.34 | 734.09 | 993.97 |
| **NTRK1** | 821.82 | 811.05 | 720.86 | 44.82 | 19.82 | 28.30 |
| **NTS** | 786.63 | 784.50 | 939.52 | 332.94 | 429.34 | 358.68 |
| **OLIG2** | 414.97 | 434.49 | 368.54 | 48.66 | 126.50 | 76.40 |
| **PCSK2** | 866.02 | 884.67 | 812.04 | 2251.19 | 1602.34 | 2126.60 |
| **PDGFRB** | 42.40 | 44.66 | 34.00 | 186.96 | 130.33 | 145.03 |
| **PIK3R1** | 1713.09 | 1757.28 | 1755.42 | 810.58 | 678.50 | 793.76 |
| **PLA2G4F** | 44.20 | 47.07 | 40.18 | 30.73 | 19.82 | 28.30 |
| **PLCB1** | 277.85 | 329.49 | 319.10 | 541.67 | 421.67 | 487.43 |
| **PLS1** | 35.18 | 36.21 | 39.40 | 66.59 | 53.67 | 61.55 |
| **POLR2L** | 2763.14 | 2295.56 | 2542.73 | 1908.01 | 1176.84 | 1257.14 |
| **PRKACA** | 3229.53 | 3002.82 | 3024.08 | 1928.50 | 1600.43 | 1659.68 |
| **PRKACB** | 7919.56 | 8032.06 | 8019.14 | 5100.40 | 4042.27 | 4575.80 |

| | | | | | | |
|---|---|---|---|---|---|---|
| **PRKCB** | 978.78 | 867.78 | 944.93 | 514.78 | 488.75 | 503.71 |
| **PRKCG** | 82.09 | 83.28 | 76.49 | 125.49 | 145.67 | 132.29 |
| **PTPRR** | 95.62 | 109.83 | 112.80 | 190.80 | 172.50 | 171.20 |
| **RAB3C** | 3861.90 | 3689.56 | 3687.00 | 2462.48 | 2048.93 | 2126.60 |
| **RASGRP1** | 204.78 | 187.07 | 147.57 | 877.17 | 582.67 | 745.65 |
| **RET** | 253.49 | 287.25 | 302.87 | 185.68 | 101.58 | 140.78 |
| **RRAS** | 16.37 | 14.69 | 19.65 | 35.86 | 38.33 | 29.01 |
| **SCN1A** | 508.79 | 558.80 | 521.53 | 404.65 | 266.42 | 367.17 |
| **SERPINB6** | 420.38 | 504.49 | 450.44 | 325.26 | 210.83 | 289.35 |
| **SHH** | 650.42 | 708.46 | 649.01 | 72.99 | 76.67 | 68.62 |
| **SLA** | 16.37 | 14.69 | 19.65 | 53.78 | 59.42 | 54.47 |
| **SLC17A6** | 851.58 | 741.05 | 780.36 | 2742.92 | 2127.51 | 2479.62 |
| **SLC18A3** | 38.79 | 28.97 | 25.50 | 19.01 | 19.82 | 15.32 |
| **SLC1A2** | 931.87 | 918.47 | 881.57 | 1818.37 | 1539.09 | 1708.50 |
| **SLC32A1** | 371.67 | 308.97 | 356.18 | 128.05 | 172.50 | 133.00 |
| **SLC4A10** | 341.90 | 453.80 | 399.45 | 1430.37 | 898.92 | 1264.92 |
| **SORL1** | 221.02 | 232.94 | 256.51 | 376.48 | 360.34 | 379.90 |
| **STAMBPL1** | 340.99 | 341.56 | 385.54 | 1413.72 | 1063.76 | 1256.44 |
| **SYT1** | 6426.58 | 5947.71 | 5968.57 | 4420.43 | 2980.43 | 3757.28 |
| **SYT7** | 558.40 | 562.43 | 615.79 | 1274.14 | 1188.34 | 1197.72 |
| **TBR1** | 165.08 | 179.83 | 189.29 | 5013.33 | 2995.77 | 4062.19 |
| **TCIRG1** | 64.05 | 72.42 | 80.35 | 26.89 | 19.82 | 36.08 |
| **TENM2** | 870.53 | 957.09 | 912.48 | 548.07 | 308.58 | 489.56 |
| **TF** | 29.77 | 35.00 | 40.95 | 60.19 | 59.42 | 49.52 |
| **TH** | 304.01 | 339.14 | 352.32 | 124.21 | 226.17 | 184.65 |
| **THY1** | 1499.29 | 1357.79 | 1476.50 | 959.13 | 795.42 | 880.78 |
| **TNC** | 109.15 | 125.52 | 113.58 | 244.58 | 322.00 | 269.54 |
| **TNFRSF10D** | 45.11 | 54.31 | 50.99 | 19.01 | 19.82 | 25.47 |
| **TSPO** | 16.37 | 14.69 | 19.65 | 42.26 | 63.25 | 36.08 |
| **UGT8** | 61.34 | 55.52 | 61.81 | 151.10 | 195.50 | 194.55 |
| **XK** | 135.32 | 115.86 | 137.53 | 58.90 | 19.82 | 36.08 |

Supplementary Table 5

| NAME | IPS d100 _1 | IPS d100 _2 | IPS d100 _3 | IVS d100 _1 | IVS d100 _2 | IVS d100 _3 |
|---|---|---|---|---|---|---|
| NTRK1 | 247.37 | 211.78 | 146.99 | 1.72 | 1 | 1 |
| SLC32A1 | 718.68 | 684.43 | 808.95 | 20.41 | 1 | 3.14 |
| NKX6-2 | 122.49 | 97.9 | 56.35 | 1.72 | 1 | 1 |
| SLC18A3 | 61.67 | 52.55 | 17.9 | 1 | 1 | 1 |
| ADCYAP1 | 191.99 | 177.51 | 177.21 | 5.46 | 1 | 6.38 |
| GAD2 | 1814.41 | 1844.38 | 2478.97 | 91.45 | 24.7 | 32.32 |
| GAD1 | 11286.15 | 11290.31 | 12911.05 | 418.6 | 127.08 | 323.23 |
| MMP2 | 421.13 | 400.23 | 100.3 | 8.26 | 1 | 15.3 |
| DDC | 73.62 | 87.82 | 45.36 | 1 | 4.59 | 1 |
| TLR4 | 70.36 | 73.71 | 56.35 | 4.52 | 1 | 1 |
| TH | 618.77 | 682.41 | 627.67 | 26.02 | 10.08 | 43.66 |
| AQP4 | 157.24 | 174.49 | 94.8 | 10.13 | 5.51 | 4.76 |
| DLX1 | 3953.75 | 3826.69 | 4772.48 | 403.65 | 133.48 | 80.94 |
| NGFR | 413.52 | 431.47 | 325.53 | 12.94 | 17.39 | 28.26 |
| DLX2 | 1879.57 | 1838.34 | 2083.44 | 195.21 | 84.12 | 33.13 |
| FRMPD4 | 294.07 | 342.79 | 284.33 | 19.48 | 7.34 | 27.45 |
| TENM2 | 1360.48 | 1366.69 | 1333.58 | 90.52 | 53.04 | 99.57 |
| DRD2 | 284.29 | 279.3 | 218.41 | 18.54 | 15.56 | 12.87 |
| OLIG2 | 263.66 | 230.93 | 177.21 | 24.15 | 21.05 | 1 |
| SHH | 335.33 | 294.42 | 443.64 | 46.59 | 26.53 | 2.33 |
| NEFL | 23874.6 | 25396.2 | 24735.72 | 1854.33 | 1277.93 | 2573.54 |
| FGF14 | 5010.39 | 4967.49 | 6483.7 | 438.23 | 253.23 | 581.73 |
| GABRA1 | 648.09 | 649.16 | 602.95 | 57.8 | 33.84 | 59.06 |
| MBP | 135.52 | 115.03 | 67.34 | 7.33 | 1 | 18.54 |
| RET | 282.12 | 264.18 | 309.05 | 30.7 | 4.59 | 42.04 |
| PTPRR | 296.24 | 298.45 | 325.53 | 33.5 | 19.22 | 33.13 |
| HAP1 | 543.84 | 615.9 | 646.9 | 60.61 | 53.04 | 81.75 |
| RAB3C | 5650.02 | 5742.48 | 5725.6 | 693.41 | 538.43 | 623.86 |
| CALB1 | 2191.24 | 2185.01 | 2316.91 | 287.74 | 90.52 | 369.42 |
| GRIA4 | 3629.05 | 3755.13 | 3838.6 | 407.39 | 431.48 | 469.9 |
| ACHE | 2914.49 | 2984.18 | 3179.38 | 336.35 | 291.62 | 469.09 |
| SLC12A5 | 1248.63 | 1224.6 | 1251.18 | 149.4 | 137.14 | 188.71 |
| PDE1B | 219.14 | 208.76 | 177.21 | 28.83 | 4.59 | 52.57 |
| DRD1 | 84.48 | 81.78 | 92.06 | 9.2 | 8.25 | 19.35 |
| NEFH | 178.96 | 189.61 | 174.46 | 26.96 | 23.79 | 26.64 |
| XK | 226.74 | 201.7 | 185.45 | 28.83 | 22.88 | 37.18 |
| DBH | 9.55 | 8.21 | 2.75 | 1 | 1 | 1 |
| HCN1 | 287.55 | 293.41 | 270.59 | 41.91 | 28.36 | 58.25 |
| CYP4X1 | 605.74 | 663.26 | 608.44 | 82.11 | 58.53 | 144.14 |
| RYR3 | 464.56 | 492.95 | 449.13 | 80.24 | 50.3 | 102.81 |
| GABRB2 | 1803.55 | 1902.83 | 1674.17 | 284 | 277.91 | 375.09 |
| SORCS3 | 414.61 | 444.58 | 506.81 | 66.22 | 56.7 | 116.59 |
| S100B | 1103.11 | 940.4 | 564.49 | 218.57 | 111.54 | 265.69 |
| NPY | 887 | 779.16 | 449.13 | 136.32 | 164.56 | 183.04 |
| CD8A | 107.28 | 78.75 | 53.6 | 10.13 | 10.99 | 34.75 |
| DGKB | 1928.44 | 1820.2 | 2034 | 532.64 | 451.59 | 401.83 |
| GRM5 | 1407.18 | 1410.03 | 1352.81 | 354.11 | 317.22 | 332.14 |
| SCN1A | 1402.83 | 1423.13 | 1487.4 | 321.39 | 285.22 | 441.54 |
| RIMS1 | 905.46 | 896.06 | 883.12 | 207.36 | 186.5 | 286.76 |

| | | | | | | |
|---|---|---|---|---|---|---|
| **NOVA1** | 5993.19 | 6335.05 | 7288.49 | 1771.14 | 1600.61 | 1651.37 |
| **HMOX1** | 99.68 | 69.68 | 111.28 | 26.96 | 6.42 | 42.04 |
| **HLA-DRA** | 5.2 | 3.17 | 2.75 | 1 | 1 | 1 |
| **PRKCB** | 1161.75 | 1175.22 | 1072.64 | 212.03 | 397.66 | 321.61 |
| **CCND1** | 816.41 | 847.69 | 723.8 | 270.92 | 116.11 | 273.8 |
| **PTPRN2** | 3404.26 | 3319.77 | 3385.39 | 924.29 | 863.85 | 1016.07 |
| **FOS** | 1079.22 | 1019.01 | 1163.28 | 251.29 | 248.66 | 435.05 |
| **RYR1** | 150.72 | 136.2 | 188.19 | 47.52 | 32.93 | 56.63 |
| **ADCY8** | 345.11 | 321.63 | 314.54 | 83.04 | 53.04 | 149 |
| **EFNA5** | 671.98 | 681.4 | 800.71 | 180.25 | 168.22 | 281.09 |
| **GRIN2D** | 644.83 | 627.99 | 715.56 | 198.01 | 190.16 | 231.66 |
| **THY1** | 3755.02 | 3907.31 | 3794.65 | 1165.44 | 1132.59 | 1302.12 |
| **CACNB2** | 604.65 | 695.51 | 674.36 | 199.88 | 155.42 | 276.23 |
| **PRKACB** | 10239.29 | 10335.94 | 11411.33 | 3245.19 | 3635.39 | 3452.76 |
| **CX3CL1** | 176.78 | 157.36 | 136 | 59.67 | 23.79 | 70.4 |
| **PLXNC1** | 1633.06 | 1627.71 | 1572.54 | 578.44 | 564.94 | 501.5 |
| **NRG1** | 1217.13 | 1172.19 | 1240.19 | 423.28 | 418.68 | 398.59 |
| **SLC1A1** | 1118.31 | 1163.12 | 1193.5 | 372.8 | 380.29 | 452.07 |
| **OLFM3** | 1104.19 | 1089.56 | 907.84 | 389.63 | 336.41 | 362.93 |
| **FGF12** | 1629.8 | 1574.3 | 1720.87 | 510.21 | 527.46 | 721.1 |
| **NSF** | 2549.61 | 2666.73 | 2531.15 | 895.31 | 902.24 | 995.81 |
| **CADPS** | 2328.07 | 2355.33 | 2341.63 | 858.85 | 761.47 | 961.77 |
| **COL4A1** | 919.58 | 846.68 | 608.44 | 304.57 | 302.59 | 273.8 |
| **MAGEE1** | 1269.26 | 1301.19 | 1306.11 | 474.69 | 477.18 | 507.98 |
| **NEGR1** | 5225.41 | 5074.32 | 5198.23 | 1587.93 | 1950.71 | 2387.97 |
| **FN1** | 867.46 | 747.92 | 677.11 | 326.07 | 226.72 | 328.9 |
| **XBP1** | 574.25 | 541.32 | 498.57 | 208.29 | 208.44 | 228.42 |
| **GSN** | 513.43 | 445.58 | 322.78 | 208.29 | 109.71 | 195.19 |
| **TNR** | 1203.02 | 1240.72 | 1198.99 | 451.32 | 572.25 | 490.16 |
| **SYT13** | 3545.43 | 3484.04 | 3434.83 | 1252.37 | 1485.43 | 1633.55 |
| **PIK3R1** | 2031.6 | 2072.14 | 2245.49 | 766.32 | 822.71 | 1092.24 |
| **MAPK10** | 4401.17 | 4492.83 | 4511.55 | 1753.38 | 1763.32 | 2178.9 |
| **ATP8A2** | 3138.2 | 3179.69 | 3127.19 | 1005.61 | 1395.85 | 1640.84 |
| **UNC13A** | 2157.58 | 2251.53 | 2388.32 | 928.96 | 914.12 | 1127.89 |
| **PLA2G16** | 810.99 | 850.71 | 792.47 | 419.54 | 183.76 | 474.76 |
| **INA** | 13472.19 | 13702.93 | 13718.59 | 5517.48 | 5740.56 | 6723.29 |
| **SCN2A** | 3544.35 | 3579.78 | 3514.48 | 1468.29 | 1652.71 | 1800.48 |
| **SNCB** | 1032.52 | 1038.16 | 998.48 | 406.45 | 514.66 | 499.07 |
| **MAPT** | 3923.35 | 3780.33 | 3682.03 | 1650.56 | 1744.12 | 1875.84 |
| **CACNB4** | 786.01 | 806.37 | 773.25 | 343.83 | 348.29 | 413.17 |
| **SERPINB6** | 693.7 | 667.3 | 613.94 | 305.5 | 264.2 | 366.17 |
| **CNTN1** | 3898.37 | 4136.07 | 3764.44 | 1733.75 | 1968.08 | 1952.01 |
| **ITPR2** | 237.6 | 236.97 | 251.37 | 147.54 | 96 | 107.68 |
| **PLXNB3** | 90.99 | 73.71 | 70.08 | 37.24 | 23.79 | 52.57 |
| **CHRNA7** | 853.34 | 890.01 | 894.1 | 444.78 | 372.98 | 459.36 |
| **PINK1** | 1554.87 | 1521.89 | 1325.34 | 687.8 | 663.66 | 794.84 |
| **GAL3ST1** | 51.9 | 43.48 | 34.38 | 12 | 24.7 | 26.64 |
| **GNG2** | 10579.2 | 10273.45 | 10367.58 | 4803.36 | 5226.84 | 5385.42 |
| **FAS** | 200.68 | 181.55 | 185.45 | 124.17 | 73.15 | 84.18 |
| **NRXN1** | 1652.6 | 1576.31 | 2212.53 | 921.48 | 929.66 | 852.38 |

| | | | | | | |
|---|---|---|---|---|---|---|
| **STAT3** | 1927.35 | 1833.3 | 1935.11 | 987.85 | 798.95 | 1069.55 |
| **PLCB4** | 1102.02 | 1052.27 | 1047.92 | 562.55 | 482.67 | 580.1 |
| **CNTNAP1** | 995.6 | 1085.52 | 1366.54 | 571.9 | 514.66 | 665.19 |
| **NOL3** | 205.02 | 227.9 | 240.38 | 127.91 | 88.69 | 126.31 |
| **ATP6V0C** | 9294.5 | 8822.25 | 9439.18 | 4460.32 | 4580.57 | 5130.16 |
| **PPARGC1A** | 644.83 | 714.66 | 718.31 | 382.15 | 207.52 | 503.93 |
| **GALC** | 471.08 | 496.98 | 451.88 | 242.88 | 266.94 | 238.95 |
| **AMIGO1** | 233.25 | 218.83 | 193.69 | 117.62 | 107.89 | 115.78 |
| **ATP13A2** | 1911.06 | 1849.42 | 1803.27 | 908.39 | 967.14 | 1071.17 |
| **CACNA1C** | 1363.74 | 1463.44 | 1448.94 | 694.34 | 643.55 | 929.36 |
| **KCNB1** | 429.81 | 415.35 | 468.36 | 223.25 | 224.89 | 259.21 |
| **PLA2G4C** | 336.42 | 348.84 | 278.83 | 154.08 | 176.44 | 188.71 |
| **SOD2** | 2578.93 | 2633.47 | 2915.7 | 1450.53 | 1409.56 | 1518.48 |
| **CAST** | 405.92 | 432.48 | 460.12 | 244.75 | 173.7 | 281.9 |
| **SLC17A6** | 848.99 | 946.45 | 833.67 | 433.56 | 278.82 | 714.62 |
| **PRKACA** | 3479.19 | 3684.59 | 3693.02 | 1897.33 | 1980.87 | 2048.44 |
| **ATP6V0E2** | 1851.33 | 1940.12 | 1882.92 | 1012.15 | 1036.61 | 1064.69 |
| **UCHL1** | 15510.54 | 15754.77 | 16756.47 | 8345.93 | 9026.75 | 9034.37 |
| **GNPTG** | 833.79 | 877.92 | 803.46 | 473.75 | 437.88 | 473.95 |
| **UGCG** | 1881.74 | 1858.49 | 1841.72 | 1028.04 | 1007.36 | 1071.17 |
| **AMPH** | 2412.78 | 2537.74 | 2465.23 | 1410.34 | 1213.95 | 1503.08 |
| **PCSK2** | 781.66 | 852.73 | 685.35 | 435.43 | 289.79 | 576.05 |
| **SLC9A6** | 1390.89 | 1478.56 | 1383.02 | 772.86 | 794.37 | 833.74 |
| **NQO1** | 491.71 | 479.85 | 484.84 | 310.18 | 159.99 | 353.21 |
| **COL4A2** | 663.29 | 700.55 | 531.53 | 368.13 | 392.17 | 319.99 |
| **CAMK2B** | 1057.5 | 1094.59 | 1130.32 | 544.79 | 561.28 | 764.86 |
| **MMP24** | 866.37 | 906.14 | 762.26 | 407.39 | 434.22 | 610.09 |
| **ERLEC1** | 2868.88 | 2933.79 | 2912.95 | 1634.67 | 1730.41 | 1635.98 |
| **EFNB3** | 2899.29 | 2891.47 | 2901.96 | 1899.19 | 1592.38 | 1541.17 |
| **ATP6V1G2** | 2316.13 | 2349.28 | 2281.2 | 1287.89 | 1420.53 | 1327.24 |
| **MYC** | 411.35 | 390.16 | 416.17 | 216.7 | 221.23 | 273.8 |
| **ATP6V1A** | 6560.06 | 6655.52 | 6719.92 | 3848.08 | 3860.26 | 4024.05 |
| **ARC** | 151.81 | 160.38 | 144.24 | 87.71 | 101.49 | 82.56 |
| **INPP5F** | 4583.61 | 4787.1 | 4665.36 | 2659.12 | 2763.34 | 2985.19 |
| **SYT1** | 10729.06 | 10449.82 | 10510.4 | 5631.52 | 6854.85 | 6584.72 |
| **GNAI1** | 3376.02 | 3522.34 | 3308.48 | 1815.07 | 2149.07 | 2184.58 |
| **L1CAM** | 4473.93 | 4787.1 | 5126.81 | 2657.25 | 2871.21 | 3182.91 |
| **PRNP** | 5133.11 | 5381.69 | 5069.13 | 3176.95 | 2388.56 | 3886.29 |
| **ATP6V0D1** | 4180.72 | 4166.31 | 4206.66 | 2506.76 | 2444.32 | 2691.04 |
| **POLR2K** | 2349.79 | 2354.32 | 2336.14 | 1370.14 | 1595.13 | 1337.77 |
| **CHMP2B** | 1085.73 | 1159.09 | 1220.96 | 677.52 | 736.79 | 719.48 |
| **SNAP91** | 2989.42 | 3016.43 | 3393.63 | 1789.83 | 2022.01 | 1998.2 |
| **ABAT** | 4226.33 | 4079.64 | 4365.97 | 2650.71 | 2679.25 | 2524.92 |
| **CYCS** | 959.76 | 1059.32 | 1067.15 | 602.74 | 614.3 | 703.28 |
| **GABRG2** | 877.23 | 874.9 | 715.56 | 520.49 | 479.92 | 540.4 |
| **MMP16** | 2008.8 | 2205.17 | 2303.17 | 1242.09 | 1338.26 | 1503.89 |
| **MAP2K1** | 1310.53 | 1383.83 | 1226.46 | 841.1 | 822.71 | 807 |
| **GRIN1** | 297.33 | 292.4 | 270.59 | 181.19 | 165.47 | 196 |
| **GABRA4** | 530.81 | 612.88 | 597.46 | 322.33 | 296.19 | 485.29 |
| **GABRB3** | 3042.63 | 3072.87 | 2869 | 1794.51 | 1845.59 | 2060.59 |

| | | | | | | |
|---|---|---|---|---|---|---|
| **TNFRSF1A** | 309.27 | 327.67 | 289.82 | 206.42 | 136.22 | 248.68 |
| **GNAO1** | 5317.72 | 5530.84 | 4896.09 | 3151.71 | 3273.41 | 3647.24 |
| **CDK5** | 1034.69 | 979.71 | 979.25 | 629.85 | 619.78 | 666 |
| **OPTN** | 2114.14 | 2184 | 2196.05 | 1334.63 | 1333.69 | 1529.01 |
| **NTS** | 2904.72 | 3426.6 | 3357.92 | 2155.31 | 1553.08 | 2567.06 |
| **ATP6V1H** | 2631.05 | 2662.7 | 2704.2 | 1605.69 | 1772.46 | 1805.34 |
| **STAMBPL1** | 450.45 | 497.99 | 564.49 | 298.02 | 284.31 | 399.4 |
| **APP** | 15729.9 | 15920.05 | 15481.99 | 9745.21 | 9856.75 | 11019.71 |
| **CDKN1A** | 1488.62 | 1546.08 | 1407.74 | 1092.53 | 749.58 | 1049.29 |
| **BACE1** | 1964.27 | 1947.18 | 1808.76 | 1146.75 | 1249.6 | 1357.22 |
| **NLGN4X** | 2361.74 | 2275.71 | 2503.69 | 1681.41 | 1519.26 | 1486.06 |
| **DDIT3** | 1005.37 | 1159.09 | 1144.05 | 746.69 | 549.4 | 888.03 |
| **BCAS2** | 1961.02 | 1955.24 | 2075.2 | 1313.13 | 1382.14 | 1287.53 |
| **PRKCE** | 410.27 | 398.22 | 314.54 | 189.6 | 255.06 | 302.16 |
| **IDE** | 342.94 | 328.68 | 292.57 | 491.51 | 457.99 | 504.74 |
| **GUSB** | 595.97 | 659.23 | 633.16 | 984.11 | 960.74 | 936.65 |
| **CCNH** | 870.71 | 880.94 | 863.89 | 1299.11 | 1364.77 | 1347.5 |
| **EPHA4** | 1368.08 | 1363.67 | 1435.21 | 2307.67 | 2272.47 | 1845.04 |
| **RAF1** | 1652.6 | 1728.49 | 1654.95 | 2530.13 | 2717.64 | 2569.49 |
| **SGPL1** | 508 | 429.46 | 451.88 | 690.61 | 771.52 | 697.6 |
| **ARHGEF10** | 201.76 | 176.51 | 116.78 | 285.87 | 248.66 | 237.33 |
| **STX2** | 997.77 | 1011.96 | 1097.36 | 1652.43 | 1577.76 | 1618.15 |
| **DLG3** | 565.56 | 604.81 | 553.51 | 855.12 | 904.07 | 938.27 |
| **CNTN4** | 270.18 | 265.19 | 254.11 | 363.46 | 383.94 | 493.4 |
| **GNAI2** | 2865.62 | 2925.73 | 2926.68 | 4696.8 | 4946.21 | 4173.15 |
| **LSM2** | 733.88 | 726.75 | 597.46 | 1101.88 | 1225.83 | 949.62 |
| **MUTYH** | 134.43 | 105.96 | 97.55 | 171.84 | 191.98 | 175.75 |
| **NES** | 4090.58 | 3811.57 | 3629.85 | 6583.06 | 6403.28 | 5718.47 |
| **TCERG1** | 1224.74 | 1291.11 | 1358.3 | 2076.79 | 2127.13 | 2084.09 |
| **GUCY1B3** | 738.23 | 699.54 | 515.05 | 1019.63 | 1114.31 | 1050.1 |
| **SLC1A2** | 1279.03 | 1343.52 | 1144.05 | 2184.28 | 2115.25 | 1862.06 |
| **CDK5RAP3** | 1553.78 | 1709.34 | 1696.15 | 2704.92 | 2522.94 | 2975.47 |
| **C5** | 105.11 | 100.92 | 75.58 | 155.01 | 150.85 | 161.97 |
| **FAM126A** | 733.88 | 682.41 | 635.91 | 1024.3 | 1115.22 | 1277.81 |
| **TAF6L** | 143.12 | 186.58 | 122.27 | 251.29 | 245 | 261.64 |
| **ACTN1** | 410.27 | 493.96 | 468.36 | 810.25 | 680.11 | 812.67 |
| **SNRPA** | 788.18 | 825.52 | 850.15 | 1406.6 | 1454.35 | 1302.12 |
| **SNCAIP** | 380.95 | 312.56 | 309.05 | 607.42 | 637.15 | 472.33 |
| **TAF4** | 611.17 | 566.52 | 506.81 | 1000 | 996.39 | 961.77 |
| **TP53** | 439.59 | 456.67 | 353 | 785.01 | 788.89 | 634.4 |
| **IKBKB** | 272.35 | 244.03 | 240.38 | 462.53 | 455.24 | 424.52 |
| **DOT1L** | 1122.66 | 1200.41 | 1286.88 | 2134.74 | 2118.9 | 2164.32 |
| **HDAC1** | 654.61 | 602.8 | 622.18 | 1165.44 | 1190.18 | 1069.55 |
| **NFE2L2** | 701.3 | 646.13 | 605.7 | 1250.5 | 1205.72 | 1175.7 |
| **MAN2B1** | 174.61 | 208.76 | 160.73 | 340.09 | 324.53 | 362.12 |
| **DLGAP1** | 1077.05 | 1209.48 | 946.29 | 1849.65 | 2203.91 | 2074.37 |
| **KATNA1** | 442.84 | 384.11 | 347.5 | 800.9 | 780.66 | 713.81 |
| **PQBP1** | 1211.7 | 1272.97 | 1330.83 | 2385.25 | 2624.4 | 2554.09 |
| **EPHA5** | 805.56 | 748.93 | 641.4 | 1385.1 | 1783.43 | 1268.89 |
| **GPR37** | 175.7 | 175.5 | 86.56 | 245.68 | 315.39 | 325.66 |

| | | | | | | |
|---|---|---|---|---|---|---|
| **PCNA** | 272.35 | 272.25 | 256.86 | 617.7 | 624.35 | 389.67 |
| **MAPKAPK2** | 384.2 | 450.62 | 388.7 | 824.27 | 826.37 | 854 |
| **MTA2** | 1319.21 | 1343.52 | 1410.49 | 2826.43 | 2978.16 | 2588.12 |
| **IPCEF1** | 155.07 | 157.36 | 108.54 | 302.7 | 281.57 | 284.33 |
| **KIAA1161** | 464.56 | 448.61 | 399.69 | 844.83 | 1033.87 | 832.93 |
| **LOX** | 52.99 | 48.52 | 15.15 | 92.39 | 79.55 | 72.02 |
| **AGER** | 99.68 | 118.06 | 67.34 | 219.51 | 200.21 | 205.73 |
| **PLCB3** | 102.94 | 92.86 | 116.78 | 221.38 | 237.69 | 226.8 |
| **SLC4A10** | 612.25 | 641.09 | 644.15 | 1282.28 | 1267.88 | 1702.42 |
| **SP1** | 144.21 | 133.17 | 103.04 | 321.39 | 309.9 | 263.26 |
| **SOX9** | 1606.99 | 1693.22 | 1578.04 | 4323.85 | 4018.4 | 3467.34 |
| **BID** | 185.47 | 177.51 | 146.99 | 395.24 | 405.88 | 440.73 |
| **PARP1** | 1450.62 | 1380.8 | 1333.58 | 3541.49 | 3559.52 | 3246.93 |
| **CNTF** | 100.77 | 97.9 | 64.59 | 206.42 | 239.52 | 229.23 |
| **PHF19** | 68.19 | 64.64 | 23.39 | 139.12 | 156.33 | 111.73 |
| **INHBB** | 60.59 | 68.67 | 34.38 | 171.84 | 147.19 | 116.59 |
| **NOTCH3** | 386.37 | 353.88 | 289.82 | 1013.08 | 946.12 | 827.26 |
| **NAGLU** | 63.84 | 60.61 | 39.87 | 148.47 | 132.57 | 165.21 |
| **PRKCG** | 51.9 | 53.56 | 53.6 | 142.86 | 132.57 | 164.4 |
| **CASP7** | 71.45 | 53.56 | 31.63 | 190.53 | 150.85 | 128.75 |
| **NELL2** | 3109.96 | 3053.72 | 3239.81 | 8658.13 | 10321.11 | 9820.4 |
| **NOTCH1** | 742.57 | 682.41 | 605.7 | 2474.05 | 2453.46 | 1771.3 |
| **GRIA3** | 1721.02 | 1684.15 | 1646.71 | 4983.76 | 6059.58 | 6160.1 |
| **AKT3** | 3013.31 | 3023.49 | 2706.94 | 9821.85 | 10960.98 | 9221.56 |
| **CASP6** | 243.03 | 213.79 | 199.18 | 771.93 | 831.85 | 741.36 |
| **EMP2** | 63.84 | 53.56 | 2.75 | 151.27 | 162.73 | 119.83 |
| **GRIN3B** | 1.09 | 1.01 | 2.75 | 7.33 | 7.34 | 5.57 |
| **PLS1** | 38.87 | 24.33 | 2.75 | 110.15 | 55.78 | 123.88 |
| **EGF** | 28.01 | 21.31 | 2.75 | 76.5 | 106.97 | 80.94 |
| **CDK2** | 71.45 | 54.56 | 12.4 | 291.48 | 268.77 | 192.76 |
| **EPHA7** | 550.35 | 640.09 | 498.57 | 3254.53 | 3759.71 | 3076.76 |
| **HGF** | 12.8 | 18.28 | 2.75 | 78.37 | 85.03 | 65.54 |
| **ADRA2A** | 97.51 | 80.77 | 26.14 | 343.83 | 617.95 | 843.46 |
| **DRD4** | 5.2 | 1.01 | 2.75 | 33.5 | 22.88 | 24.21 |
| **GRIN2C** | 12.8 | 1.01 | 2.75 | 45.65 | 45.73 | 60.68 |
| **CA2** | 157.24 | 166.43 | 141.5 | 1775.81 | 561.28 | 2174.85 |
| **KCNJ10** | 11.72 | 5.18 | 2.75 | 71.82 | 58.53 | 63.92 |
| **NOSTRIN** | 1.09 | 1.01 | 2.75 | 19.48 | 18.31 | 14.49 |
| **MGMT** | 1.94 | 1.15 | 2.75 | 33.5 | 23.79 | 16.11 |
| **NTF3** | 1.09 | 1.01 | 2.75 | 41.91 | 28.36 | 41.23 |
| **TSPO** | 3.03 | 3.17 | 2.75 | 59.67 | 96 | 65.54 |
| **EFNA1** | 1.09 | 1.01 | 2.75 | 65.28 | 97.83 | 123.07 |
| **TBR1** | 59.5 | 67.67 | 39.87 | 5189.4 | 6225.95 | 6626.04 |
| **SLA** | 1.09 | 1.01 | 2.75 | 860.72 | 1258.74 | 1202.44 |

Supplementary Figure 3

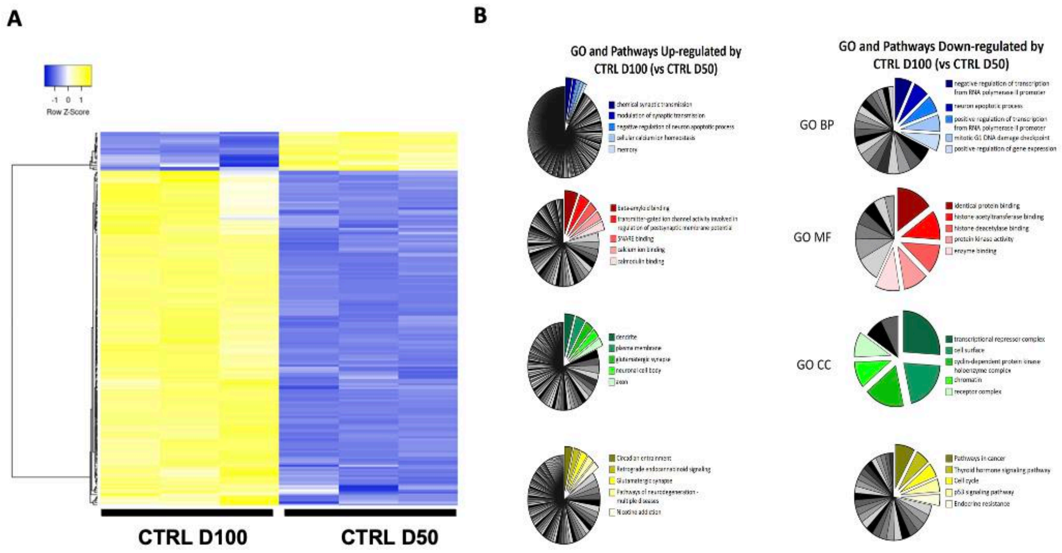


**Figure S3. A)** Heat map of supervised hierarchical clustering of 183 differentially expressed (DE) genes between D50 CTRL and D100 CTRL (164 genes up- and 19 genes down-regulated in D100 CTRL). Average linkege and Pearson Distance Measurement Methods was performed for Hierarchical cluster using Heatmapper tools (http://www.heatmapper.ca) . The significant DE genes are selected by multiple t-test performed with Graph Pad Prism 6, with *pvalue* $\leq 0.05$ and a fold change cut-off values of $\geq 1.5$ or $\leq 0.66$ for upmodulated and downregulated genes respectively. **B)** Gene Ontology (GO) and KEGG Pathways enrichment analysis of 183 down- and up-regulated genes by D100 CTRL (respect to D50 CTRL) have been analyzed by DAVID database. An overview of top 5 significantly enriched terms in KEGG pathways and three GO categories (biological process (BP), cellular component (CC), and molecular function (MF)) are showed. The cut-off of p-value was set to 0.05 and are selected only enriched terms in which are represented a minimum of 3 genes. The −log10 p-value associated with each category is represented by the size of the pie slice.

Supplementary Figure 4

Supplementary Table 6

| Probe Name | IPS d50 _1 | IPS d50 _2 | IPS d50 _3 | IPS d100 _1 | IPS d100 _2 | IPS d100 _3 |
|---|---|---|---|---|---|---|
| ACHE | 1207.81 | 1199.61 | 1183.83 | 2656.07 | 2719.58 | 2897.47 |
| ADCY5 | 559.67 | 613.33 | 615.74 | 915.24 | 897.43 | 919.96 |
| ADCY8 | 122.39 | 122.26 | 159.50 | 314.51 | 293.11 | 286.65 |
| ADCYAP1 | 117.21 | 73.50 | 109.87 | 174.96 | 161.77 | 161.49 |
| ADORA1 | 138.81 | 106.00 | 110.61 | 243.25 | 223.31 | 299.17 |
| AGER | 214.86 | 231.38 | 221.71 | 90.84 | 107.59 | 61.37 |
| AMIGO1 | 131.04 | 124.58 | 110.61 | 212.57 | 199.43 | 176.51 |
| APOE | 23.88 | 20.09 | 21.00 | 278.88 | 273.82 | 274.14 |
| AQP4 | 10.91 | 1.16 | 1.00 | 143.30 | 159.02 | 86.40 |
| ARC | 102.52 | 60.73 | 89.14 | 138.35 | 146.16 | 131.45 |
| ARHGAP44 | 284.00 | 302.20 | 312.07 | 521.35 | 553.02 | 549.49 |
| ATP13A2 | 1014.23 | 1067.26 | 1030.51 | 1741.61 | 1685.44 | 1643.38 |
| ATP2B3 | 128.44 | 167.53 | 128.39 | 242.26 | 264.64 | 234.09 |
| ATP6V0C | 5725.75 | 5402.20 | 5353.72 | 8470.39 | 8040.01 | 8602.24 |
| ATP6V0E2 | 1206.08 | 1053.33 | 1057.17 | 1687.18 | 1768.10 | 1715.97 |
| ATP6V1G2 | 1379.78 | 1364.46 | 1354.18 | 2110.76 | 2140.98 | 2078.93 |
| B4GALT6 | 570.04 | 557.61 | 547.60 | 891.49 | 902.94 | 819.83 |
| BACE1 | 1060.03 | 1125.31 | 1030.51 | 1790.11 | 1774.53 | 1648.39 |
| BCHE | 246.84 | 205.84 | 206.16 | 709.39 | 644.86 | 351.74 |
| BDNF | 51.53 | 47.96 | 57.29 | 103.71 | 107.59 | 101.42 |
| BNIP3 | 685.84 | 629.59 | 632.78 | 962.74 | 1011.32 | 1035.10 |
| CA2 | 103.38 | 68.85 | 66.92 | 143.30 | 151.67 | 128.95 |
| CACNA1B | 432.64 | 527.42 | 526.86 | 724.23 | 784.47 | 804.81 |
| CACNB2 | 344.49 | 364.89 | 377.25 | 551.04 | 633.84 | 614.57 |
| CADM3 | 1868.04 | 1872.95 | 1812.64 | 2738.21 | 2832.55 | 2957.55 |
| CALB1 | 1330.52 | 1155.49 | 1100.13 | 1996.95 | 1991.27 | 2111.47 |
| CALB2 | 232.14 | 225.58 | 215.05 | 1048.84 | 1097.65 | 667.14 |
| CALM1 | 3181.60 | 2794.74 | 2954.73 | 5478.61 | 5652.11 | 5152.85 |
| CAMK2D | 1752.24 | 1939.13 | 1892.63 | 3499.27 | 3516.77 | 3916.27 |
| CAMK2G | 389.43 | 421.78 | 451.32 | 666.83 | 734.87 | 734.72 |
| CAST | 186.34 | 195.40 | 220.23 | 369.93 | 394.14 | 419.32 |
| CCL5 | 35.98 | 23.58 | 29.14 | 13.65 | 15.75 | 2.50 |
| CCND1 | 1241.51 | 1276.23 | 1254.19 | 744.03 | 772.53 | 659.63 |
| CD44 | 10.91 | 13.13 | 9.89 | 71.05 | 67.18 | 41.34 |
| CDK2 | 126.71 | 150.12 | 112.84 | 65.11 | 49.73 | 11.30 |
| CDS1 | 79.18 | 103.68 | 108.39 | 229.40 | 265.56 | 146.47 |
| CHL1 | 1469.65 | 1528.15 | 1613.41 | 3029.18 | 3072.26 | 2454.41 |
| CHMP2B | 661.64 | 676.02 | 720.17 | 989.46 | 1056.32 | 1112.70 |
| CHRNA7 | 404.98 | 370.70 | 389.10 | 777.67 | 811.10 | 814.82 |
| CLU | 2347.66 | 2072.63 | 2170.38 | 7786.52 | 7881.12 | 7783.69 |
| CNR1 | 627.94 | 640.03 | 609.82 | 1373.46 | 1217.96 | 1548.26 |
| CNTN1 | 1816.19 | 1904.30 | 1876.34 | 3552.71 | 3769.34 | 3430.65 |
| CNTNAP1 | 576.09 | 681.83 | 623.15 | 907.32 | 989.27 | 1245.37 |
| COL4A1 | 274.49 | 272.02 | 250.60 | 838.04 | 771.61 | 554.49 |
| CPLX1 | 284.86 | 283.63 | 296.52 | 560.94 | 542.92 | 496.92 |
| CRH | 76.59 | 51.44 | 52.84 | 169.03 | 238.92 | 429.33 |
| CSF1 | 50.67 | 43.31 | 45.44 | 105.69 | 75.44 | 73.88 |
| CX3CL1 | 64.49 | 68.85 | 69.88 | 161.11 | 143.41 | 123.95 |
| CXCR4 | 513.00 | 407.85 | 466.87 | 816.27 | 788.14 | 742.23 |

| | | | | | | |
|---|---|---|---|---|---|---|
| **CYP4X1** | 273.63 | 301.04 | 314.30 | 552.03 | 604.45 | 554.49 |
| **DAGLA** | 336.71 | 366.05 | 300.22 | 546.09 | 552.10 | 539.47 |
| **DGKB** | 681.52 | 785.15 | 787.57 | 1757.45 | 1658.80 | 1853.65 |
| **DLGAP1** | 347.08 | 391.59 | 403.17 | 981.55 | 1102.24 | 862.39 |
| **DRD2** | 149.18 | 143.15 | 146.91 | 259.09 | 254.54 | 199.04 |
| **EFR3A** | 410.17 | 403.20 | 409.10 | 708.40 | 706.40 | 764.76 |
| **EGFR** | 96.47 | 104.84 | 105.43 | 197.73 | 225.15 | 181.52 |
| **EPHA4** | 1931.13 | 2139.97 | 2007.43 | 1246.78 | 1242.76 | 1307.95 |
| **EPHA5** | 175.97 | 189.59 | 179.50 | 734.13 | 682.52 | 584.53 |
| **EPHA6** | 93.01 | 116.45 | 126.17 | 239.29 | 239.84 | 184.02 |
| **ERLEC1** | 1751.38 | 1732.48 | 1730.43 | 2614.50 | 2673.66 | 2654.67 |
| **FGF12** | 550.16 | 486.79 | 561.67 | 1485.29 | 1434.71 | 1568.28 |
| **FOS** | 500.91 | 539.03 | 546.86 | 983.53 | 928.66 | 1060.14 |
| **FRMPD4** | 177.70 | 179.14 | 189.86 | 267.99 | 312.40 | 259.12 |
| **GAA** | 240.79 | 261.57 | 260.97 | 361.02 | 405.16 | 436.84 |
| **GABRA1** | 234.74 | 268.53 | 267.63 | 590.63 | 591.60 | 549.49 |
| **GABRA4** | 144.86 | 152.44 | 180.24 | 483.74 | 558.53 | 544.48 |
| **GABRB2** | 831.02 | 912.86 | 988.29 | 1643.64 | 1734.11 | 1525.73 |
| **GABRG2** | 361.77 | 346.32 | 359.48 | 799.45 | 797.32 | 652.12 |
| **GAD1** | 3793.44 | 4288.86 | 4409.38 | 10285.44 | 10289.23 | 11766.26 |
| **GAL3ST1** | 10.91 | 9.65 | 19.51 | 47.30 | 39.62 | 31.33 |
| **GBA** | 246.84 | 232.55 | 260.23 | 564.90 | 636.60 | 516.95 |
| **GDPD2** | 136.22 | 167.53 | 140.24 | 677.72 | 690.79 | 579.53 |
| **GLRB** | 428.31 | 458.93 | 423.91 | 971.65 | 1092.14 | 929.97 |
| **GLS** | 1412.62 | 1553.69 | 1632.66 | 2414.59 | 2432.12 | 2381.82 |
| **GNAO1** | 2888.64 | 2923.60 | 3047.32 | 4846.21 | 5040.44 | 4461.97 |
| **GNPTAB** | 1260.52 | 1271.59 | 1321.59 | 1874.23 | 1987.60 | 1986.32 |
| **GNPTG** | 410.17 | 350.96 | 346.88 | 759.86 | 800.08 | 732.22 |
| **GPD1L** | 290.91 | 277.82 | 331.33 | 646.05 | 684.36 | 602.05 |
| **GRIA1** | 1471.38 | 1467.78 | 1507.49 | 2608.57 | 2769.18 | 2849.91 |
| **GRIA2** | 2817.78 | 3314.84 | 3131.01 | 5621.12 | 5630.98 | 7002.70 |
| **GRIA4** | 2086.68 | 2245.61 | 2225.93 | 3307.27 | 3422.18 | 3498.24 |
| **GRIK2** | 520.78 | 532.07 | 590.56 | 897.42 | 866.20 | 889.92 |
| **GRIN1** | 86.10 | 100.20 | 63.21 | 270.96 | 266.47 | 246.60 |
| **GRIN2B** | 897.57 | 990.64 | 1041.62 | 1718.85 | 1725.85 | 1778.55 |
| **GRM5** | 446.46 | 533.23 | 526.86 | 1282.41 | 1285.01 | 1232.86 |
| **GRN** | 460.29 | 375.34 | 345.40 | 743.04 | 766.10 | 734.72 |
| **GUCY1B3** | 281.40 | 244.16 | 264.67 | 672.77 | 637.52 | 469.38 |
| **HAP1** | 290.05 | 313.81 | 318.74 | 495.62 | 561.29 | 589.54 |
| **HCN1** | 55.85 | 54.92 | 65.43 | 262.06 | 267.39 | 246.60 |
| **HDAC1** | 966.70 | 908.21 | 930.52 | 596.56 | 549.35 | 567.01 |
| **HEXB** | 408.44 | 395.08 | 406.88 | 683.66 | 731.20 | 649.61 |
| **HMOX1** | 23.88 | 13.13 | 9.14 | 90.84 | 63.50 | 101.42 |
| **HSPB1** | 305.60 | 249.96 | 260.23 | 610.42 | 523.63 | 559.50 |
| **IL13RA1** | 78.32 | 47.96 | 58.77 | 91.83 | 95.65 | 96.41 |
| **INHBB** | 10.05 | 6.16 | 13.59 | 55.21 | 62.58 | 31.33 |
| **INPP5F** | 2608.65 | 2577.64 | 2563.67 | 4177.19 | 4362.64 | 4251.70 |
| **ITPR1** | 29.06 | 40.99 | 41.73 | 114.59 | 124.12 | 61.37 |
| **ITPR2** | 20.42 | 17.77 | 18.77 | 216.53 | 215.96 | 229.08 |
| **LAMB2** | 103.38 | 96.72 | 110.61 | 208.61 | 204.02 | 153.98 |

| | | | | | | |
|---|---|---|---|---|---|---|
| **LAMP1** | 1212.13 | 1102.09 | 1031.99 | 1675.31 | 1888.41 | 1871.17 |
| **LIF** | 28.20 | 29.38 | 22.48 | 10.68 | 19.42 | 2.50 |
| **LPAR1** | 93.88 | 75.82 | 66.92 | 257.11 | 226.98 | 138.96 |
| **MAGEE1** | 652.14 | 662.09 | 673.51 | 1156.72 | 1185.82 | 1190.30 |
| **MAP2K1** | 655.59 | 672.54 | 652.77 | 1194.33 | 1261.13 | 1117.71 |
| **MAPT** | 2078.90 | 1942.61 | 2055.58 | 3575.47 | 3445.14 | 3355.56 |
| **MYC** | 226.96 | 222.10 | 196.53 | 374.88 | 355.56 | 379.27 |
| **NEFH** | 61.04 | 57.24 | 66.92 | 163.09 | 172.80 | 158.99 |
| **NEFL** | 12334.15 | 11433.28 | 11307.85 | 21757.71 | 23144.39 | 22542.47 |
| **NES** | 6665.98 | 6645.57 | 5963.28 | 3727.88 | 3473.61 | 3308.00 |
| **NGF** | 4.86 | 6.16 | 3.22 | 1.77 | 1.00 | 2.50 |
| **NLGN4X** | 1399.66 | 1459.66 | 1403.80 | 2152.33 | 2073.93 | 2281.69 |
| **NMB** | 169.06 | 144.31 | 156.54 | 306.59 | 302.29 | 244.10 |
| **NOL3** | 105.11 | 101.36 | 99.50 | 186.84 | 207.70 | 219.07 |
| **NOTCH3** | 528.56 | 455.45 | 432.06 | 352.12 | 322.50 | 264.12 |
| **NPAS4** | 25.61 | 21.25 | 29.88 | 118.55 | 156.26 | 96.41 |
| **NPC2** | 799.91 | 706.21 | 767.58 | 1584.26 | 1511.86 | 1598.32 |
| **NPTN** | 2174.83 | 2156.22 | 2248.89 | 3537.87 | 3698.62 | 3668.46 |
| **NPY** | 2.27 | 7.32 | 3.96 | 808.35 | 710.07 | 409.31 |
| **NQO1** | 112.89 | 117.61 | 98.02 | 448.11 | 437.30 | 441.85 |
| **NRXN1** | 780.90 | 696.92 | 656.48 | 1506.07 | 1436.55 | 2016.35 |
| **NSF** | 1613.97 | 1511.90 | 1502.31 | 2323.54 | 2430.28 | 2306.72 |
| **NTNG1** | 104.25 | 172.18 | 159.50 | 438.22 | 395.97 | 554.49 |
| **NTRK1** | 773.12 | 763.09 | 676.48 | 225.44 | 193.00 | 133.96 |
| **NTS** | 739.42 | 737.55 | 886.08 | 2647.16 | 3122.77 | 3060.18 |
| **OLFM3** | 468.93 | 454.28 | 445.39 | 1006.29 | 992.95 | 827.34 |
| **OLIG2** | 383.38 | 400.88 | 338.74 | 240.28 | 210.45 | 161.49 |
| **OPTN** | 1138.67 | 1158.97 | 1229.01 | 1926.68 | 1990.35 | 2001.33 |
| **PCNA** | 548.44 | 471.70 | 439.47 | 248.20 | 248.11 | 234.09 |
| **PDGFRB** | 26.47 | 25.90 | 18.03 | 322.43 | 271.07 | 174.01 |
| **PIK3CB** | 120.67 | 94.39 | 101.73 | 250.18 | 236.17 | 204.05 |
| **PINK1** | 788.68 | 757.29 | 763.13 | 1417.00 | 1386.95 | 1207.82 |
| **PLA2G16** | 192.39 | 190.75 | 203.20 | 739.08 | 775.28 | 722.21 |
| **PLA2G4C** | 137.08 | 159.41 | 170.61 | 306.59 | 317.91 | 254.11 |
| **PLCB1** | 252.02 | 299.88 | 291.33 | 582.71 | 597.11 | 511.94 |
| **PLXNB3** | 15.24 | 28.22 | 18.03 | 82.93 | 67.18 | 63.87 |
| **PMP22** | 76.59 | 46.80 | 40.25 | 264.04 | 239.84 | 151.48 |
| **PPARGC1A** | 297.82 | 316.13 | 335.77 | 587.66 | 651.29 | 654.62 |
| **PPT1** | 1057.44 | 923.30 | 994.96 | 1733.70 | 1743.30 | 1718.47 |
| **PRKCA** | 1229.41 | 1301.77 | 1363.06 | 2106.80 | 2136.38 | 2048.90 |
| **PRKCE** | 199.31 | 207.00 | 193.57 | 373.89 | 362.91 | 286.65 |
| **PRNP** | 2351.12 | 2102.82 | 2285.18 | 4677.97 | 4904.51 | 4619.67 |
| **PTDSS1** | 1184.47 | 1104.41 | 1123.09 | 1726.77 | 1786.46 | 1640.88 |
| **PTPRR** | 77.46 | 88.59 | 93.58 | 269.97 | 271.99 | 296.67 |
| **RIMS1** | 410.17 | 362.57 | 377.99 | 825.18 | 816.61 | 804.81 |
| **RTN4** | 6812.89 | 6659.50 | 6788.37 | 10387.38 | 10513.32 | 10061.59 |
| **RYR1** | 46.35 | 92.07 | 85.43 | 137.36 | 124.12 | 171.51 |
| **RYR2** | 164.74 | 253.44 | 208.38 | 583.70 | 557.61 | 554.49 |
| **RYR3** | 245.11 | 254.60 | 283.93 | 423.37 | 449.24 | 409.31 |
| **S100B** | 58.44 | 37.51 | 46.18 | 1005.30 | 857.02 | 514.44 |

| | | | | | | |
|---|---|---|---|---|---|---|
| **SCN1A** | 473.25 | 520.46 | 485.39 | 1278.45 | 1296.95 | 1355.51 |
| **SCN2A** | 1780.76 | 1879.92 | 1914.11 | 3230.08 | 3262.37 | 3202.86 |
| **SHH** | 608.93 | 664.41 | 607.59 | 305.60 | 268.31 | 404.30 |
| **SIRT2** | 1148.18 | 1088.16 | 1147.53 | 1915.80 | 1950.86 | 1861.16 |
| **SLC1A1** | 594.24 | 573.86 | 594.26 | 1019.15 | 1059.99 | 1087.67 |
| **SLC2A1** | 240.79 | 244.16 | 228.38 | 126.47 | 128.71 | 133.96 |
| **SLC32A1** | 341.90 | 280.14 | 326.89 | 654.96 | 623.74 | 737.23 |
| **SLC4A10** | 313.38 | 419.46 | 368.36 | 557.97 | 584.25 | 587.03 |
| **SLC9A6** | 801.64 | 799.08 | 824.61 | 1267.56 | 1347.46 | 1260.39 |
| **SNAP91** | 1585.45 | 1695.33 | 1611.18 | 2724.36 | 2748.97 | 3092.72 |
| **SNCA** | 544.98 | 551.80 | 506.87 | 1044.89 | 1180.31 | 1127.72 |
| **SNCB** | 332.39 | 337.03 | 331.33 | 940.97 | 946.11 | 909.95 |
| **SORCS3** | 210.54 | 245.32 | 210.60 | 377.85 | 405.16 | 461.88 |
| **SORL1** | 197.58 | 207.00 | 231.34 | 500.57 | 499.75 | 511.94 |
| **SP1** | 184.61 | 182.63 | 173.57 | 131.42 | 121.36 | 93.91 |
| **STX1A** | 636.58 | 628.43 | 649.07 | 976.60 | 998.46 | 1087.67 |
| **SYT1** | 6142.29 | 5704.05 | 5707.01 | 9777.74 | 9523.26 | 9578.48 |
| **SYT13** | 1792.86 | 1756.86 | 1775.61 | 3231.07 | 3175.12 | 3130.27 |
| **SYT4** | 1582.00 | 1482.88 | 1452.68 | 3110.33 | 3210.02 | 3115.25 |
| **SYT7** | 520.78 | 523.94 | 575.75 | 814.29 | 849.67 | 782.28 |
| **TBR1** | 144.00 | 155.92 | 166.90 | 54.23 | 61.67 | 36.33 |
| **TGFB1** | 84.37 | 93.23 | 89.14 | 167.05 | 215.04 | 148.98 |
| **TH** | 277.08 | 309.17 | 323.18 | 563.91 | 621.90 | 572.02 |
| **THY1** | 1422.12 | 1289.00 | 1400.84 | 3422.08 | 3560.86 | 3458.19 |
| **TLR4** | 3.14 | 1.16 | 1.00 | 64.12 | 67.18 | 51.35 |
| **TNC** | 90.42 | 103.68 | 94.32 | 578.75 | 612.72 | 662.13 |
| **TNFRSF10B** | 503.50 | 570.38 | 500.20 | 326.38 | 348.21 | 304.17 |
| **TNFRSF1A** | 179.43 | 169.85 | 146.17 | 281.85 | 298.62 | 264.12 |
| **TNR** | 455.97 | 442.68 | 479.46 | 1096.35 | 1130.71 | 1092.68 |
| **TP53** | 923.49 | 932.59 | 906.08 | 400.61 | 416.18 | 321.70 |
| **TSPO** | 1.00 | 1.16 | 1.00 | 2.76 | 2.89 | 2.50 |
| **UNC13A** | 1225.09 | 1403.93 | 1460.83 | 1966.27 | 2051.89 | 2176.56 |
| **WFS1** | 114.62 | 111.81 | 113.58 | 204.66 | 228.82 | 229.08 |
| **XK** | 115.48 | 94.39 | 117.28 | 206.63 | 183.82 | 169.00 |

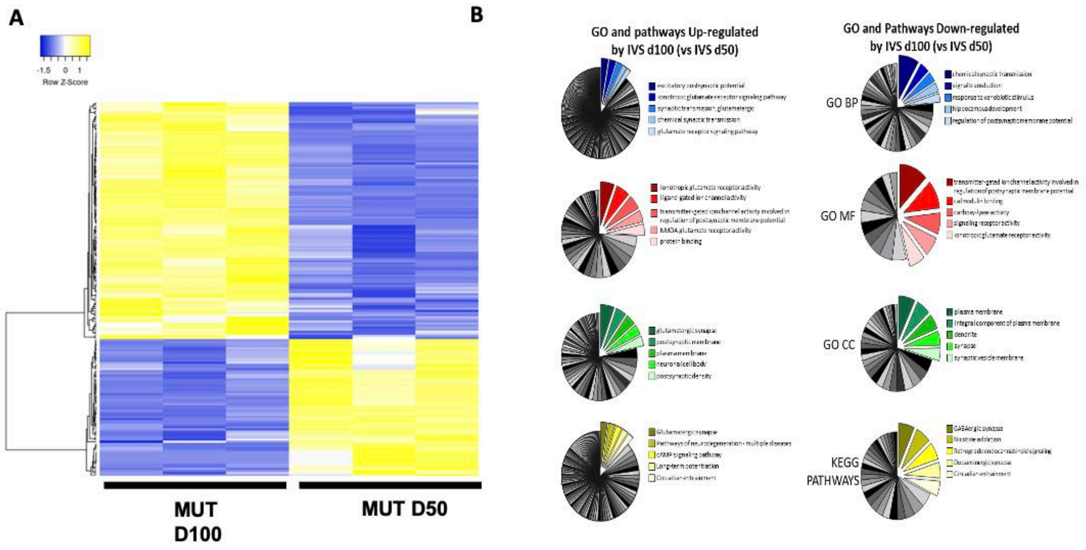


**Figure S4**. **A)** Heat map of supervised hierarchical clustering of 180 differentially expressed (DE) genes between D50 MUT and D100 MUT (114 genes up- and 66 genes down-regulated in D100 MUT). Average linkege and Pearson Distance Measurement Methods was performed for Hierarchical cluster using Heatmapper tools (http://www.heatmapper.ca) . The significant DE genes are selected by multiple t-test performed with Graph Pad Prism 6, with *pvalue* ≤ 0.05 and a fold change cut-off values of ≥1.5 or ≤0.66 for upmodulated and downregulated genes respectively. **B)** Gene Ontology (GO) and KEGG Pathways enrichment analysis of 180 down- and up-regulated genes by D100 MUT (respect to D50 MUT) have been analyzed by DAVID database. An overview of top 5 significantly enriched terms in KEGG pathways and three GO categories: biological process (BP), cellular component (CC), and molecular function (MF). The cut-off of p-value was set to 0.05 and are selected only enriched terms in which are represented a minimum of 3 genes. The −log10 p-value associated with each category is represented by the size of the pie slice.

Supplementary Figure 5

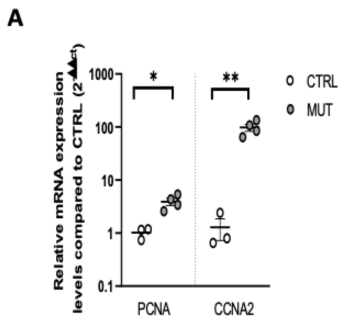


Figure S5. **A)** Dot plot of Real-time PCR analysis of PCNA and CCNA2 on control (CTRL) and tau-mutant (MUT) derived cortical organoids at D100. (CTRL n= 3/2, MUT n=4/2; replicates/batches). Gene expression is normalized to the housekeeping gene ATP5O ( *p<0.05, MW test; ** p<0.001 MW test).

Supplementary Figure 6

Supplementary Table 7

| Probe Name | IVS d50 _1 | IVS d50 _2 | IVS d50 _3 | IVS d100 _1 | IVS d100 _2 | IVS d100 _3 |
|---|---|---|---|---|---|---|
| ACAA1 | 531.82 | 431.67 | 430.42 | 787.08 | 773.70 | 847.09 |
| ACHE | 695.36 | 730.74 | 684.34 | 350.59 | 303.97 | 488.95 |
| ACIN1 | 2713.22 | 2282.48 | 2581.41 | 4091.90 | 3976.09 | 4462.21 |
| ACTN1 | 546.04 | 514.02 | 523.80 | 844.56 | 708.91 | 847.09 |
| ADORA1 | 135.08 | 143.43 | 193.47 | 224.91 | 275.39 | 265.12 |
| ADORA2A | 31.27 | 15.56 | 49.15 | 85.58 | 85.78 | 94.50 |
| ADRA2A | 103.79 | 19.90 | 59.95 | 358.39 | 644.12 | 879.18 |
| ALDH1L1 | 1.42 | 2.17 | 2.07 | 15.43 | 18.13 | 8.34 |
| AMPH | 3439.88 | 2848.12 | 3261.37 | 1470.06 | 1265.35 | 1566.73 |
| AP3M2 | 918.62 | 700.40 | 869.57 | 1269.36 | 1302.51 | 1254.21 |
| APC | 1645.27 | 1190.20 | 1453.82 | 2193.96 | 2240.07 | 2365.77 |
| AQP4 | 1.42 | 2.17 | 1.00 | 10.56 | 5.74 | 4.97 |
| ATP6V0E1 | 1397.84 | 1144.68 | 1199.13 | 2150.12 | 1911.36 | 2152.08 |
| BAX | 1577.02 | 1565.12 | 1301.77 | 2385.90 | 2127.64 | 2199.38 |
| BCHE | 126.54 | 214.95 | 175.72 | 613.65 | 762.27 | 529.50 |
| BID | 167.78 | 165.10 | 146.39 | 411.97 | 423.07 | 459.39 |
| C3 | 1.42 | 2.17 | 2.07 | 1.00 | 1.00 | 1.00 |
| CACNA1C | 1719.22 | 1196.70 | 1564.19 | 723.75 | 670.80 | 968.72 |
| CACNB2 | 337.00 | 323.31 | 421.15 | 208.34 | 162.00 | 287.92 |
| CADPS | 1945.32 | 1242.21 | 1574.99 | 895.22 | 793.71 | 1002.50 |
| CALB2 | 261.64 | 290.80 | 269.88 | 1107.62 | 863.27 | 505.85 |
| CALM1 | 3486.81 | 3194.88 | 3096.98 | 4470.90 | 5132.80 | 5099.92 |
| CASP6 | 423.75 | 364.48 | 376.39 | 804.61 | 867.08 | 772.76 |
| CASP7 | 96.68 | 63.24 | 73.84 | 198.60 | 157.24 | 134.20 |
| CAST | 472.10 | 329.81 | 404.17 | 255.11 | 181.06 | 293.84 |
| CCND1 | 1474.63 | 1456.76 | 1273.99 | 282.39 | 121.03 | 285.39 |
| CDK5 | 453.61 | 429.50 | 417.29 | 656.52 | 646.03 | 694.20 |
| CLDN15 | 2.83 | 24.23 | 7.47 | 42.71 | 36.23 | 27.77 |
| CLN3 | 76.77 | 54.57 | 68.44 | 107.99 | 113.41 | 118.15 |
| CLU | 5213.15 | 3279.40 | 3802.40 | 8411.94 | 5885.52 | 9574.90 |
| CNKSR2 | 1594.08 | 1296.39 | 1446.10 | 2877.92 | 2923.24 | 2627.62 |
| CNTF | 126.54 | 93.58 | 137.13 | 215.16 | 249.66 | 238.93 |
| CRH | 592.97 | 557.37 | 525.35 | 49.53 | 53.38 | 141.80 |
| CRTC2 | 226.09 | 178.10 | 223.57 | 318.44 | 312.54 | 310.73 |
| CTNS | 28.42 | 6.89 | 27.54 | 60.25 | 47.66 | 70.85 |
| CUL1 | 1962.39 | 1955.22 | 1955.49 | 2995.81 | 3038.53 | 2783.88 |
| CUL3 | 1595.50 | 1532.62 | 1443.79 | 2236.83 | 2461.13 | 2235.70 |
| CXXC1 | 132.23 | 93.58 | 126.33 | 215.16 | 204.88 | 221.20 |
| CYP4X1 | 464.99 | 238.79 | 382.56 | 85.58 | 61.00 | 150.25 |
| DAGLA | 324.21 | 282.13 | 354.01 | 474.33 | 471.66 | 510.91 |
| DDC | 17.05 | 19.90 | 25.99 | 1.00 | 4.79 | 1.00 |
| DDIT3 | 1124.81 | 1580.30 | 1243.12 | 778.31 | 572.66 | 925.64 |
| DDX23 | 568.80 | 581.21 | 531.52 | 873.79 | 835.64 | 874.11 |
| DGKB | 351.22 | 178.10 | 252.90 | 555.19 | 470.71 | 418.85 |
| DLGAP1 | 551.73 | 579.04 | 658.87 | 1927.98 | 2297.24 | 2162.21 |
| DLX1 | 1080.73 | 1504.44 | 1239.26 | 420.74 | 139.13 | 84.36 |
| DLX2 | 543.20 | 973.47 | 746.85 | 203.47 | 87.68 | 34.53 |
| DRD4 | 73.93 | 41.57 | 70.76 | 34.92 | 23.84 | 25.24 |
| EFNA1 | 34.11 | 17.73 | 43.74 | 68.04 | 101.97 | 128.28 |

| | | | | | | |
|---|---|---|---|---|---|---|
| EGF | 39.80 | 37.23 | 56.09 | 79.74 | 111.50 | 84.36 |
| EGR1 | 18.47 | 13.39 | 40.66 | 57.33 | 68.63 | 108.01 |
| EPHA3 | 3512.40 | 2360.50 | 3264.46 | 580.53 | 581.24 | 664.64 |
| EPHA5 | 274.43 | 253.96 | 270.65 | 1443.75 | 1858.95 | 1322.63 |
| EPHA7 | 861.73 | 804.43 | 935.94 | 3392.35 | 3918.92 | 3207.05 |
| FA2H | 1.42 | 2.17 | 2.07 | 1.00 | 1.00 | 1.00 |
| FAM126A | 573.06 | 479.35 | 481.35 | 1067.68 | 1162.45 | 1331.92 |
| FGF12 | 413.79 | 297.30 | 369.44 | 531.81 | 549.79 | 751.64 |
| FGF14 | 2791.43 | 1697.33 | 2325.18 | 456.79 | 263.95 | 606.36 |
| GABRA1 | 129.39 | 119.59 | 154.11 | 60.25 | 35.28 | 61.56 |
| GABRB2 | 725.22 | 676.56 | 706.72 | 296.03 | 289.68 | 390.97 |
| GAD1 | 1353.76 | 1380.91 | 1370.46 | 436.33 | 132.46 | 336.91 |
| GAD2 | 432.28 | 581.21 | 522.26 | 95.33 | 25.75 | 33.68 |
| GALC | 543.20 | 412.16 | 471.32 | 253.16 | 278.24 | 249.07 |
| GBA | 351.22 | 217.11 | 314.65 | 549.35 | 483.10 | 634.23 |
| GDPD2 | 446.50 | 214.95 | 330.85 | 685.75 | 628.88 | 710.25 |
| GLRB | 614.30 | 561.70 | 620.28 | 1435.96 | 1599.79 | 1523.65 |
| GNAI1 | 1386.46 | 1181.53 | 1294.06 | 1891.93 | 2240.07 | 2277.09 |
| GNGT1 | 1.42 | 2.17 | 2.84 | 1.00 | 1.00 | 1.00 |
| GPD1L | 305.72 | 344.98 | 314.65 | 899.12 | 1050.97 | 1041.36 |
| GRIA1 | 1548.58 | 1378.74 | 1491.64 | 2941.25 | 4038.98 | 3035.58 |
| GRIA2 | 10043.80 | 9713.88 | 11036.46 | 5869.02 | 5573.95 | 6001.17 |
| GRIA3 | 1035.22 | 839.11 | 953.69 | 5194.81 | 6316.19 | 6420.96 |
| GRIA4 | 837.56 | 622.38 | 818.63 | 424.64 | 449.75 | 489.80 |
| GRIK2 | 699.62 | 503.19 | 682.79 | 910.81 | 1147.20 | 1067.54 |
| GRIN1 | 32.69 | 41.57 | 64.58 | 188.86 | 172.48 | 204.30 |
| GRIN2A | 39.80 | 32.90 | 53.01 | 105.07 | 85.78 | 79.29 |
| GRIN2B | 1130.50 | 1008.15 | 1193.72 | 1739.94 | 2213.40 | 2027.91 |
| GRIN2C | 1.42 | 2.17 | 6.70 | 47.58 | 47.66 | 63.25 |
| GRIN2D | 514.76 | 592.04 | 557.76 | 206.40 | 198.21 | 241.47 |
| GRIN3B | 4.25 | 2.17 | 1.00 | 7.64 | 7.65 | 5.81 |
| HGF | 1.42 | 2.17 | 5.15 | 81.69 | 88.64 | 68.31 |
| HIF1A | 2526.93 | 2256.47 | 2239.51 | 3784.02 | 3835.07 | 3627.69 |
| HNRNPM | 5281.41 | 4161.46 | 4463.06 | 8462.60 | 8759.18 | 7589.96 |
| HSPB1 | 292.92 | 169.43 | 195.02 | 493.81 | 478.33 | 494.86 |
| IDE | 261.64 | 243.12 | 313.10 | 512.33 | 477.38 | 526.12 |
| IDH1 | 1511.60 | 1333.23 | 1383.58 | 2374.21 | 2599.28 | 2459.53 |
| IGF1R | 2170.00 | 1849.03 | 2010.28 | 1249.87 | 1255.82 | 1174.81 |
| IKBKB | 253.10 | 236.62 | 238.24 | 482.12 | 474.52 | 442.50 |
| INHBB | 34.11 | 28.56 | 46.83 | 179.11 | 153.43 | 121.53 |
| ITPR2 | 36.95 | 2.17 | 33.71 | 153.78 | 100.07 | 112.24 |
| KATNA1 | 501.96 | 511.86 | 513.77 | 834.82 | 813.72 | 744.04 |
| KCNB1 | 437.97 | 412.16 | 424.24 | 232.70 | 234.41 | 270.19 |
| KCNJ10 | 1.42 | 2.17 | 4.38 | 74.86 | 61.00 | 66.63 |
| KIAA1161 | 534.67 | 546.53 | 540.01 | 880.61 | 1077.65 | 868.20 |
| LAMB2 | 184.85 | 110.92 | 162.60 | 319.41 | 267.76 | 332.69 |
| LAMP1 | 1130.50 | 841.27 | 927.45 | 1541.18 | 1532.14 | 1542.24 |
| LIF | 4.25 | 6.89 | 9.79 | 43.69 | 21.94 | 52.27 |
| LRRC25 | 1.42 | 2.17 | 2.07 | 1.00 | 1.00 | 1.00 |
| MAPK10 | 3569.28 | 2655.24 | 3090.80 | 1827.63 | 1837.99 | 2271.17 |

| | | | | | | |
|---|---|---|---|---|---|---|
| **MBP** | 38.38 | 43.74 | 49.92 | 7.64 | 1.00 | 19.32 |
| **MGMT** | 69.66 | 37.23 | 62.27 | 34.92 | 24.80 | 16.79 |
| **MSN** | 516.18 | 475.01 | 481.35 | 1072.55 | 1079.56 | 918.04 |
| **MTA2** | 1743.39 | 1842.53 | 1696.16 | 2946.12 | 3104.27 | 2697.72 |
| **MTOR** | 564.53 | 457.67 | 547.73 | 773.44 | 811.82 | 798.10 |
| **MYD88** | 1.42 | 2.17 | 15.96 | 41.74 | 37.18 | 38.75 |
| **NAGLU** | 92.41 | 76.24 | 86.19 | 154.76 | 138.18 | 172.21 |
| **NEFH** | 1.42 | 2.17 | 19.05 | 28.10 | 24.80 | 27.77 |
| **NEFL** | 5437.84 | 5942.91 | 5513.47 | 1932.85 | 1332.05 | 2682.52 |
| **NELL2** | 2522.67 | 1597.63 | 1842.80 | 9024.77 | 10758.17 | 10236.26 |
| **NGFR** | 103.79 | 76.24 | 97.00 | 13.48 | 18.13 | 29.46 |
| **NLGN4X** | 2652.07 | 2488.36 | 2589.90 | 1752.61 | 1583.59 | 1548.99 |
| **NMB** | 152.14 | 95.75 | 125.56 | 236.60 | 256.33 | 265.12 |
| **NOSTRIN** | 1.42 | 2.17 | 5.15 | 20.30 | 19.08 | 15.10 |
| **NOVA1** | 5096.55 | 3816.87 | 4672.99 | 1846.14 | 1668.39 | 1721.30 |
| **NPAS4** | 1.42 | 2.17 | 21.36 | 56.35 | 40.99 | 104.63 |
| **NPY** | 8.51 | 11.23 | 19.05 | 142.09 | 171.53 | 190.79 |
| **NTNG1** | 1176.00 | 787.09 | 1072.55 | 334.03 | 227.75 | 451.79 |
| **NTS** | 342.69 | 442.50 | 379.48 | 2246.58 | 1618.84 | 2675.76 |
| **P2RX4** | 5.67 | 4.73 | 20.59 | 23.23 | 28.61 | 26.93 |
| **PARP1** | 2625.05 | 2304.15 | 2256.49 | 3691.46 | 3710.26 | 3384.43 |
| **PCSK2** | 2472.90 | 1768.84 | 2308.20 | 453.87 | 302.06 | 600.45 |
| **PDE1B** | 156.41 | 126.09 | 167.23 | 30.05 | 4.79 | 54.80 |
| **PDE4D** | 312.83 | 210.61 | 295.35 | 402.23 | 516.45 | 554.84 |
| **PDGFRB** | 180.58 | 104.42 | 146.39 | 515.25 | 350.66 | 554.84 |
| **PECAM1** | 1.42 | 2.17 | 2.07 | 1.00 | 1.00 | 1.00 |
| **PHF19** | 36.95 | 56.74 | 62.27 | 145.01 | 162.95 | 116.46 |
| **PIK3CB** | 152.14 | 104.42 | 174.18 | 317.47 | 340.18 | 284.55 |
| **PLCB1** | 574.48 | 433.84 | 519.94 | 892.30 | 953.78 | 808.23 |
| **PLCB3** | 152.14 | 132.59 | 146.39 | 230.75 | 247.75 | 236.40 |
| **PLXNC1** | 1666.60 | 1688.66 | 1710.83 | 602.93 | 588.86 | 522.74 |
| **PMP22** | 68.24 | 48.07 | 62.27 | 159.63 | 209.64 | 199.24 |
| **POLR2B** | 2235.42 | 2113.43 | 2158.47 | 3394.30 | 3531.13 | 3228.17 |
| **POLR2J** | 119.43 | 139.09 | 153.34 | 188.86 | 218.22 | 216.13 |
| **PPM1L** | 628.52 | 587.71 | 614.10 | 889.38 | 1139.58 | 991.52 |
| **PRKCA** | 1046.60 | 1079.67 | 1135.07 | 2464.82 | 2360.13 | 2303.27 |
| **PRKCB** | 544.62 | 509.69 | 537.69 | 221.01 | 414.50 | 335.23 |
| **PSMB8** | 1.42 | 2.17 | 1.00 | 9.59 | 14.32 | 11.72 |
| **PTPRN2** | 2403.22 | 1944.39 | 2001.79 | 963.43 | 900.43 | 1059.09 |
| **PTPRR** | 184.85 | 152.10 | 174.95 | 34.92 | 20.03 | 34.53 |
| **RAB3C** | 2707.53 | 2273.81 | 2308.20 | 722.77 | 561.23 | 650.28 |
| **RAF1** | 1844.36 | 1606.30 | 1710.83 | 2637.27 | 2832.72 | 2678.30 |
| **RAPGEF2** | 995.41 | 912.79 | 928.22 | 1536.31 | 1564.53 | 1556.59 |
| **RASGRP1** | 947.06 | 615.88 | 801.65 | 258.03 | 269.67 | 349.58 |
| **RET** | 179.16 | 71.91 | 141.76 | 32.00 | 4.79 | 43.82 |
| **RYR1** | 95.26 | 95.75 | 130.19 | 49.53 | 34.33 | 59.02 |
| **RYR2** | 275.86 | 260.46 | 273.74 | 829.95 | 766.08 | 755.86 |
| **RYR3** | 226.09 | 156.43 | 217.40 | 83.63 | 52.43 | 107.17 |
| **SCAMP2** | 474.94 | 338.48 | 354.78 | 646.78 | 625.07 | 633.39 |
| **SF3A2** | 1110.59 | 1465.43 | 1235.40 | 822.15 | 778.47 | 852.15 |

| | | | | | | |
|---|---|---|---|---|---|---|
| **SIRT7** | 349.80 | 312.47 | 295.35 | 578.58 | 532.64 | 688.29 |
| **SLA** | 32.69 | 24.23 | 47.60 | 897.17 | 1312.04 | 1253.36 |
| **SLC12A5** | 698.20 | 698.24 | 657.32 | 155.73 | 142.95 | 196.70 |
| **SLC17A6** | 3018.96 | 2362.66 | 2693.33 | 451.92 | 290.63 | 744.88 |
| **SLC18A3** | 1.42 | 2.17 | 2.07 | 1.00 | 1.00 | 1.00 |
| **SLC32A1** | 115.17 | 152.10 | 133.27 | 21.28 | 1.00 | 3.28 |
| **SLC8A1** | 4715.44 | 4040.09 | 4384.33 | 2752.24 | 2854.64 | 2818.51 |
| **SNCA** | 683.98 | 449.01 | 509.14 | 1382.37 | 1493.07 | 1411.31 |
| **SNCAIP** | 335.58 | 388.32 | 360.18 | 633.14 | 664.13 | 492.33 |
| **SORCS3** | 248.84 | 201.94 | 258.30 | 69.02 | 59.10 | 121.53 |
| **SOX9** | 2828.41 | 2616.23 | 2515.81 | 4506.95 | 4188.57 | 3614.17 |
| **STAMBPL1** | 1542.89 | 1159.85 | 1358.89 | 310.65 | 296.35 | 416.31 |
| **STX1A** | 557.42 | 509.69 | 523.80 | 836.77 | 868.98 | 1046.42 |
| **SYT1** | 4881.82 | 3327.08 | 4087.19 | 5869.99 | 7145.13 | 6863.56 |
| **SYT13** | 2583.82 | 2670.41 | 2492.66 | 1305.40 | 1548.34 | 1702.72 |
| **SYT4** | 1975.19 | 1627.97 | 1754.82 | 3294.92 | 3464.43 | 4843.99 |
| **TAF6L** | 174.89 | 132.59 | 157.20 | 261.93 | 255.38 | 272.72 |
| **TCIRG1** | 2.83 | 2.17 | 27.54 | 36.87 | 47.66 | 60.71 |
| **TENM2** | 581.59 | 305.97 | 522.26 | 94.35 | 55.29 | 103.79 |
| **TF** | 39.80 | 24.23 | 42.20 | 19.33 | 13.36 | 13.41 |
| **TGFB1** | 51.18 | 115.25 | 86.97 | 221.98 | 208.69 | 139.27 |
| **TH** | 110.90 | 212.78 | 189.61 | 27.12 | 10.50 | 45.51 |
| **TNC** | 244.57 | 321.14 | 282.23 | 741.29 | 556.46 | 684.07 |
| **TP53** | 1379.35 | 1315.89 | 1246.21 | 818.25 | 822.30 | 661.26 |
| **TRIM37** | 826.18 | 776.26 | 721.38 | 1278.12 | 1301.56 | 1300.66 |
| **TSPO** | 19.89 | 28.56 | 27.54 | 62.20 | 100.07 | 68.31 |
| **UBE2N** | 1.42 | 2.17 | 2.07 | 1.00 | 1.00 | 1.00 |
| **UGT8** | 140.76 | 178.10 | 200.42 | 76.81 | 68.63 | 81.83 |
| **VCP** | 2064.77 | 1753.67 | 1909.18 | 3034.78 | 3165.25 | 3007.71 |
| **WFS1** | 133.65 | 74.08 | 102.40 | 198.60 | 162.95 | 197.55 |
| **XBP1** | 449.34 | 468.51 | 387.97 | 217.11 | 217.26 | 238.09 |

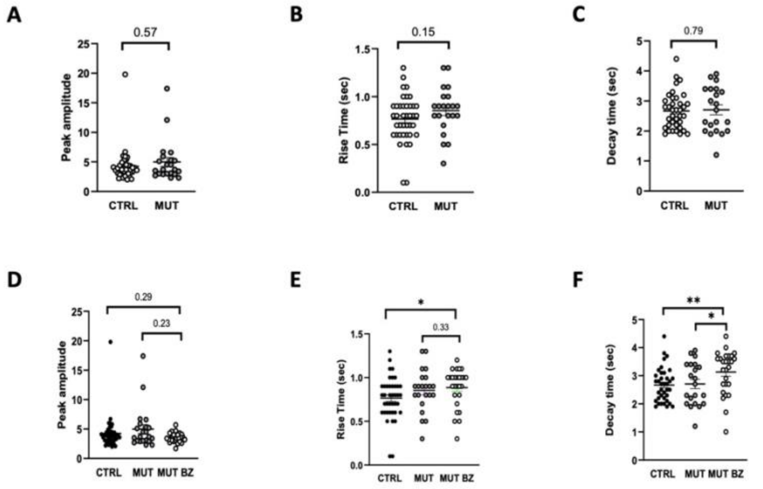


**Figure S6.**
**A)** Bar chart ( dot plots) representing the peak amplitude of spontaneous calcium oscillation within control and Tau mutant cortical organoids recorded within the field of view (FOV) (CTRL n= 47/….. FOVs/batches; MUT n=29/
**B)** Bar chart ( dot plots) representing the rise tim
e of the spontaneous calcium oscillation recorded within control and Tau mutant cortical organoids. (CTRL n= 47/….. FOVs/batches; MUT n=29/
**C)** Bar chart ( dot plots) representing the decay time of the spontaneous calcium oscillation recorded within control and Tau mutant cortical organoids (CTRL n= 47/….. FOVs/batches; MUT n=29/
**D)** Bar chart ( dot plots) representing the peak amplitude of spontaneous calcium oscillation within control and Tau mutant cortical organoids recorded within the field of view (FOV) (CTRL n= 47/….. FOVs/batches; MUT n=29/
**E)** Bar chart ( dot plots) representing the rise tim
e of the spontaneous calcium oscillation recorded within control and Tau mutant cortical organoids. (CTRL n= 47/….. FOVs/batches; MUT n=29/
**F)** Bar chart ( dot plots) representing the decay time of the spontaneous calcium oscillation recorded within control and Tau mutant cortical organoids (CTRL n= 47/….. FOVs/batches; MUT n=29/

Supplementary Movie1 – Ca activity CTRL

Supplementary Movie2 – Ca activity MUT

Supplementary Movie3– Ca activity MUT

Supplementary Movie4– Ca activity MUT BZ
